# Programmed mineralization of metal-oxides by de novo designed protein templates

**DOI:** 10.1101/2024.06.24.600095

**Authors:** Amijai Saragovi, Harley Pyles, Evelyn B. Huddy, Andrew J. Borst, Nikita Hanikel, Paul S. Kwon, Fátima A. Dávila-Hernández, Xinqi Li, Alexis Courbet, Ryan D. Kibler, Wenzhao Dai, Lukasz Goldschmidt, Dionne K. Vafeados, Aza Allen, Kenneth D. Carr, Asim K. Bera, Alex Kang, Evans Brackenbrough, Sakshi Yadav Schmid, Yuna Bae, Lance Stewart, Shuai Zhang, James J. De Yoreo, David Baker

**Affiliations:** Institute for Protein Design, University of Washington, Seattle, WA, 98105, USA; Department of Biochemistry, University of Washington, Seattle, WA, 98195, USA; Physical Sciences Division, Pacific Northwest National Laboratory, Richland, WA, USA; Department of Materials Science and Engineering, University of Washington, Seattle, WA, USA; Howard Hughes Medical Institute, University of Washington, Seattle, WA, 98105, USA; Centre de Biologie Structurale, Université de Montpellier, CNRS, INSERM. Montpellier, France; Department of Chemistry, University of Zürich, Winterthurerstrasse, 190, 8057, Zürich, Switzerland

## Abstract

We describe a coupled de novo design and screening approach to identify protein interfaces which guide the mineralization of specific metal-oxides, including technologically relevant materials that are not found in biological systems. We identify two interfaces promoting hematite growth in conditions typically yielding magnetite, and three interfaces promoting heterogeneous ZnO nucleation under conditions where traditional ZnO-binding peptides and control proteins are ineffective. Cryo-electron microscopy analysis of a designed nanocage with ZnO nucleating interfaces lining interior cavities revealed atomic density, likely corresponding to growing ZnO, directly adjacent to the designed interfaces. Our findings suggest that designed proteins can template mineralization through modes beyond electrostatics, likely by ordering ions and water molecules, and provide a route to protein-metal-oxide hybrid materials with tunable structure and function.

## Main Text

The design of protein–mineral hierarchical assemblies, where proteins program the formation of defined compounds and custom geometries at atomic resolution, has not yet been achieved. Designed protein templates can use charge complementarity to manipulate the crystallization of native inorganic materials, such as calcium carbonate, by selectively nucleating specific polymorphs (*1*) or regulating intermediate mineral species in solution (*2*). However, the vast majority of the universe of possible protein-inorganic hybrids, including technologically relevant metal oxides, are not explored by nature, and require a general design approach. Previous work has leveraged native proteins, most notably ferritin, to encapsulate minerals (*3–5*). Random peptide sequences, typically 8-12 residues, have been selected from phage display libraries with biopanning and shown to bind a variety of inorganic materials, including sequences that modify growth (in conditions which also allow homogeneous growth) and promote nucleation when densely displayed on protein assemblies (*6–11*). However, native proteins and synthetic peptides do not provide a mechanistic understanding of how the structure of biomolecular interfaces controls inorganic crystallization and therefore do not provide a path to develop fully defined structured hybrid materials.

We reasoned that structurally and chemically defined protein interfaces could bind metal oxides more strongly and specifically and thereby exert greater control over nucleation and growth. At the lengths required to present such interfaces (>50 residues), sequence space is far too large to sample randomly, necessitating a directed approach. De novo protein design enables the construction of molecular scaffolds with a wide variety of surface geometries (*12*, *13*), and the patterning of diverse sidechain chemistries on these surfaces. We set out to use de novo design to identify structured arrangements of chemical moieties that promote binding to an inorganic phase and lower the interfacial free energy of nascent nuclei, templating heterogeneous growth below the homogeneous nucleation threshold.

The optimal surface for interacting with inorganic lattices is unknown, and hence we explored three design hypotheses and sought to generate proteins presenting 1) periodic arrays of charges (*1*, *14*), 2) residues partially coordinating metal ions (*15*), and 3) ordered hydration layers promoting adsorption (*16*, *17*) (**Fig. 1A**). We presented these interfaces on designed scaffolds with different topologies, periodicities, and curvatures. These scaffolds included primarily alpha helical designed helical repeat (DHR) proteins (*13*, *18*), designed alpha-beta proteins (*19–22*) (**Fig. 1, B and C**), and proteins containing flat beta-sheet surfaces: designed idealized beta-solenoid (DBS) repeat proteins (see Methods), a native antifreeze protein (RiAFP), and an idealized penta-peptide repeat (PPR) fold (*23*, *24*) (**Fig. 1D**). We selected two target materials that can be synthesized in aqueous solutions but are not known to occur as primary mineral phases in native bio-inorganic hybrid materials: zinc oxide (ZnO) and hematite (α-Fe_2_O_3_) (*25*). ZnO has diverse applications in emerging technologies (*26*), energy scavenging nanosystems (*27*), thin film transistors (*28*), field emitters (*29*), and can be detected using photoluminescence (PL) spectroscopy (*30*). Hematite is a hydrogen-generating photocatalyst (*31*) that can be distinguished from other iron oxides by color (*32*).

**Fig. 1.**
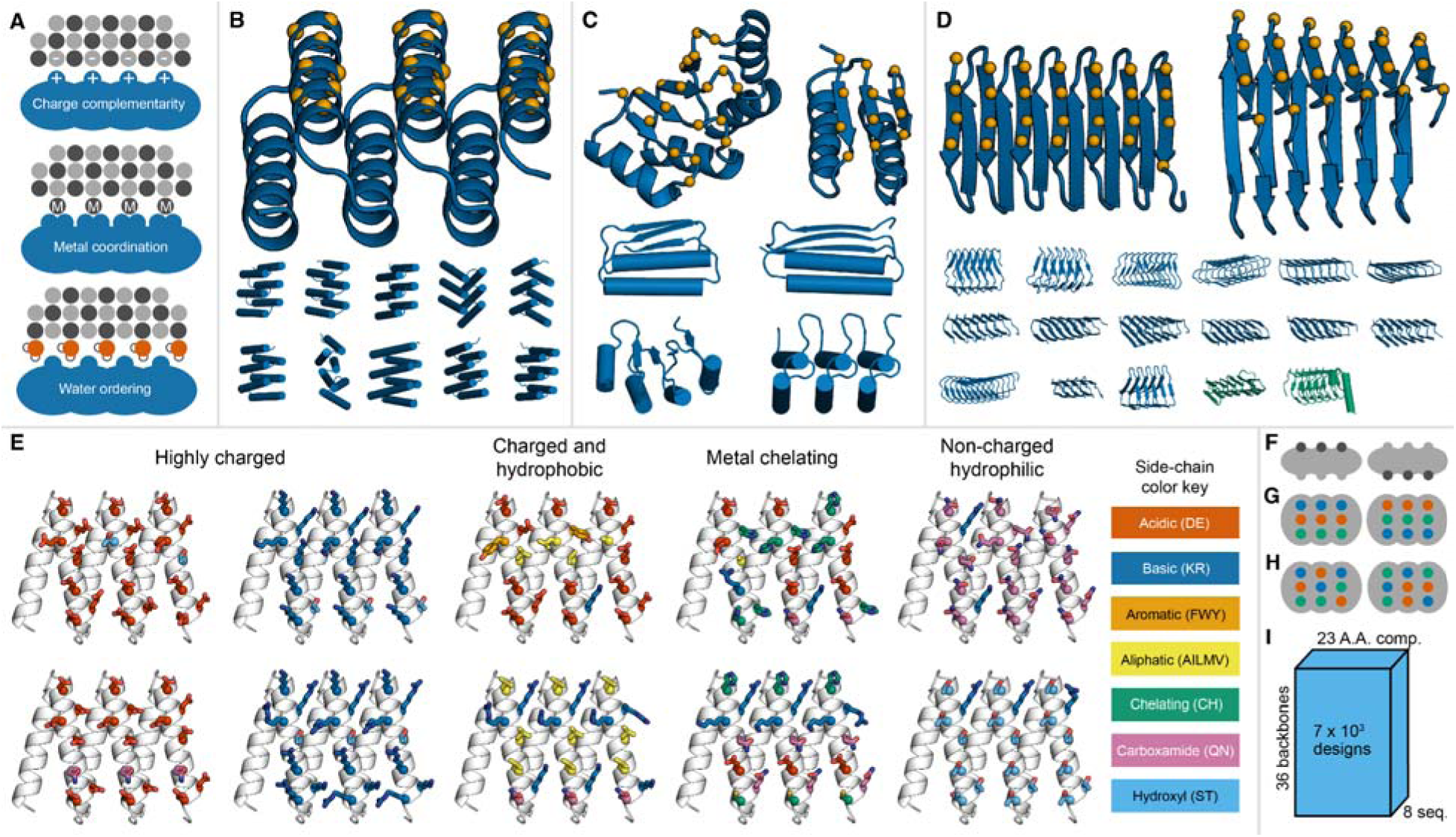
Design principles for templating mineral growth. (**A**) Three hypothesized mechanisms of mineral binding: charge complementarity, metal coordination, and water ordering. (**B to D**) 36 protein backbones used in the library. Orange spheres show alpha carbons of residues in a designed interface. Scaffolds include (**B**) 11 designed helical repeats (DHR), (**C**) 6 alpha-beta topologies, and (**D**) 17 designed beta solenoids (DBS) and 2 native solenoids (in green). (**E**) Example designed surfaces defined by 10 (out of 23; **fig. S1**) amino acid compositions applied to the DHR backbone DHR14. The colors of the amino acid side chains correspond to chemical categories identified in rectangles. (**F to H**) Illustration of the designs generated for each combination of scaffold and composition. (**F**) Two sets of interface residues are selected on opposing surfaces of protein. For each interface and amino acid composition, two repeat surfaces (**G**), with the same residue in analogous positions in each repeat subunit, and two non-repeat surfaces (**H**), wherein residues are scrambled among repeats, are produced. (**I**) 4 designed interfaces for each of 2 surfaces yields 8 sequences per combination of 36 protein backbones and 23 amino acid composition, resulting in a library of approximately 7 × 10^3^ designs.

To test the three above design hypotheses, we defined interface amino-acid compositions enriched in distinct amino acid residues (**Fig. 1A and fig. S1**): histidines and cysteines to coordinate metal ions, threonines to order water molecules as in ice-binding proteins, and negative and positively charged residues for charge patterning. We generated interfaces on each scaffold (**Fig. 1, B to D**) with each composition in both periodic and aperiodic arrangements (**Fig. 1, E to H**), resulting in 7 × 10^3^ designs (**Fig. 1I**).

We developed a yeast display flow cytometry (FC) screen that relies on the light scattering properties of nanoparticles (NPs) to identify and enrich yeast cells with increased binding propensity to ZnO or hematite NPs (**Fig. 2, A and B, and figs. S2 to S4**). Next-generation sequencing (NGS) was used to compare the abundance of designs in sixteen subpopulations of the library before and after sorting and identify enriched sequences (**Fig. 2C, figs. S5 and S6**; see Methods). Individual enriched clones were retested by FC on yeast (**Fig. 2C, figs. S7 and S8, and tables S1 to S6;** see Methods). The identified ZnO binders included eight DBSs, two DHRs, two alpha-beta topologies, and a surface redesign of RiAFP. Five of these surfaces contained histidine and cysteine residues, which can coordinate zinc ions (*33*), six contained large hydrophobic patches, five contained threonine motifs observed in ice-binding proteins (*16*), and eight contained periodic charges (**Fig. 2, D to H, fig. S9, and tables S1, S3, and S5**). We also identified hematite binders: two histidine-rich DHRs, two charge-rich DHRs, and three hydrophobic beta-solenoids (**Fig. 2I, fig. S10, and tables S2, S4, and S6**). We expressed these proteins in *Escherichia coli* (*E. coli)* with 6xHis tags and purified them with immobilized metal affinity chromatography (IMAC; **Fig. 2, D to F and I, and tables S7 and S8**).

**Fig. 2.**
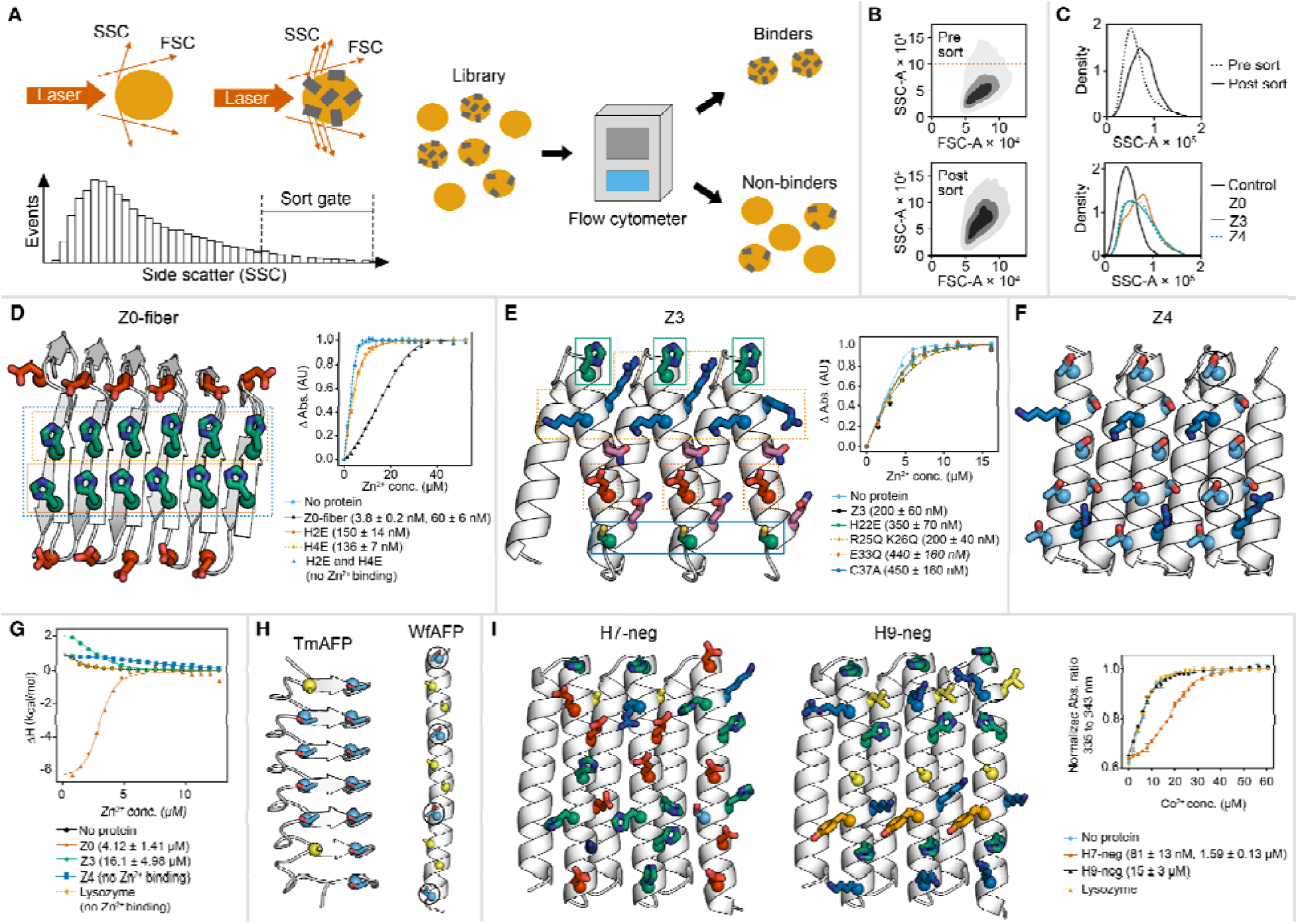
Mineral binding screen. (**A**) Illustration of screening strategy. (**B**) SSC-A vs. FSC-A signals of yeast populations before and after ZnO enrichment sorting. Plots are kernel density plot estimates from 3,507 and 2,576 events respectively. (**C**) SSC-A signals of an example sublibrary before and after sorting, and yeast clones displaying three enriched designs vs. control yeast. Yeast displaying both the designs and the control yeast displayed HA and c-Myc tags used to verify expression of the display system immunocytochemically, but the control yeast displayed no designed protein. (**D and E**) Z0-fiber and Z3 interfaces and mutational analysis of Zn^2+^ ion binding with ratiometric dye (Mag-Fura-2). Although mutants are named by the first residue that is mutated, all repeated instances of these residues, circumscribed by lines on the models, are mutated as well. (**F**) Model of the Z4 enriched design with circles highlighting threonines repeated in pattern resembling TXXXAXXXAXX motif in native antifreeze proteins (AFP). (**G**) Isothermal titration calorimetry analysis of Zn^+2^ against three designed proteins, lysozyme, and solutions without protein (protein concentration = 30 μM). (**H**) Example native ice-binding proteins: beta solenoid Tenebrio molitor AFP (TmAFP) containing threonine-rich parallel beta-sheets (PDBid: pdb_00001ezg) and alpha-helical winter flounder AFP (WfAFP) with threonines in TXXXAXXXAXX motif circled (PDBid: pdb_00001wfa). (**I**) H7-neg and H9-neg interfaces and analysis of Co^+2^ ion binding with ratiometric dye (Mag-Fura-2). Colors of all amino-acid side chains correspond to the chemical categories defined in Fig. 1E.

We selected five ZnO binding proteins (Z0, Z1, Z3, Z4, and Z5) with substantial FC binding signals to ZnO NP and which expressed well in E. coli as soluble proteins for further characterization (**figs. S11 to S13**). We also regularized existing head-to-tail interactions in Z0 to generate an unbounded fiber (Z0-fiber) (**fig. S14**). We selected two hematite binders (H7 and H9), and mutated non-interfacial positive residues to increase yield and solubility (*34*), producing the versions H7-neg and H9-neg (**figs. S15 to S17, and table S8; see Methods**). Design solubility trends and detailed biochemical characterization are described in the supplementary text (**tables S1 and S2**).

Using isothermal titration calorimetry (ITC), Z0-fiber was found to bind Zn^2+^ exothermically with a Kd of 4.1 ± 1.4 μM, and Z3 was found to bind Zn^2+^ endothermically with a Kd of 16 ± 5.0 μM (**Fig. 2G**; see Methods). A ratiometric dye (Mag-Fura-2) assay indicated multiple Zn^2+^ binding sites in both Z0-fiber and Z3, with nanomolar affinities (**Fig. 2, D and E**; see supplementary text). Interfacial mutations indicated the two rows of repeating histidines in Z0-fiber, and glutamate, cysteine, and histidine residues in Z3, are critical for Zn^2+^ binding, whereas basic residues in Z3 are not (**Fig. 2, D and E**). Mutating these histidines also reduced ZnO NP binding on yeast (**fig. S18**). No heat signal was detected upon titration of Zn^2+^ ions to Z4 (**Fig. 2G**), and the Kd was estimated to lie above the quantification limit of the Mag-Fura-2 assay (> 1 μM; **fig. S19**). Z4 contains a threonine motif found in alpha-helical ice-binding proteins (**Fig. 2, F and H**), and four other enriched proteins (Z8, Z9, Z12, and Z14) contain another threonine motif found in beta-solenoid ice-binding proteins (**Fig. 2H, fig. S9**)(*16*).

Ratiometric dye assays showed that both H7-neg and H9-neg bind Co²^+^ ions, a commonly used analog of Fe²^+^, with H7-neg exhibiting strong nanomolar binding affinity, but H9-neg only weakly coordinated Co²^+^ with micromolar affinity (**Fig. 2I)**.

### Assessment of metal oxide templating activity

We assayed the ability of the purified ZnO-binding proteins to promote the growth of ZnO NPs by incubating them in supersaturated solutions (3 mM Zn(NO_3_)_2_, 50 mM NaCl, 100 mM HEPES, pH 8.2), which allowed heterogeneous growth of ZnO, but not homogeneous growth (**Fig. 3, A and B**). We measured the photoluminescence (PL) of ZnO NPs to assess protein-induced growth and seeded positive controls with 12.5 μg/mL of commercial ZnO NPs, an ideal template. After 2 hours, supersaturated solutions containing lysozyme or no additive showed no PL, whereas solutions seeded with 0.1 mg/mL of Z0-fiber, Z3, or Z4 had PL spectra consistent with ZnO NPs (**Fig. 3B**)(*35*). X-ray diffraction of the material grown in the presence of Z0-fiber, Z3, and Z4 produced diffraction patterns indicative of ZnO (**Fig. 3C**). PL increased immediately in the ZnO NP control, but only after 30 to 60 minutes with the proteins (**Fig. 3D**). Additional control proteins BSA, DNase I, and DHR10-mica6 (*14*) had no observable effect on the PL signal (**fig. S20**). Z0-fiber showed growth in only two of six replicates, whereas all replicates containing Z3, Z4, and ZnO NP showed growth (**fig. S21**). To confirm the designed interfaces were responsible for the ZnO templating activity, we tested the motifs when hosted in different scaffold proteins. Reasoning that aggregation of Z0-fiber may be impeding its function, we extracted the putative functional motifs and designed topologies with RFdiffusion (*36*) and MPNN (*37*) to host the Z0 motifs in alpha-beta proteins (**Fig. 3E**). We expressed eleven of these ‘zab’ proteins in *E. coli* and selected four, zab3, zab5, zab10, and zab11, with CD spectra and SEC traces consistent with the models (**fig. S22**). Having observed ice-binding like threonine motifs in Z4 (**Figs. 2H and 3F**), we tested a previously designed ice-binding twistless-helical repeat protein, b-iTHR-201 (**Fig. 3G**), which presents a larger array of this motif (*38*). This ice-binding protein and two of the redesigns based on the Z0 interface (zab3 and zab10) showed strong ZnO templating signals (**Fig. 3, H to J**). Four previously reported ZnO binding peptides (*7–10*), fused to Small Ubiquitin-like Modifier (SUMO) to aid solubility and expression, showed no activity (**Fig. 3J**, **and table S9**). For replicates that triggered ZnO growth, we compared nucleation times (see Methods). Five design templates promoted an increase in photoluminescence at times greater than or equal to the control NPs, but the increase promoted by zab3 occurred substantially earlier (**Fig. 3K**).

**Fig. 3.**
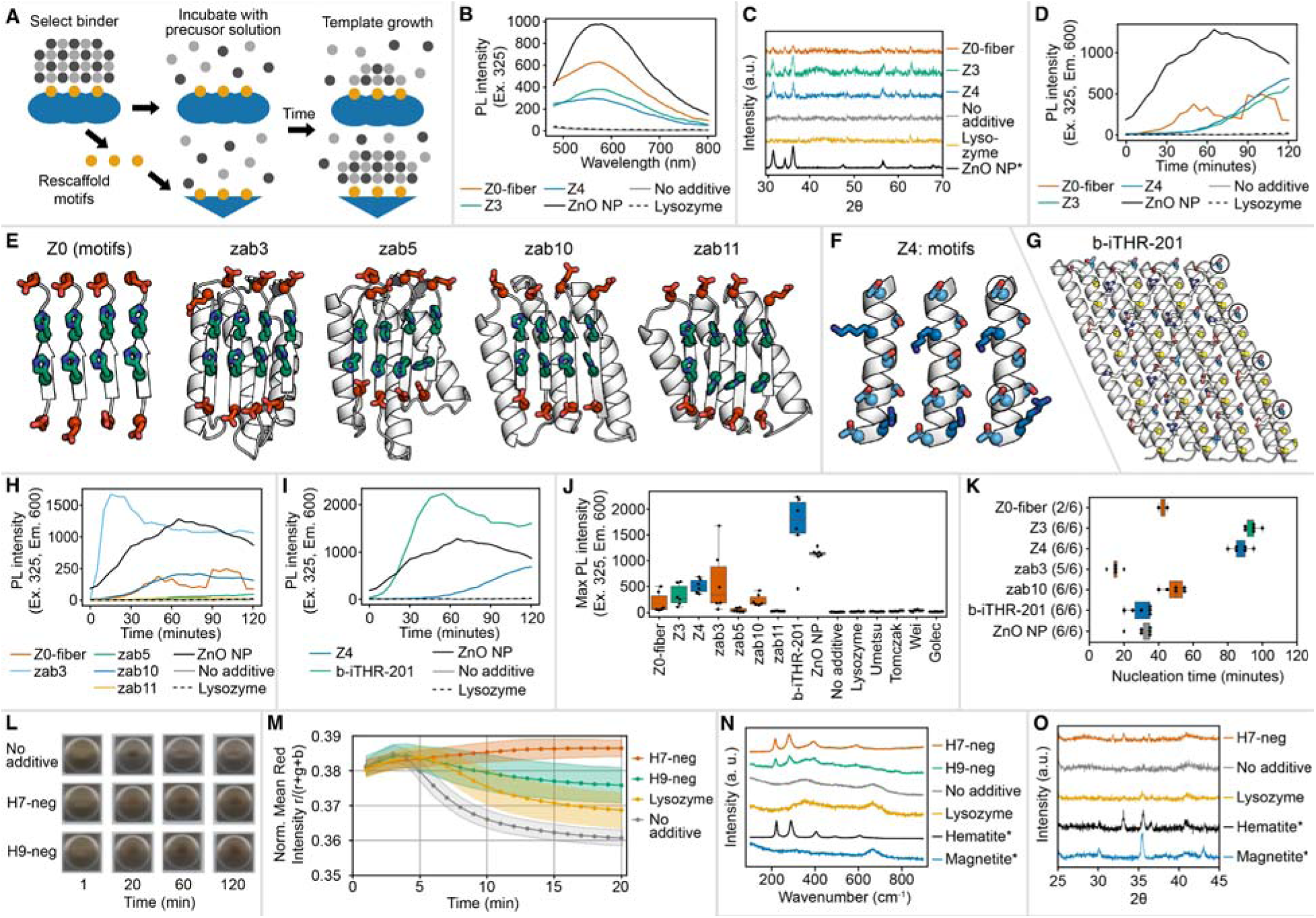
Metal oxide nucleation by designed interfaces. (**A**) Nucleation assay and motif grafting schematic. (**B**) Photoluminescence (PL) of ZnO-supersaturated solutions seeded with designs, ZnO nanoparticle (NP), lysozyme, or no additive. (**C**) X-ray diffraction of the ZnO nucleation products alongside ZnO NP* reference. (**D**) PL over time (highest of six replicates; see Methods). (**E**) Z0 interface motifs and four zab designs. (**F and G**) Z4 and b-iTHR-201 motifs resembling the TXXXAXXXAXX repeat of α-helical ice-binding proteins (circled in one repeat). Colors of side chains in all panels defined in Fig. 1E. (**H and I**) PL emission over time of zab designs, Z0-fiber, and controls (**H**) and b-iTHR-201, Z4, and controls (**I**). (**J and K**) Box plots showing 6 replicates of growth experiments using designed proteins, controls, and four previously reported ZnO-binding peptides (*7–10*). (**J**) Average maximum PL intensity (n=6). (**K**) Nucleation times for samples with maximum PL > 100, n as indicated. (**L**) Digital scans of iron oxide growth reactions under applied magnetic field at 1, 20, 60, and 120 min (**movie S1)** Magnetic material formed in no additive condition was pulled from the sample well by the applied field between 20 min and 60 min. (**M**) Quantitative colorimetric analysis of iron oxide growth reactions. Mean values (n=6), shading ±1 SD (see Methods). (**N**) Raman spectroscopy of iron-oxide reaction products with reference NP. (**O**) X-ray diffraction of the same products with hematite and magnetite NP references. Peak assignments in Methods. *indicates samples that are commercial NP references, not reaction products. All growth reaction conditions in Methods.

We examined if the H7 and H9 interfaces could direct the formation of hematite, the specific compound they were selected for, over other iron oxides. Operating under anaerobic growth conditions in a magnetic field, which typically yield magnetite (**fig. S23**), we found that supersaturated solutions containing H7-neg were more red in color after 20 minutes incubation, consistent with the presence of hematite, compared to supersaturated solutions (see Methods) containing H9-neg, lysozyme, no additive, or hematite or magnetite NPs (**Fig. 3, L and M, fig. S24, and movie S1**). Color was compared within the first 20 minutes, after which magnetite was pulled from solution by the magnetic field (**Fig. 3L and movie S1**). Raman spectroscopy of the reaction products indicated hematite was produced in solutions containing H7-neg and H9-neg, whereas magnetite was produced in solutions without protein or with lysozyme (**Fig. 3N and fig. S25)**. H9-neg also showed a minor unidentified ≈350 cm⁻¹ band. X-ray diffraction of material grown with H7-neg showed peaks corresponding to hematite but not magnetite; the hematite peak at 33.1° was not observed with lysozyme or no additive (**Fig. 3O**). SAXS analysis showed that H7-neg exhibited a shallower low-q power-law exponent (−2.26) than the lysozyme (−2.70) and no-additive (−2.67) controls, consistent with a more anisotropic or open nanoparticle morphology (fig. S26). The absence of pronounced form-factor oscillations further indicates a heterogeneous particle size and/or morphology distribution (fig. S26). After a 2-hour aerobic incubation, solutions containing H7-neg accumulated less rust at the iron-water interface than samples without protein, an effect not seen for lysozyme (**fig. S27**).

### Confinement of zinc oxide mineralization

We examined the effect of confinement on the rate of nucleation by lining the interior cavities of oligomers with ZnO nucleating interfaces. We chose Z4 as a starting point because its perpendicular-helix DHR topology is particularly amenable to oligomer design (*13*). We used RFdiffusion (*36*) and MPNN (*37*) to design cyclic oligomers to host the Z4 interface within the interior of rings (**Fig. 4, A to D**, see Methods). We experimentally tested twenty-three oligomers and selected three designs forming monodisperse species of the expected size, Z4-C3i, Z4-C3ii, and Z4-C6, for further characterization (**fig. S28**). The Z4-C3ii-XL trimer was derived from Z4-C3ii by adding another repeat of the Z4 interface to each monomer. The structures of Z4-C3i and Z4-C3ii were solved with X-ray crystallography, aligning to the design models with Cα root mean square deviations (RMSD) of 2.14 Å and 1.57 Å respectively (**Fig. 4, A and B, and table S10**). The 2mFo-DFc map of Z4-C3i showed electron density near the designed interface consistent with water ordering; however due to the limited resolution of the data (3.06 Å) we chose not to place waters at all locations during structure refinement (**fig. S29**). The structures of Z4-C3ii-XL and Z4-C6 were examined with ns-TEM, yielding 3D reconstructions with the expected triangular and hexagonal shapes of the trimeric and hexameric designs respectively, with slightly larger than expected diameters (**Fig. 4, C and D**). All four oligomers retained the templating activity of the Z4 monomer. Knockouts of Z4-C3i, Z4-C3ii, and Z4-C6, in which the interfacial threonines were mutated to glutamate (Fig. 4E), assembled correctly **(fig. S30)** but had no templating activity (**Fig. 4F**). Cryo-EM of ZnO NPs grown with Z4-C6 showed both empty rings and solid heterogeneous NP of the same diameter as the designed hexamer (**fig. S31**).

**Fig. 4.**
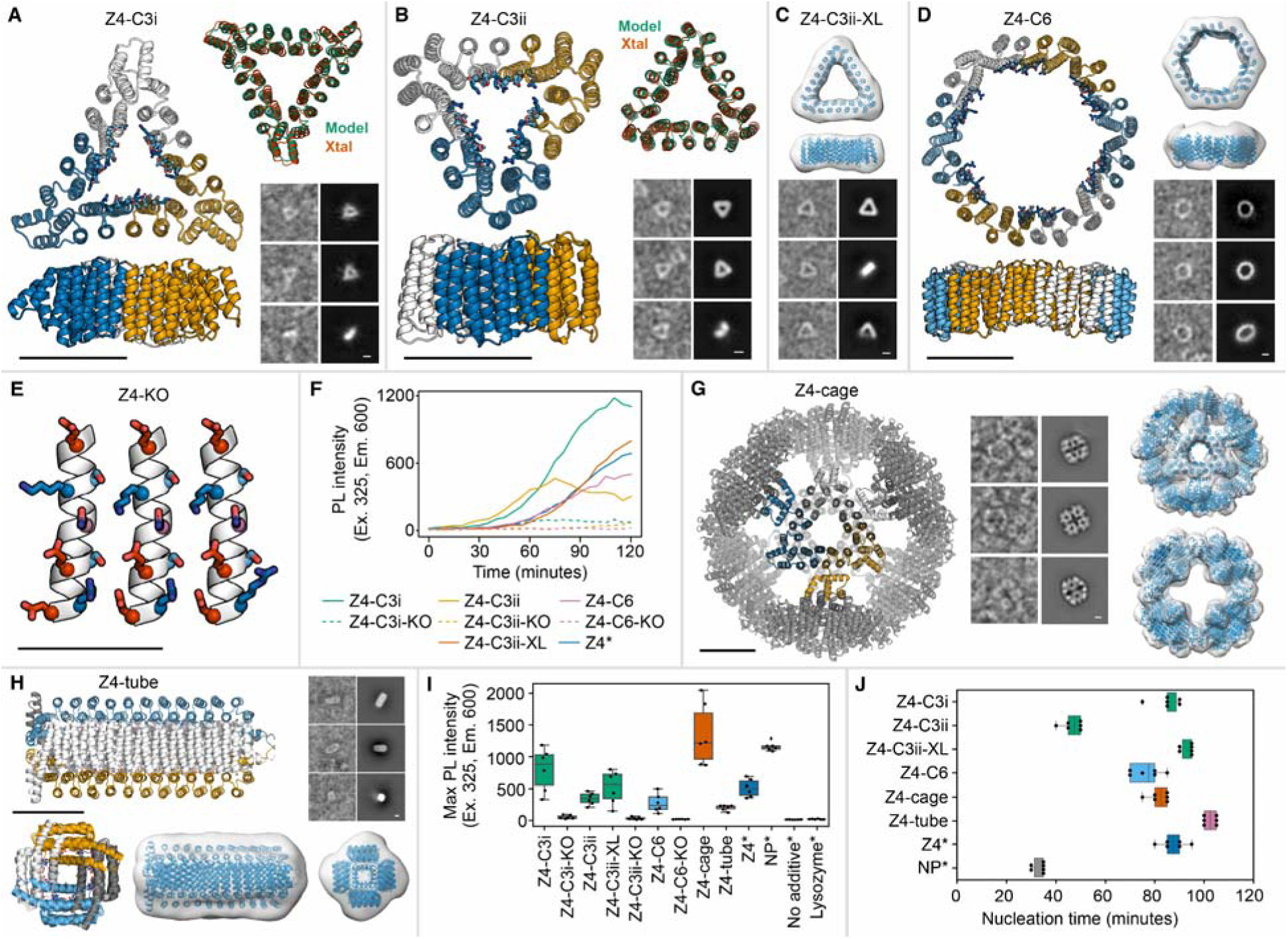
ZnO nucleation by Z4 motif-containing designed oligomers. (**A to D**) Cyclic oligomers with Z4 motifs lining interior cavities. (**A and B**) Two C3 designs with design models colored white, blue, and orange by subunit with Z4 motifs shown as sticks. Design model backbones shown in green superimposed with X-ray crystal structures in red (PDB IDs: pdb_00009D92, pdb_000037TT). The Cα RMSD between the design model and crystal structure was 2.14 Å for Z4-C3i and 1.57 Å for Z4-C3ii. Images at bottom right are representative ns-TEM picked particles (left) and 2D class averages (right). (**C**) Z4-C3ii-XL model superimposed with ns-TEM 3D reconstruction, and examples of picked particles and 2D classes. (**D**) Z4-C6 colored by subunit with motifs shown as sticks, model superimposed with ns-TEM 3D reconstruction, and examples of picked particles and 2D classes. (**E**) Model of the mutated Z4 knock out (Z4-KO) interface with threonines replaced with glutamates. (**F**) Plot of PL emission over time (Ex. 325, Em. 600) of Z4-motif containing cyclic oligomers compared to oligomers with interface KO mutants and the Z4 monomer (*sample replicated from Fig. 3; highest-signal replicate of six replicates shown; see Methods)). (**G**) Octahedral cage designed using Z4-C3i, ns-TEM picked particles and 2D class averages, and model superimposed with ns-TEM 3D reconstruction. (**H**) Z4-tube colored by subunit with Z4 motifs shown as sticks, ns-TEM picked particles, 2D-averages, and 3D reconstructions superimposed with models. (**I and J**) Effect of Z4-motif containing oligomers on ZnO nucleation compared to Z4 monomer, ZnO NP, and no additive solutions (**\***samples are replicated from Fig. 3). (**I**) Maximum PL intensity after nucleation assay (n=6). (**J**) Nucleation times for samples with maximum PL > 100, n as indicated (see Methods). Colors of amino acid side chains defined in Fig. 1E. All scale bars are 4 nm.

We next incorporated the ZnO templating interface into larger protein assemblies. To the Z4-C3i trimer, we incorporated a homodimeric protein-protein interface with an orientation that generates a 24-subunit octahedral nanocage, called Z4-cage (**Fig. 4G**). Z4-tube was designed by adding tetrameric domains to the termini of an extended version of Z4, such that ZnO-interacting interfaces surround a rod-like cavity (**Fig. 4H**). These were expressed in *E. coli* and purified with Strep-tag affinity chromatography and SEC (**fig. S32**) and ns-TEM was used to confirm the formation of the designed assemblies (**Fig. 4, G and H**). In the ZnO NP growth assay, Z4-cage promoted a large increase in PL, but Z4-tube showed a substantially lower increase (**Fig. 4I)**. There was no clear relationship between the size of the cavity or number of interfaces and the magnitude or rate of ZnO NP nucleation (**Fig. 4, I and J**).

We used small-angle X-ray scattering (SAXS) to characterize the heterogeneously nucleated ZnO products in solution to avoid drying artifacts. The low-q power-law exponents followed the order zab3 (−2.32) ≈ Z4-cage (−2.46) > Z0-fiber (−2.87) ≈ Z4 (−2.90), consistent with progressively decreasing structural anisotropy and increasing particle compactness (**fig. S33**). Form-factor oscillations were absent, indicating heterogeneous size and morphology distributions. To compare these products to a no additive condition, we increased precursor concentrations to allow homogeneous nucleation, which yielded a sample with a low-q power-law exponent of −4.86, suggesting unbounded aggregation (**fig. S33**).

### Cryo-EM characterization of protein-ZnO interface

We used cryo-EM to investigate the structure of ZnO nuclei within the Z4-cage. Although the majority of Z4-cage particles appeared empty in raw micrographs **(Fig. 5A)**, cryo-EM micrographs indicated stochastic NP growth originating from a subset of designed protein nanocages, with three distinct morphologies **(Fig. 5, A and B)**: first, within the cage (“filled”), second, growth extending outward (“blebbed”), and third, connecting sets of cages (“bridged”) **(Fig. 5B)**.

**Fig. 5.**
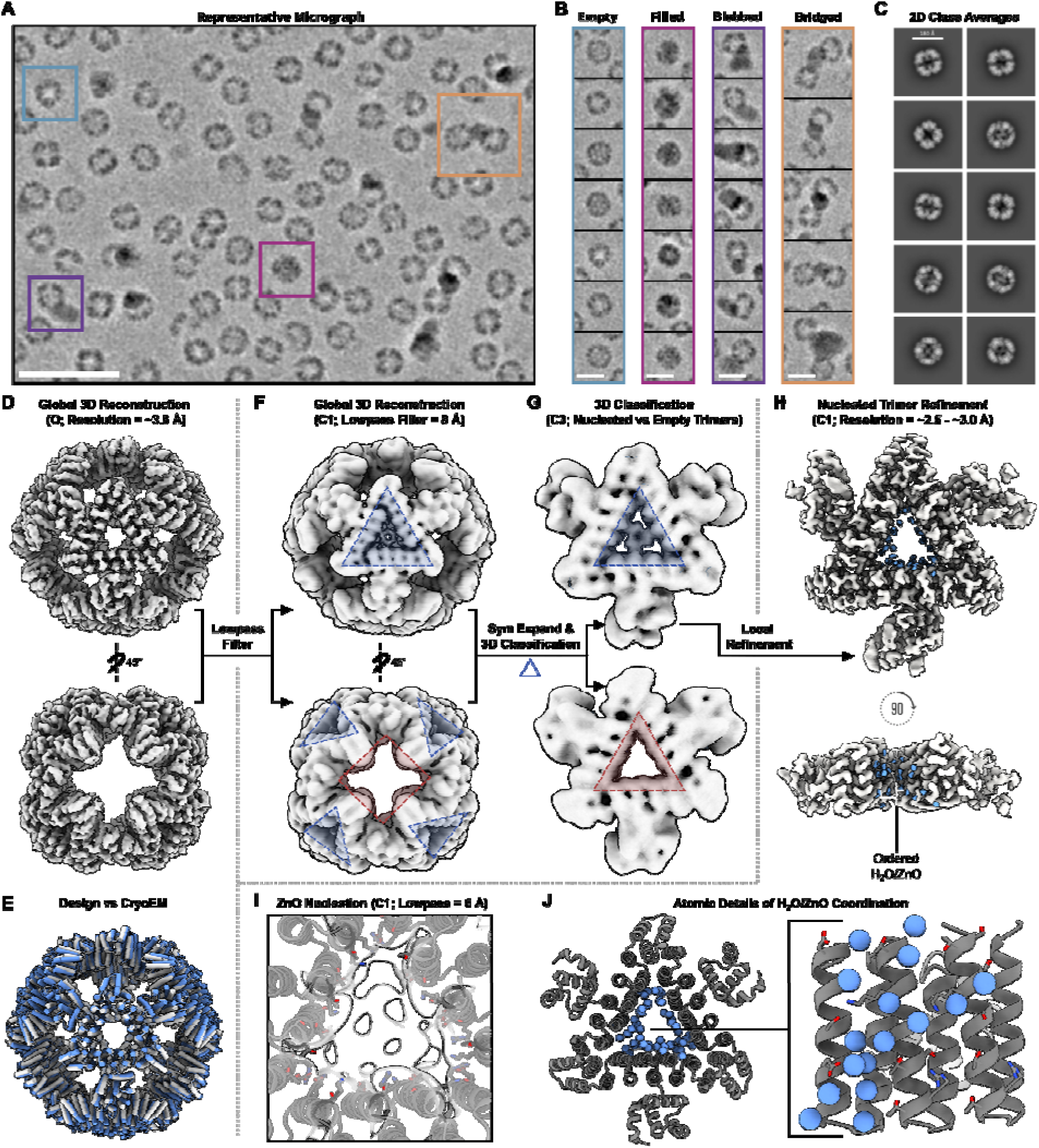
Cryo-EM characterization of ZnO mineralized Z4-cage. **(A)** Representative denoised micrograph of Z4-cage particles imaged in the presence of 3 mM Zn(NO_3_)_2_ for ∼30 minutes prior to freezing. Highlighted are examples of empty (teal), "filled" (maroon), "blebbed" (purple), and "bridged" (orange) nanoparticle morphologies observed in a single micrograph (scale bar = 100 nm). **(B)** Additional examples of each morphology observed in the dataset (scale bar = 180 Å). **(C)** 2D class averages of Z4-cage in the presence of nucleation conditions (scale bar = 180 Å). **(D)** 3.8 Å cryo-EM 3D reconstruction of Z4-cage refined with relaxed octahedral symmetry. **(E)** Comparison of the final built cryo-EM model of Z4-cage (grey) with the computational design model (blue). **(F)** C1 reconstruction of Z4-cage, low-pass filtered to 8 Å, showing ZnO nucleation localized along the C3 symmetry axis (blue), and no ZnO density along the C4 axis (red). **(G)** Octahedral symmetry expansion and 3D classification of all Z4-C3i trimers from all Z4-cage nanoparticles show both nucleated (blue) and empty (red) trimers exist within the data. **(H)** Near-atomic resolution 3D reconstruction of nucleated Z4-C3i trimers in the Z4-cage nanoparticle, highlighting density from coordinated H_2_O or ions (blue) near the designed ice-binding motif. **(I)** Low-pass filtered view of (H), zoomed in on the C3 axis, illustrating stochastic ZnO growth within the Z4-C3i trimer. **(J)** Placement of atoms into the density (see Methods) suggests that initial H_2_O/ion organization and nucleation occur along the designed Z4 ice-binding motif (sticks).

2D class averages of Z4-cage particles revealed secondary structure elements consistent with the design **(Fig. 5C and fig. S34)**, and 3D refinement with relaxed octahedral symmetry produced a ∼3.8 Å cryo-EM structure closely aligning with the design model **(Fig. 5, D and E, fig. S34, and table S11)**. However, asymmetric and stochastic NP growth hindered reliable analysis of ZnO density at this resolution. A global C1 3D refinement using a near-spherical mask allowed detailed and unbiased analysis of the pores along both the C3 and C4 axes together with an overall averaged representation of the Z4-cage **(fig. S35)**. The resulting ∼4 Å asymmetric map was low-pass filtered to 8 Å to highlight lower-resolution features corresponding to observed stochastic ZnO nucleation and growth patterns **(Fig. 5, A and B, and fig. S35)**. This approach revealed additional non-protein density, likely ZnO, localized along the center of each C3 axis and proximal to the designed ZnO binding interface **(Fig. 5F and fig. S35)**. In contrast, no extra density was observed along the C4 axes (distant from the designed ZnO binding interface) at any contour level **(fig. S35)**. To evaluate whether ZnO nucleation occurred uniformly across all C3 trimers, we next performed octahedral symmetry expansion followed by 3D classification for all Z4-C3i trimers within the Z4-cage. As expected, a subset of C3 trimers displayed ZnO density along the C3 axis, with several 3D classes appearing empty, consistent with both the empty particle morphologies and asymmetric growth of ZnO observed in raw micrographs **(Fig. 5, A and G)**.

3D local refinement of the threefold-symmetry axis of the mineralized Z4-cage yielded a high resolution cryo-EM map, ranging from 2.5 Å resolution in the center of the C3 axis to ∼3.5 Å at the interface between adjacent Z4-C3i trimers in the nanocage **(Fig. 5H, fig. S34, and table S11)**. Density consistent with solvent organization was visible along the ice-binding-protein-like threonine motifs in the Z4 designed interface (*16*), although the identity of the species could not be determined at this resolution **(Fig. 5, H to J; see Methods and supplementary text**). These results are consistent with stochastic ZnO nucleation within the trimers promoted by atomic ordering at the designed interface, followed by further ZnO growth into heterogeneous morphologies **(Fig. 5)**.

## Conclusion

Nature has evolved biomineralizing proteins that control the formation of hybrid organic-inorganic materials, such as bone and nacre, with greater complexity than accessible to conventional top-down manufacturing (*25*). Modern protein design tools readily program protein structures and protein-protein interactions, but inorganic-templating protein interfaces are inadequately understood. Our high-throughput design and evaluation approach to identifying protein interfaces that promote mineralization of zinc and iron oxides provides a route to overcoming this limitation that can potentially be extended to a wide range of inorganic crystals. These interfaces control whether nucleation occurs, which phase forms, and where it forms within an assembly. Future work will aim for complete control over final particle size, shape, and dispersity. In a complementary co-submission (*39*), we show that protein assemblies can template and confine calcium phosphate, a biologically relevant mineral, within defined nanoscale compartments. Together, these studies open the door to a world of designed hybrid materials containing inorganic components–including those not explored by nature–that are controlled by de novo proteins.

## Materials and Methods

### Library design

Our library of 6730 proteins with potential mineral-binding interfaces was designed as follows. 36 scaffold proteins (**Fig. 1, B to D**) were selected to host the interfaces, including 11 designed helical repeats (DHRs) (*1*, *13*, *18*), 17 designed beta-solenoids (DBS) created for this work, 1 native beta-solenoid antifreeze protein (*23*), an idealized beta-solenoids derived from a PPR protein (*24*), 3 alpha-beta proteins generated by Foldit users (*19*, *21*), and 3 repeat proteins with alpha-helical and beta-sheet character (*20*, *22*).

23 sequence compositions (**Fig. 1 and fig. S1**) were defined and used to design interfaces containing diverse surface chemistries on each scaffold. Interface design was performed with RosettaScripts using amino acid composition (AA comp) constraints to apply the defined sequence compositions (*40*). Interfaces were designed on surfaces designated ‘surfA’ and ‘surfB’ on opposite sides of the scaffold proteins, or surfaces of different sizes on scaffolds with internal pseudosymmetry (i.e., proteins where both surfaces contain the same backbone structure). One interface design protocol enforced repeat symmetry by requiring the designed interface to contain the same sequence in every repeat unit, while another protocol broke repeat symmetry as much as possible while still containing the defined sequence composition. These two protocols generated designs designated ‘repeat’ and ‘antirep’ respectively. The two designs with the lowest Rosetta energy (total_score term with beta_nov16 scorefunction) for each combination of scaffold, AA comp, surface, and repeat symmetry were selected for inclusion in the library. Multiple versions of the DBS scaffolds with different surface chemistries outside of the designed mineral-binding surfaces were used to host the Thr90 AA comp as we reasoned that firstly this combination of topology and sequence composition was particularly promising because it resembles native antifreeze proteins that order waters on their surface (*16*) and this may promote binding to ordered waters on mineral surfaces (*17*) and secondly these DBS scaffolds were unverified so including multiple scaffold sequences for each backbone would increase the likelihood of finding successful binders. Sequences outside the binding surface remained fixed and the original scaffold sequences were also included in the library.

This combination of 36 scaffolds * 23 compositions * 2 surfaces * 2 repeat symmetries * 2 designs yielded 6624 designs. The original scaffolds and additional DBS Thr90 designs (described above) were added to this set, which was then filtered to 6730 designs by removing designs with less than 5 amino acid differences to any other sequence in the library. Designs were named with the format of: (scaffold)_(AA comp)_surf(A or B)_(repeat or antirep)_thread(#)_####, where # are arbitrary numbers to identify particular designs in each category. The design models in the library as well as the protocols and input files used to generate them are available in the GitHub repository associated with this publication (https://github.com/pylesharley/protein_mineral_library) and Zenodo archive (*41*).

### Designed beta-solenoid repeats

Designed beta-solenoid repeat proteins were generated by an iterative in silico evolution process, conceptually similar to previously reported strategies (*12*). Sequences were initialized by concatenating random stretches of beta-sheet propensity motifs (QxQx, where x is a hydrophobic residue) with random stretches of loop propensity residues (GSTPND), and repeating the sequence in tandem as many times as possible while remaining under 200 residues. The structures of these sequences were predicted with RoseTTAFold (*42*) and the best scoring models (models in the 95th percentile of both cce and kl scoreterms) were selected. These sequences were mutated while maintain repeat symmetry and the overall pattern of chemistries (hydrophilic residues to hydrophilic residues, hydrophobic residues to hydrophobic residues, loop residues to loop residues), and the RoseTTAFold prediction and sequence selection steps repeated 3-5 times. Next, the structures of the sequences were also predicted with AlphaFold2 (*43*) and the designs with the lowest root-mean-squared deviation (RMSD) between their RoseTTAFold and AlphaFold2 models were selected for visual inspection. 17 designs with large flat surfaces and well packed hydrophobic cores were selected as scaffolds, and named based on the base sequence repeat used to generate their structure, for example FQIGSSGQ and QAQLQIQASGT designate beta-solenoids comprised of 8 and 11 residue repeats, respectively. Lastly the surfaces of the beta-solenoids were redesigned with RosettaScripts to be hydrophilic and non-repetitive (*40*). Further details and an implementation of this protocol are available in the GitHub repository associated with this publication (*41*).

### Incorporating identified motifs into protein monomers and assemblies

ZnO binding motifs identified in the proteins Z0, Z3, and Z4 were incorporated into protein assemblies using several design strategies. Both the full backbones, and the isolated ZnO interface motifs, were incorporated using different strategies, and different approaches were used for the DHR and the DBS based designs.

As the DBS topologies showed a strong tendency to form fibers and larger bundled assemblies of fibers, we sought to transfer the parallel beta-sheet ZnO binding motif from Z0-fiber into a mixed alpha-beta topology. With the proper input variables, see GitHub repo (*41*), RFdiffusion generated alpha-helices that connected the motifs and preserved their structure and relative orientations. These Z0-motif alpha-beta proteins were designated zab proteins.

Cyclic oligomers containing the full DHR backbone and ZnO binding surfaces of Z4 were generated by positioning the original backbone in a desired cyclic confirmation, cleaving each source chain to two fragments at a central loop and then applying RFdiffusion (*36*) to link the two fragments together. Orientation of source chains into the desired cyclic positions was done by first identifying the C-alpha coordinates of the helices containing the ZnO interface. Using these coordinates, the center of mass of the helices was computed. Singular value decomposition (SVD) was then applied to compress the 3D coordinates into a 2D plane that exhibits the highest variance. A normal vector, representative of a 2D plane, was utilized to calculate a rotation matrix, aligning the normal vector with the x-axis. Subsequently, another SVD was performed to compress the 3D coordinates into a 2D plane, this time with the lowest variance. The corresponding normal vector from this operation was aligned with the y-axis. Upon visual inspection, if the active helices were not oriented inward towards the oligomer, a 180-degree rotation around the y-axis was executed. Translations along the x-axis were sampled as needed. To create symmetric new chains with respect to the z-axis, a series of rotations were carried out, the number of which was determined by the desired oligomeric state. A central loop at the center of each chain was then used to define two segments in each chain. A linker between a chain 1 segment B and chain 2 segment A was generated by RFdiffusion (*36*). MPNN (*37*) was used to generate numerous sequences for the diffused section in each cyclic unit. Designs were then selected based on RosettaFold and AlphaFold2 metrics followed by visual inspection.

### Design of large protein assemblies

The octahedral nanocage design, Z4-cage, was designed by taking the Z4-C3i model and docking it against a set of hallucinated C2 structures (*12*) using the “O32” architecture in RPXdock. Docking outputs were then processed with RosettaScripts, where the ConnectChainsMover was used to make a short helix-loop-helix connection between adjacent termini of the two docked components (*40*). Dock contacts and the new loop were sequence-redesigned with ProteinMPNN.

The C4 tube design, Z4-tube, began with extending the Z4 repeat protein model with the RepeatPropagationMover in RosettaScripts (*18*). The extended repeat was then oriented manually in PyMOL such that (i) the repeat extension trajectory was approximately parallel to the Z axis, (ii) the motif surface was facing inward toward the origin, and (iii) there was appropriate offset from the origin such that if a C4 symmetrizing operation were applied, a hollow tube model would be generated that displays the motif inward and has barely-contacting cyclic contacts between copies. 3 spacings were chosen for downstream sampling, with the intent that these loop contacts along the long axis would not be the driving force for interaction, but would just be close enough to enclose the interior. To enclose the ends of the tube and to provide favorable interactions to promote C4 assembly, the N and C terminal ends of the symmetric tube models were separately used as inputs for symmetric RFDiffusion, where a new C4 backbone was built in-place (16). These terminal C4 segments were independently designed with ProteinMPNN and filtered by AlphaFold2 metrics for folding 4 chains together. Then, combinations of N and C terminal end-capping structures were appended to the parent tube models to make the full structures.

### DNA design and yeast library construction

DNA sequences encoding 6730 library design proteins were generated by DNAworks with additional GS padding, adapters and peptide barcodes, added for length adjustment and to streamline data collection. Oligos were sorted into 16 subpools, divided into two fragments and optimized using OligoOverlapOpt (*44*), to minimize hybrid formation.

DNA fragments up to 300 bp in length were synthesized by semiconductor-based synthetic DNA manufacturing. Fragment pairs were amplified and combined following a standard assembly procedure (*45*). In short, oligos of interest were amplified via polymerase chain reaction (PCR) using outer and pool-specific primers (’qPCR1’). Subsequently, to minimize background oligos, the initial PCR reaction was diluted and amplified again (’qPCR1b’). Following PCR clean-up, the amplified oligos underwent USER digest and NEBNext repair to eliminate uracils and other impurities. The DNA fragments were then assembled and amplified to generate full-length oligos (’qPCR2’). Gel electrophoresis was utilized to visualize and isolate the assembled bands, which were subsequently extracted and eluted to obtain the final DNA product. Finally, scale-up PCR reactions were performed, followed by PCR cleanup to obtain the desired DNA libraries. Sixteen subpools libraries were then cloned to pETCON3, a yeast display vector derived from pETCON, by gibson assembly and transformed into EBY100 yeast using Gene Pulser Xcell electroporator (*46*).

### Yeast display

Yeast display and expression sorting was performed as previously described (*47*). Saccharomyces cerevisiae EBY100 strain cultures were grown in C-Trp-Ura medium supplemented with 2% (w/v) glucose. For induction of expression, yeast cells 500ul of C-Trp-Ura cell culture were transferred into tubes containing SGCAA medium supplemented with 0.2% (w/v) glucose and induced at 30 °C for 16–24 h. For determination of expression cells were washed with TBS (Tris 30mM NaCl 150mM) and labeled with anti-c-Myc labeled with Alexa-Fluor (Alexa647, 9B11, Mouse mAb, Cell-Signaling) and anti-HA Phycoerythrin (PE, HA-Tag (C29F4) Rabbit mAb, Cell-Signaling).

### Mineral screen

The flow cytometry based ZnO/Hematite binding screen was conducted as follows. First individual subpools expressing library clones were mixed with a solution containing ZnO nanoparticles (NP) from Sigma-aldrich (cat. 721077) or Hematite NP from US Research Nanomaterials, Inc (cat. US3981). Specifically, 100 μL of yeast culture from each library subpool at an optical density of 5 (OD 5) were resuspended in sort buffer solution (Tris NaCl buffer at pH 8.0) containing 0.02 wt/vol% ZnO nanoparticles ≤40 nm avg. part. size, or 0.02 wt/vol% Hematite nanoparticles (≤100 nm average particle size). Following an incubation period of one hour, yeast-ZnO/yeast-Hematite samples were resuspended in a 1200ul sort buffer solution and subsequently sorted utilizing the Sony SH800 Cell Sorter. Initial protein expression selection based on HA/Myc-tag fluorescence was followed by a series of three back-scatter/side-scatter (BSC/SSC) based sorting and growing enrichment cycles. Each enriched subpool was then evaluated in comparison to the initial expression library using BSC/SSC signal. Because yeast aggregates introduce noise to the assay, two initial filtration gates were added. First a forward-scatter area (FSC-A) gate on the main events population; second a singlet events gate using the forward-scatter height (FSC-H), forward-scatter area (FSC-A) correlation (**fig. S2**).

### NGS Analysis

Analysis of the next generation sequencing (NGS) results taken from the naive and enriched libraries was performed as follows. First DNA sequences predicted to be in the library were assembled *in silico* using BLAST+ 2.12.0 (*48*) to identify homologous overlaps between the purchased DNA sequences. In addition to the sequences encoding the library designs, some predicted chimera sequences containing the N and C terminal halves of similar designs identified and designated (design1)_CHIMERA_(design2). Next the NGS reads from populations of yeast sorted for expression of the protein on their surface (the naive condition) and for promoting adsorption of the ZnO nanoparticles for each of the library’s 16 subpools matched against the predicted sequences with BLAST. The genes encoding the designs were long relative to the length of the sequencing reads and the 5’ and 3’ paired reads often did not overlap. Therefore, the 5’ and 3’ paired reads were matched with BLAST to the designed sequences separately, and instances where both reads matched the same sequence with >90% sequence identity were counted as that design (or predicted chimera). Instances where the forward and reverse reads matched different sequences were classified as observed chimeras and designated (design1)_chimera_(design2) This permissive assignment strategy addressed the problem of read length and allowed the majority of the sequencing reads in the library to be assigned rather than a very small subset of them.

Sequences that were highly prevalent within sorted subpools showing an increased SSC signal were identified, and alignments of the NGS reads assigned to these sequences were manually inspected to check for point mutations, frameshifts, and nonsense mutations. As this was an exploratory screen, we considered mutants as well as the designed sequences for further study. For abundant mutated, truncated, or chimeric sequences in the sorted subpools, we tested the observed sequences as well as the design that they had the highest identity too (**tables S4 to S6**). For repeat protein designs, we also ordered ‘fiber’ versions in which the sequence based capping features in the N-terminal and C-terminal repeats were removed to allow head-to-tail interactions resembling the packing between adjacent repeats within the protein (**tables S4**). Alphafold2 (*43*) predicted models for all of these sequences are available in the GitHub repository (https://github.com/pylesharley/protein_mineral_library/tree/main/selected_designs)(41).

### Verifying activity of sequences in clonal yeast

Sequences of interest were ordered as gBlock fragments (Integrated DNA Technologies) with complementary overlaps tailored for assembly into a yeast display plasmid bearing an N-terminal HA tag and a C-terminal Myc-tag (pETcon3). Following assembly, transformed clones were labeled with anti-c-Myc labeled with Alexa-Fluor (Alexa647, 9B11, Mouse mAb, Cell-Signaling) and anti-HA Phycoerythrin (PE, HA-Tag (C29F4) Rabbit mAb, Cell-Signaling) and screened for surface expression of the desired protein using InCell Analyzer 2500 HS fluorescence microscope. These clones were compared to a control yeast strain displaying the same c-Myc and HA tags but no designed protein. Protein verification of clonal sequences was conducted using Sanger sequencing.

To assess the binding of individual yeast clones to ZnO and hematite nanoparticles, 100 μL of yeast culture (OD 5) was pelleted by centrifugation and resuspended in Tris-NaCl buffer (pH 8.0) containing either 0.02 wt/vol% ZnO nanoparticles (≤40 nm average particle size) or 0.02 wt/vol% hematite nanoparticles (≤100 nm average particle size). Following 1 hour of incubation, yeast-nanoparticle suspensions were resuspended in 1,200 μL of sorting buffer. Binding was then analyzed using a BD FACSymphony A3 Cell Analyzer flow cytometer, and data were processed using FlowJo software.

### Small scale protein expression and screening

Small scale protein expression screening was done as previously described (*12*). In short, Golden Gate reaction products were transformed into BL21(DE3) (New England Biolabs C2527I) and cultured using a 96 well plate format. Soluble fractions were purified by either 6xHis tag-based immobilized metal affinity chromatography (IMAC) or Strep affinity purification. For IMAC, clarified lysate was applied to Ni-NTA resin, the resin was washed with wash buffer (20 mM Tris, 150 mM NaCl, 30 mM imidazole, pH 8.0), and protein was eluted in the same buffer containing 150 mM imidazole. Proteins carrying a Strep-tag were purified using Strep-Tactin resin (0.5 mL resin per well). The resin was equilibrated with wash buffer (100 mM Tris-HCl, 150 mM NaCl, 1 mM EDTA, pH 8), and clarified lysate was applied by gravity flow to maximize binding efficiency. After loading, the resin was washed twice with wash buffer, and bound protein was eluted with elution buffer containing 100 mM Tris-HCl, 150 mM NaCl, 1 mM EDTA, and 2.5 mM desthiobiotin, pH 8 (*49*). Elution fractions containing the target protein were pooled and further purified by size-exclusion chromatography where indicated. Expression and solubility screening of purified samples was conducted using SDS-PAGE (Biorad Criterion 26 well stain free - anykD, #5678125) analysis.

### Large scale protein purification

Large scale protein expression was based on a method previously described (*12*). Full genes or golden gate reaction products were transformed into BL21(DE3) (New England Biolabs C2527I), grown overnight on kanamycin plates for single colony selection, and cultured using 50-500 milliliter cultures of Studier autoinduction media growth media supplemented with 0.1 mg/ml Kanamycin. For cell lysis cell pellets were resuspended in 10 or 25 mL wash buffer (20 mM Tris, 150 mM NaCl, 30 mM Imidazole, pH 8.0) on ice, supplemented with 0.1 mg/mL Lysozyme, 0.01 mg/mL, Deoxyribonuclease I (DNAse I, Millipore Sigma), 1 mM PMSF). Lysis was conducted via either by incubating the suspension at room temperature for two hours following addition of TritonX-100 (4% [v/v]) or by ice cold sonication (Qsonica, Q500 with a: 4-pronged horn) with 1 s ON, 1 s OFF, at 80 % amplitude for 5 minutes. Subsequent to lysis samples were centrifuged for 30 minutes at (25,000 x g ) to remove the insoluble fraction. Soluble fractions were then purified by either 6xHis tag-based immobilized metal affinity chromatography (IMAC) or via Strep affinity tags purification with Strep-tactin beads.

### Size Exclusion Chromatography (SEC)

IMAC or strep eluates were sterile-filtered through a 24 well filter plate (0.2 μm polyethersulphone (PES) membrane, Agilent). SEC was then performed using an autosampler-equipped Akta pure system (Cytiva) on a Superdex S200 Increase 10/300 GL/Superdex S75 Increase 10/300 GL (GE Life Sciences) columns at room temperature. Running buffer was 50 mM NaCl, 100 mM HEPES pH 8.2. Purified fractions were confirmed by reverse-phase liquid chromatography–mass spectrometry (LC-MS), and stored at 4 °C before downstream experiments.

### Circular dichroism

Circular dichroism analysis was conducted using an AVIV Model 420 circular dichroism spectrometer. The samples were prepared in either 10 mM NaPi pH 7 or 20 mM Tris pH 8 buffers and placed in a 1-mm quartz cuvette for measurement.

### Isothermal Titration Calorimetry (ITC)

Characterization of binding affinity was conducted as follows. First purified protein and control samples were equilibrated by repeated cycles of buffer exchange with a metal-free buffer solution containing 100 mM HEPES and 50 mM NaCl. Because high concentration of Zn ions is unstable at pH 8, all buffers and solutions were adjusted to pH 7.2. Subsequently, protein samples at a concentration of 30 μM underwent overnight treatment with chelex100 to ensure removal of any residual metal contaminants. Protein samples and control samples were then assayed using a microcal PEAQ ITC-Automated instrument. The assay protocol entailed nineteen successive injections of 2 mM Zn in buffer at 25°C. Resulting titration data was analyzed utilizing the Malvern Panalytical ITC Analysis software, and analyzed by a single-site binding model.

### Evaluation of metal ion binding by ratiometric dye

The binding of Co^2+^ and Zn^2+^ ions by proteins was assessed through a competition assay employing the ratiometric dye Mag-Fura-2. The assay relied on the significant absorbance alteration at 323 and 335 nm upon Zn^2+^ and Co^2+^ ion binding, respectively. For Zn^2+^, binding was measured as the normalized change in absorbance at 323 nm. For Co^2+^, binding was measured as the normalized ratio of absorbance at 335 nm to 343 nm. Measurements involving Zn^2+^ ions were carried out in a metal-free buffered solution comprising 100 mM HEPES and 50 mM NaCl at pH 8.2, and Co^2+^ ion binding was assessed in a metal-free buffered solution comprising 100 mM CHES and 50 mM KNO_3_ at pH 8.7. Both buffer systems have been demetalated using Chelex-100 prior to utilization.

Initially, the dissociation constant of the dye (Kd(Mag-Fura-2, Zn^2+^) = 10 ± 4 nM) was determined through competition with nitrilotriacetic acid (Kd(NTA, Zn^2+^) = 529 pM under the experimental conditions of I = 0.05 M, pH 8.2, and 25 °C) as well as through a direct titration with Co^2+^ (Kd(Mag-Fura-2, Co^2+^) = 840 ± 90 nM). Subsequently, the dissociation constants for the proteins were determined utilizing this dye.

In each Zn^2+^ ion binding measurement, the proteins, Mag-Fura-2, and/or NTA were deployed at a concentration of 5 µM in a 200 µL buffered solution. Following this, 1 µL of a 250, 500, or 1000 µM Zn(NO_3_)_2_ solution was stepwise added to the solution. The solution was shaken for at least 15 minutes in the dark between each addition to facilitate equilibration. In each Co^2+^ ion binding measurement, the proteins and Mag-Fura-2 were deployed at a concentration of 10 µM in a series of buffered solutions (each 10 µL) with varying Co^2+^ ion concentrations. The respective solutions were incubated in the dark and monitored until an equilibrium was reached. Duplicate measurements were conducted for each experiment.

### X-ray diffraction (XRD)

For ZnO growth reactions, protein samples (0.3 mg/ml) suspended in a buffer solution consisting of 100 mM HEPES and 50 mM NaCl at pH 8.2 were combined with 30 mM Zn(NO_3_)_2_ in a 1:10 ratio, resulting in a final concentration of 3 mM Zn^2+^. To produce larger quantities of ZnO, the mixture was then incubated at 50°C for 20 minutes within a Synergy NEO2 plate reader. Samples from the nucleation assay were then deposited onto 0.45 μm nitrocellulose filters (Whatman Protran 85) using a whatman manifold setup (Whatman Minifold I, Dot-Blot system). Samples were washed twice with 400 µl buffer to remove residual zinc ions. X-ray diffraction (XRD) measurements were conducted with the wet filter serving as a background substrate. A reference ZnO NP sample was produced by drop casting ZnO NP onto filter paper. Diffraction patterns were acquired using a Bruker D8 Discover X diffractometer equipped with a 2 mm collimator. Scans were executed within the range of 20 to 86 degrees, with an 11-degree increment and a read time of 30 seconds. Spectral analysis and background subtraction were carried out using Diffrac Suite EVA software.

For iron oxide growth reactions, the same protocol as described for the growth assay was followed with a few modifications. After 1 h of reaction, 5 μL of 5 M KOH was added to each well, and the plate was incubated at 50 °C for 1 h to terminate the reaction. Samples were then deposited onto a 0.4 μm nitrocellulose membrane and washed once with 1 M NaCl, followed by several washes with double-distilled water to remove residual buffer and salts. Membranes were kept wet throughout the measurements. X-ray diffraction (XRD) measurements were performed directly on the wet membranes using a Bruker D8 Advance diffractometer equipped with a LynxEye XE-T detector and a Cu Kα X-ray source. Reflections at 33.1° and 36.4° correspond to hematite and are present in the H7-neg product. The 33.1° reflection is absent from the lysozyme and no-additive products. Magnetite reflections at 30.1° and 43.1° are absent from the H7-neg product. The peak at 31.8° is attributable to NaCl from residual buffer salt.

### Raman spectroscopy

X-ray diffraction (XRD) analysis of small iron oxide nanoparticles is complicated by fluorescence interference, which limits the signal-to-noise of the resulting patterns. We therefore also used Raman spectroscopy, a well-established technique for fingerprinting polymorphs, to assess the phase composition of the different samples. For Raman measurements, H7-neg, H9-neg, and control samples were incubated for 3 hours with Fe intermediates under the conditions detailed in the Iron oxide growth assay Methods section. Following incubation, 100 µL aliquots were collected and mixed thoroughly by repeated pipetting to ensure sample homogeneity. To minimize drying effects and remove excess ions and buffer components, the nanoparticle pellets were washed multiple times with Milli-Q water. The samples were then resuspended in 30 µL of ethanol, mixed, and 10 µL was cast onto a glass microscope slide for analysis. Raman spectra were acquired using a 785 nm laser, scanning between 100–1000 cm⁻¹, with an X50 objective lens focused on nanoparticle aggregates. Measurements were conducted on H7-neg, H9-neg, control samples, and standards for comparative analysis.

### Small-Angle X-ray Scattering (SAXS) analysis

ZnO mineralization reactions were performed by mixing protein samples (0.3 mg mL⁻¹) in 100 mM HEPES, 50 mM NaCl (pH 8.2) with 30 mM Zn(NO_3_)_2_ at a 1:10 volume ratio, resulting in a final Zn²⁺ concentration of 3 mM. The reaction mixtures were incubated at 50 °C for 20 min to generate sufficient quantities of ZnO for SAXS analysis. To obtain a protein-free reference, precursor concentrations were increased twofold to permit homogeneous nucleation. Following mineralization, samples were transferred into glass capillaries and measured at 4 °C using a Xenocs Ganesha SAXS instrument or Xenocs Xeuss 3.0 laboratory SAXS instrument with acquisition time of 30 min per sample. Corresponding reaction buffer samples were measured under identical conditions and used for background subtraction. Data reduction, background subtraction, and analysis were performed using RAW software. Where indicated, the linear low-q region was fitted with a power-law function to determine the scattering exponent.

For iron oxide samples SAXS measurements were performed using a Xenocs Ganesha SAXS instrument. Samples were prepared as described in the Iron oxide growth assay section. Following the mineralization reaction, samples were transferred into glass capillaries and measured at 4 °C with an acquisition time of 30 min per sample. Corresponding reaction buffer samples were measured under identical conditions and used for background subtraction. Data reduction, background subtraction, and analysis were performed using RAW software. Where indicated, the linear low-q region was fitted with a power-law function to determine the scattering exponent.

### Atomic force microscopy (AFM)

Solutions of the Z0-fiber, Z3, and Z4 designed proteins as well as BSA and lysozyme controls were prepared at a concentration of 1 nM in 20 mM HEPES buffer (pH 8) containing 50 mM NaCl and 1 µM Zn(NO_3_)_2_. Commercialized ZnO (0001) crystals (MTI Corp., CA) were cut into pieces measuring 3 × 3 × 0.5 mm. The crystal surfaces were sonicated in DI water for 15 minutes, followed by UV-ozone treatment for 15 minutes. The cleanliness of each ZnO crystal surface was checked in DI water using AFM before the protein-binding measurement. Then, the ZnO crystal was dried and incubated in the protein solution for 5 minutes. AFM measurements were conducted in liquid immediately after incubation to assess protein binding.

All AFM measurements were done with a MultiMode VIII AFM (Bruker, CA) in PeakForce tapping mode, employing an SNL-10-B probe (Bruker, CA). The imaging buffer consisted of 20 mM HEPES (pH 8), 50 mM NaCl, and 1 µM Zn(NO₃)₂. The offline data processing was performed using Nanoscope Analysis 2.0 (Bruker, CA) and ImageJ (NIH).

### Protein size and density analysis

To identify proteins observed on the crystal surface with AFM, ImageJ with the Labkit plugin was employed for size and morphology-based identification. Although ZnO(001) is chemically stable at pH 8, its surface slowly forms Zn-hydroxide species (*50*) giving rise to large clusters distinguishable from proteins by size. Protein molecules, Zn-hydroxide clusters, and ZnO background were labeled green, red, and blue, respectively (**fig. S13B**). Protein coverage was determined by dividing the total pixel count of protein molecules (green) by the total pixel (512 by 512 pixels, each pixel equals 0.608 nm^2^) of the AFM images. Four images were analyzed for each protein for the accuracy of protein coverage measurement. The protein size was analyzed by calculating the pixel number of each green block, followed by a statistical analysis to determine the average size, statistically. For protein binding density, the coverage is converted to the protein binding area in µm^2^, and then divided by the average protein size.

### Pulldown assays

100 μL samples containing 0.7 mg/mL protein in 150 mM NaCl 20 mM Tris-HCl pH 8 buffer (TBS) were prepared for designed ZnO binding protein as well as bovine serum albumin and lysozyme controls and placed in strip PCR tubes. 5 μL of suspensions containing 2% w/v of anatase (TiO_2_), rutile (TiO_2_), zinc oxide (ZnO), or hematite (Fe_2_O_3_) nanoparticles were added and the samples were incubated at room temperature while shaking at 1000 rpm in a plate shaker for 30 minutes. Samples were spun at 5000 relative centrifugal force (rcf) for 5 minutes, after which the supernatant was carefully removed so as to not disturb the pelleted inorganic material. The pellets were thoroughly washed by respending in 100 μL TBS with pipetting and vortexing, incubating while shaking at 1000 rpm for 5 minutes, and spinning at 5000 rcf for 5 minutes. This process was repeated for a total of three TBS washes, after which the final pellet was resuspended in 100 uL sodium dodecyl sulfate-polyacrylamide gel electrophoresis (SDS-PAGE) x2 loading buffer (Bio-Rad) and incubated at 98°C in a thermocycler for 30 minutes. 7 μL of the samples were loaded into Mini-PROTEAN® TGX™ Precast Gels (Bio-Rad) and run under electrophoresis at 180 volts with either Kaleidoscope or Dual Xtra prestained standard ladders (Bio-Rad). The resulting gels were Coomassie stained with an eStain™ machine (GenScript) and photographed under visible light with a ChemiDoc™ XRS imager (GenScript).

### Protein ZnO nucleation photoluminescence assay

Protein samples, taken from stock solutions ranging from 0.25 to 1.2 mg/ml concentration and suspended in a buffer solution consisting of 100 mM HEPES and 50 mM NaCl at pH 8.2, were combined with 30 mM Zn(NO_3_)_2_ in a 1:10 ratio, resulting in a final Zn^2+^ concentration of 3 mM and final protein concentration of 0.1 mg/mL. All ZnO growth solutions contain 3 mM Zn(NO_3_)_2_, 50 mM NaCl, 100 mM HEPES pH 8.2, and 0.1 mg/mL protein or 12.5 μg/mL ZnO NP as indicated. This mixture was then incubated at 25°C for 2 hours within a Synergy NEO2 plate reader. Kinetic measurements of ZnO photoluminescence (PL) were conducted at 5-minute intervals using excitation at 325 nm and emission wavelengths of 600 nm. For time-series plots, the replicate with the highest peak PL signal is shown rather than the mean, as stochastic variation in nucleation onset makes averaging uninformative. All six replicates are quantified in the accompanying box plots. All replicates in this work are technical replicates, independent reactions prepared from a common protein stock, unless otherwise stated. Nucleation time was defined as the time at which the PL signal reached the halfway point between its initial value and maximum value. Only replicates with a maximum PL signal greater than 100 (arbitrary units) were considered to have undergone nucleation and are included in nucleation time analyses.

### Iron oxide growth assay

To evaluate the impact of various proteins on hematite formation, we established conditions that strongly favor the formation of magnetite (Fe_3_O_4_) over hematite (Fe_2_O_3_). We thought that the ferrimagnetic properties of magnetite would enable real-time monitoring of material assembly from a supersaturated solution. To detect magnetite formation, we designed and 3D-printed a custom cover for a 96-well plate, positioning a magnet at the center of each well (**fig. S23**). This localized magnetic field allowed us to distinguish between centrally aggregated, ferrimagnetic material and peripheral, non-ferrimagnetic deposits. Experiments were conducted within a glovebox to eliminate oxygen-induced biases. The assay was recorded using a benchtop scanner configured to capture a color image every 30 seconds (with an additional 30-second acquisition time per frame). To induce the formation of magnetite/hematite, seed samples were prepared by adding 100 μL of a solution containing either H7-neg, H9-neg, or lysozyme (final concentration 0.15 mg/mL), buffer alone, or hematite/magnetite nanoparticles (final Fe concentration 0.2 mM) to six separate wells. These samples were suspended in a purged buffer containing 100 mM CHES and 50 mM KNO₃ (final pH 8.7). Iron oxide formation was initiated by sequentially adding Iron(III) nitrate nonahydrate (Millipore Sigma, Cat. 254223) to a final concentration of 1.0 mM, followed by ammonium iron(II) sulfate hexahydrate (Millipore Sigma, Cat. 09719) to a final concentration of 0.5 mM, and finally adjusting the total volume to 200 μL with purged Milli-Q water. The wells were then covered with the magnetic holder and placed on the scanner for time-lapse data collection (Canon LiDe 110).

### Colorimetric quantification

Quantitative analysis was performed by comparing the red channel intensity in the scans to the total intensity (RGB) within a defined ring across each frame (**fig. S23, and movie S1**). Normalized red intensity was calculated for each well as the mean red channel intensity divided by the sum of mean red, green, and blue channel intensities within the defined ring region. Mean normalized red intensities (n = 6) are plotted in **Fig. 3M**, and **fig. S24**.

### Rust inhibition assay

To evaluate the ability of the identified hematite designs to inhibit rust formation on exposed iron surfaces, iron substrates were first polished to achieve a uniform surface finish, followed by sequential washing with Milli-Q water and ethanol (three times each). To ensure consistent application of the sample solutions, a plastic stencil cover tape with uniformly sized holes was fixed over the iron surface. 10 µL of H7-neg, lysozyme (0.6 mg/mL), and buffer (100 mM CHES, 50 mM KNO₃, pH 8.7) were then applied to the exposed areas and incubated for 2 hours. Following incubation, the treated surfaces were covered with plastic tape and scanned using a desktop office scanner (HP PageWide Pro MFP 477dn) to document the results.

### Negative stain transmission electron microscopy (ns-TEM)

3 μl of protein samples at 0.01 mg / mL was applied to plasma discharged carbon coated copper grids (01844 TedPella) and incubated for 2 minutes. The grid was blotted with filter paper, touched to a drop of 40 uL Milli-Q water, blotted, and stained by applying and blotting three droplets of 2% uranyl formate, with the last being incubated for 2 minutes before blotting.

A FEI Talos L120C TEM (Thermo Scientific) with a 4K × 4K Gatan camera was used at 57kx magnification and 2.49 Å pixels. Thermo Scientific EPU software was used to automate data collection and CryoSPARC was used to process data (*51*). 300 particles were manually selected to generate 2D classes for subsequent template picking. Template picked particles were used to generate 2D classes and some were selected for ab initio 3D reconstruction with C1 symmetry. Non-uniform refinement with the observed symmetries produced the final models.

### Visualization and figures preparation

Designs and structural images were produced with either PyMOL or ChimeraX. Experimental data was analyzed and plotted using python MatplotLib and Seaborn libraries. Figures for the manuscripts were rendered using Adobe Illustrator and Inkscape.

### X-Ray crystallography

Crystallization experiments for this project were conducted using the sitting drop vapor diffusion method. Crystallization trials were set up in 200 nL drops using the 96-well plate format at 20°C.

Crystallization plates were set up using a Mosquito LCP from SPT Labtech, then imaged using UVEX microscopes from JAN Scientific. Diffraction quality crystals of Z4-C3i formed in 0.2M Sodium chloride, 0.1M Sodium acetate pH 4.6, and 30% v/v (+/-)-2-Methyl-2,4-pentanediol and Z4-C3ii crystals formed in 0.03M Sodium fluoride, 0.03M Sodium bromide, 0.03 Sodium iodide, 0.0501M MOPS, 0.0499M Sodium HEPES, 12.5% v/v MPD, 12.5% w/v PEG 1000, 12.5% w/v PEG 3350.

Diffraction data were collected at the National Synchrotron Light Source II at beamline 17ID-1. Z4-C3i diffracted to 3.06 Å resolution and Z4-C3ii to 2.99 Å. X-ray intensities and data reduction were evaluated and integrated using XDS (*52*) and merged/scaled using Pointless/Aimless in the CCP4 program suite (*53*). Structure determination and refinement starting phases were obtained by molecular replacement using Phaser (*54*) using the designed model structure. Following molecular replacement, the models were improved using PHENIX (*55*) and manually built in Coot (*56*), and refined in Phenix. Model building was performed using Coot. The final model was evaluated using MolProbity (*57*). Data collection and refinement statistics are recorded in **table S10**. Data deposition, atomic coordinates, and structure factors reported for in this paper have been deposited in the Protein Data Bank (PDB), http://www.rcsb.org/ with accession codes pdb_00009D92 and pdb_000037TT.

### CryoTEM Z4-C6

Cryo-EM sample preparation: Cryo-EM grids were prepared by diluting protein samples with TBS 1 to 5 times immediately before applying 3.5 μL to glow-discharged 400 mesh, C-flat, 2 micron holes, 2 micron spacing, CF-2/2-4C (CF-224C-100) (Electron Microscopy Sciences, Hatfield, PA) cryo-EM grids. Grids were blotted using a blot force of 0 and 5.5 second blot time at 100% humidity and 4°C and plunge-frozen in liquid ethane using a Vitrobot Mark IV (FEI Thermo Scientific, Hillsboro, OR).

Cryo-EM data collection and processing: cryo-EM grids were screened and data was collected on a Krios transmission electron microscope (FEI Thermo Scientific, Hillsboro, OR) operated at 300 kV and equipped with a Gatan K3 Summit direct detector. Data collection was performed automatically using SerialEM. Movies were acquired in super-resolution mode at 0.4124 Å per pixel at a nominal magnification of 105,000, 40 frames, at a dose rate of 30(e-/Å/s), for a total dose of 50(e-/Å^2^) and a defocus range of 0.8 - 2.0 (µm).

All data processing was carried out in CryoSPARC (*51*). Alignment of movie frames was performed using Patch Motion with an estimated B-factor of 500 Å^2^, with a maximum alignment resolution set to 3. Defocus and astigmatism values were estimated using Patch CTF with default parameters. 119 particles were initially manually picked and averaged into 20 2D classes. Classes of particles clearly containing ZnO crystals and classes of crystal free particles were selected and divided into two, followed by multiple rounds of 2D classification and subsequent template-picking and extracted with a box size of 400 pixels. The processing pipeline for this design is illustrated in **fig. S31**.

### Z4-Cage cryo-EM sample preparation

1 mg/mL Z4-cage following 60 minutes of co-incubation with supersaturated precursor solution (3 mM Zn(NO3)2, 50 mM NaCl, 100 mM HEPES, pH 8.2) was applied to glow-discharged Quantifoil R 2/2 300 mesh copper grids overlaid with an additional thin layer of carbon. Vitrification was performed on a Mark IV Vitrobot at 22°C at 100% humidity, with a wait time of 7.5 seconds, a blot time of 6 seconds, and a blot force of 0 before being immediately plunged frozen into liquid ethane. The sample grids were clipped following standard protocols before being loaded into a ThermoFisher Glacios 200 kV transmission electron microscope for imaging.

### Z4-Cage cryo-EM data collection

Z4-Cage data were collected automatically using SerialEM (*58*) and used to control a ThermoFisher Glacios 200 kV TEM equipped with a standalone K3 Summit direct electron detector and operating in counting mode. Random defocus ranges spanned between −0.8 and −1.8 μm using stage move, with one-shot per hole and a single hole per stage move. Altogether, 4,018 movies were recorded with a pixel size of 0.885 Å with a total dose of 50 e^−^/Å^2^.

### Z4-Cage cryo-EM data processing

All data processing was performed in either CryoSPARC LIVE or CryoSPARC (*51*, *59*). The video frames were aligned using Patch Motion with an estimated B factor of 500 Å^2^. The maximum alignment resolution was set to 5. Defocus and astigmatism values were estimated using the patch CTF estimation with the amplitude contrast set to 0.07. For the Z4-cage, data was initially processed in CryoSPARC Live (*60*) with blob picking parameters set to a particle size range of 160–200 Å and an extraction box size of 480 pixels. Approximately 225,000 particles were initially picked and subjected to 2D classification. From this, the most well-defined 40,134 particles were selected as templates for a second round of template-based particle picking. Particles were then re-extracted with a 480-pixel box size, followed by another round of 2D classification into 100 classes. This classification employed three iterations, 50 online-EM iterations, and a batch size of 200 particles per class. The top 608,979 particles from this round were exported to CryoSPARC for further processing.

Using a subset of ∼40,000 particles with C1 symmetry, a 3D ab initio reconstruction was performed to generate three initial models. This yielded maps of well-formed octahedral nanocages in multiple classes. The full set of 608,961 template-picked particles was then subjected to 2D classification, with the best 551,868 particles selected for high-resolution homogeneous refinement under octahedral symmetry constraints. Despite achieving a reported global resolution estimate of ∼3.3 Å, the map quality was unexpectedly poor, with poorly defined helices and a lack of structural details characteristic of this resolution. This was suspected to result from the presence of ZnO around many cages.

To address this, refinement protocols incorporating symmetry relaxation (using maximization-based relaxation) were employed. This approach produced a map with a reported resolution of 3.83 Å, slightly lower than the previous estimate but revealing significantly improved structural detail consistent with protein features. Visualization of the ZnO density indicated its presence along the C3 axes but not the C4 axes. Subsequently, the refinement was repeated without applying symmetry constraints, yielding a map with a reported resolution of 3.87 Å. Low-pass filtering this map to 8 Å and adjusting contour levels confirmed the presence of ZnO localized exclusively at the designed C3 interface.

To better resolve ZnO density at the C3 trimer, octahedral symmetry expansion was performed, focusing on the Z4-cage’s C3 trimer. Subsequent 3D classification identified classes with either filled or empty C3 interiors. Multiple rounds of 3D classification were conducted to improve the ZnO density quality further. Ultimately, 3D local refinement under C1 symmetry, applied to the best class containing a well-resolved and filled Z4-C3i trimer, yielded a final map with a reported global resolution estimate of 3.34 Å. This refinement utilized 1,332,943 symmetry-expanded and 3D-classified particles.

### Z4-Cage cryo-EM model building and validation

For the structure of the Z4-Cage, the model was initially built using the predicted computational design model. Following rigid-body docking of the design model into the cryo-EM density map, the structure was relaxed into density in ChimeraX (*61*) using ISOLDE (*62*). Side chains were trimmed to their Cβ carbons due to the absence of discernible side-chain features at the achieved resolution. For the symmetry-expanded, 3D-classified, and locally refined Z4-C3i trimeric structure, a similar process was employed. However, instead of trimming side chains after ISOLDE refinement, all atomic positions were manually inspected and adjusted using Coot (*56*).

The atomic density corresponding to potential H_2_O molecules or ions along the designed ZnO binding interface was examined manually. We modeled this density as water molecules comparing multiple high-resolution maps generated from the Z4-C3i trimer dataset. This juxtaposition allowed differentiation between real density features and noise. Following the manual placement of H_2_O molecules, the structure underwent ISOLDE refinement, with water coordinates further optimized using density-guided molecular dynamics flexible fitting (MDFF). Placed waters that did not form at least one hydrogen bond or failed to align with the center of the corresponding density were manually removed. This process was iteratively repeated until convergence was achieved, with high agreement between the structure, density map, and known atomic principles.

Residues with poor density were truncated to their corresponding Cβ carbons. Phenix real-space refinement was subsequently performed as a final step before the final model quality was analyzed using MolProbity (*57*). Figures were generated using either UCSF Chimera (*63*) or UCSF ChimeraX (*61*). The final structures of the Z4-cage and the Z4-C3i trimer structure within the cage were deposited in the PDB under accession numbers pdb_00009O59 and pdb_00009O5Q respectively, and the EMDB under EMD-70146 and EMD-70128 respectively.

## Acknowledgments

We thank J. M. Galloway and F. Praetorius for helpful discussions.

## Author contributions

Conceptualization: AS, HP, FADH, DB. Data curation: AS, HP, AJB, AKB, NH. Formal analysis: HP, AS, AJB, AKB, NH. Funding acquisition: LS, JJDY, DB. Investigation: AS, HP, EBH, PSK, AJB, WD, SYS, XL, SZ, AKB, AK, EB, AC, KDC, AA, DKV, YB, NH. Methodology: AS, HP, EBH, PSK. Resources: DKV, AA, RDK, LG. Software: HP, EBH, PSK. Supervision: DB, JJDY, SZ. Visualization: HP, AJB, AS, NH. Writing – original draft: HP, AS, DB. Writing – review & editing: All authors.

## Data, code, and materials availability

All data are available in the main text or the supplementary materials. Source data for all plotted panels in the main text and supplementary figures are provided in Data S1. All protein design models and scripts are available at https://github.com/pylesharley/protein_mineral_library and archived at Zenodo (*41*). The crystal structures of Z4-C3i and Z4-C3ii are available in the RCSB Protein Data Bank (ids: pdb_00009D92 and pdb_000037TT). Cryo-EM structures of the Z4-cage and the Z4-C3i trimer structure within the cage are available in the RCSB Protein Data Bank (ids: pdb_00009O59 and pdb_00009O5Q) and EMDB (ids: EMD-70146 and EMD-70128). The Rosetta Macromolecular Modelling suite is available for non-commercial use at https://www.rosettacommons.org (*40*). Oligo pool overlap optimization used OligoOverlapOpt archived at Zenodo (*44*). All materials described in this work are available from the corresponding author upon reasonable request.

## License Information

Copyright © 2026 the authors, some rights reserved; exclusive licensee American Association for the Advancement of Science. No claim to original US government works. https://www.science.org/about/science-licenses-journal-article-reuse. This article is subject to HHMI’s Open Access to Publications policy. HHMI lab heads have previously granted a nonexclusive CC BY 4.0 license to the public and a sublicensable license to HHMI in their research articles. Pursuant to those licenses, the Author Accepted Manuscript (AAM) of this article can be made freely available under a CC BY 4.0 license immediately upon publication.

### Supplementary Materials

Supplementary Text

MDAR Reproducibility Checklist

Figs. S1 to S35

Tables S1 to S11

Movie S1

Data S1

