## Supplementary materials for "Programmed mineralization of metal-oxides by de novo designed protein templates"

**The PDF file includes:**

Supplementary Text
Figs. S1 to S35
Tables S1 to S11
Caption for Movie S1
Caption for Data S1

**Other Supplementary Materials for this manuscript include the following:**

Movie S1
Data S1

### Supplementary Text

#### Biochemical characterization of metal-oxide binding proteins

Of the five well-expressing and soluble ZnO binding proteins, Z3 and Z5 appeared monomeric by size exclusion chromatography (SEC), while Z0, Z1, and Z4 appeared to form large assemblies (**fig. S11**). Despite having surface exposed cysteine residues, Z3 remained monomeric in solution (**fig. S11**). Z0, Z3, Z4, and Z5 produced circular dichroism (CD) spectra consistent with their predicted secondary structure, while Z1 appeared disordered (**fig. S11**). Z0 formed long fibers and bundles of fibers visible with negative-stain transmission electron microscopy (ns-TEM), presumably through strand pairing head-to-tail interactions (**fig. S14A**). To regularize these interactions, we designed and produced a version without these capping features, Z0-fiber, and used it in further analysis of this design (**fig. S14B**). In a pull-down assay followed by SDS-PAGE, all of the purified ZnO designs bound ZnO NPs and did not bind to anatase ($TiO_2$), rutile ($TiO_2$), or hematite NPs, while bovine serum albumin (BSA) and lysozyme controls showed no binding (**fig. S12**). Atomic force microscopy (AFM) of the zinc-terminated (001) surface incubated with the proteins showed binders had similar (Z3, Z4) or greater (Z0-fiber) coverage in respect to controls (**fig. S13**). The three designs which produced the highest yields of purified protein, Z0-fiber, Z3, and Z4, were selected for further study.

We characterized two hematite binders, H7 and H9, which expressed as soluble protein in *E. coli* and demonstrated robust hematite NP binding signals when displayed as single clones on yeast (**Fig. 2I, fig. S8, and table S2).** In order to increase the yield of solubly expressed protein and improve the monodispersity of these proteins we mutated non-interfacial positively-charged residues to neutral and negatively-charged residues (*34*), resulting in H7-neg and H9-neg (**figs. S15 and S16, and table S8**). Hematite nanoparticles incubated with H7-neg showed decreased flocculation compared to solutions without protein or BSA (**fig. S17**).

#### Summary of design topologies

Designs based on DHR, DBS, and mixed alpha-beta topologies bound target NP when displayed on yeast (**figs. S7 to S10**), but DHRs were the most likely to express as soluble protein in *E. coli* (**tables S1 and S2)** and the most amenable to designing into assemblies (**Fig. 4**). DHR designs with poor solubility could be rescued by introducing directed mutations away from the designed inorganic-binding interface to increase the net negative charge (**fig. S16 and table S2)**.

#### Details of $Zn^{2+}$ ion binding

A ratiometric dye (Mag-Fura-2) assay indicated Z0-fiber binds $Zn^{2+}$ ions with a best fit model indicating six binding sites, three with $3.8 \pm 0.2$ nM affinity and three with $60 \pm 6$ nM affinity (**Fig. 2D**), and Z3 showed three binding sites with $200 \pm 60$ nM affinity (**Fig. 2E;** see Methods). The difference in measured affinities is likely due to the lower pH for the ITC measurements (see Methods).

#### Summary of zinc oxide binding mechanisms

We identified thirteen de novo ZnO binding protein interfaces (**fig. S9**) and demonstrate that three of them, Z0-fiber, Z3, and Z4, promote ZnO growth (**Fig. 3, B to D, and fig. S9)**. Our results suggest these interfaces function by two distinct mechanisms. Firstly, there is evidence for direct ion-mediated interactions; Z0-fiber and Z3 coordinate $Zn^{2+}$ ions and mutating metal-coordinating residues reduces their affinity to both ions and ZnO NPs (**Fig. 2, D, E, and G, and fig. S18**). Secondly, ZnO binding interfaces, including Z4, contain two types of threonine motifs observed in both alpha-helical and beta-sheet ice-binding proteins (**Fig. 2, F and H, and fig. S9**), suggesting they may interact with ZnO via ordered water layers, similar to proposed mechanisms of ice binding by ice-binding proteins (*16*, *17*, *64*). 3D local refinement of the cryo-EM data at the threefold axis of the mineralized Z4-cage (2.5 Å at the center of the C3 axis, ~3.5 Å at the inter-trimer interface) showed density near the designed interface that could be explained by water organization (**Fig. 5H, fig. S34, and table S11**). Sixty-one waters were modeled into this density (see Methods), though their identity as water rather than $Zn^{2+}$, $OH^{-}$, or another species could not be confirmed at this resolution. The lack of a detectable $Zn^{2+}$ ion binding signal by the Z4 interface (**Fig. 2G**, **fig. S19**) argues against assigning this density to $Zn^{2+}$ ions. The same Z4 interface in the Z4-C3i crystal structure also showed density consistent with ordered waters near its threonine motifs (**fig. S29**), but the resolution was insufficient to place waters at most positions. The positions of the density near the Z4 interfaces were different in the mineralized Z4-cage cryo-EM structure and unmineralized Z4-C3i crystal structure, which is unsurprising given the different conditions: the Z4-cage was incubated in solution supersaturated for ZnO (3 mM $Zn(NO_3)_2$, 50 mM NaCl, 100 mM HEPES, pH 8.2), whereas Z4-C3i was crystallized in 0.2 M NaCl, 0.1 M sodium acetate pH 4.6, 30% v/v MPD.

Hydrophobic interfaces were also enriched in our assay, but none of these proteins could be purified as soluble protein from *E. coli* so the mechanism of their binding, and whether or not they template ZnO, is unclear (**tables S1 and S2**). While periodic charges were present in interfaces alongside other features (**Fig. 2 and tables S1 and S2**), no interfaces composed entirely of arrays of charged groups, like those seen in previous mineral-templating and mineral-binding designed proteins (*1*, *14*), were enriched in our library sorting. ZnO NP binding was more predictive of ZnO nucleation activity than binding to a bulk zinc-terminated ZnO (001) surface (**Fig. 3 and figs. S12 and S13**).

**Summary of hematite binding mechanisms**

We identified seven surfaces that promote hematite binding (**fig. S10**) and demonstrate that two proteins, H7-neg and H9-neg, direct the formation of hematite in conditions that otherwise produce magnetite (**Fig. 3, L to N**), and appear to reduce the rate of oxidation at iron-water interfaces (**fig. S27**). These proteins coordinate $Co^{2+}$, an established proxy for $Fe^{2+}$ (**Fig. 2I**), suggesting they function through metal-mediated interactions.

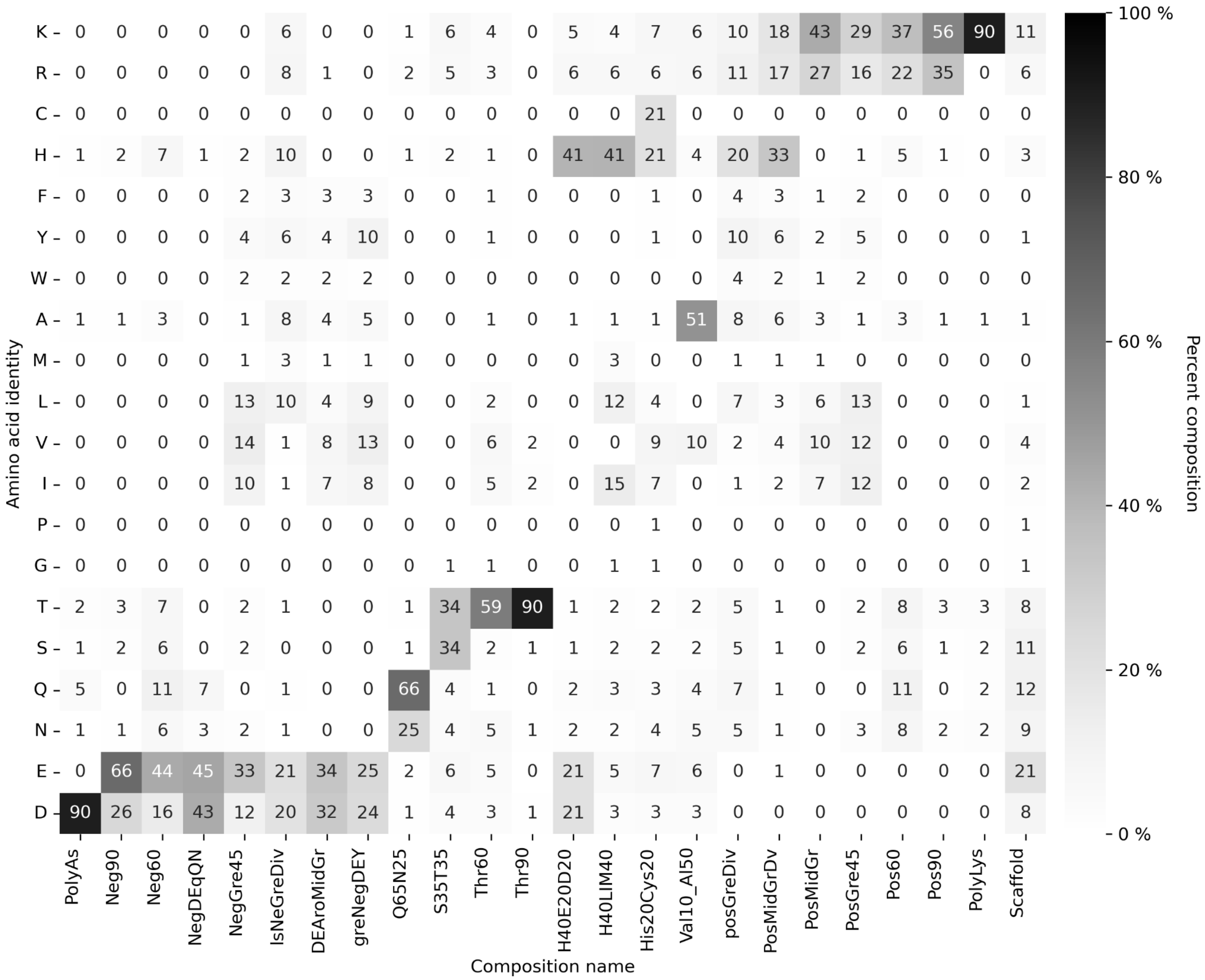


**Fig. S1. Library design, amino acid compositions vs. frequencies.** Frequency heat-map showing the mean percentage of each amino acid (Y-axes) in the defined compositions (X-axes) used to decorate the scaffold proteins to generate the metal oxide binders library. Rosetta amino-acid constraint files to enforce these compositions are available in the repository associated with this publication (*40*).

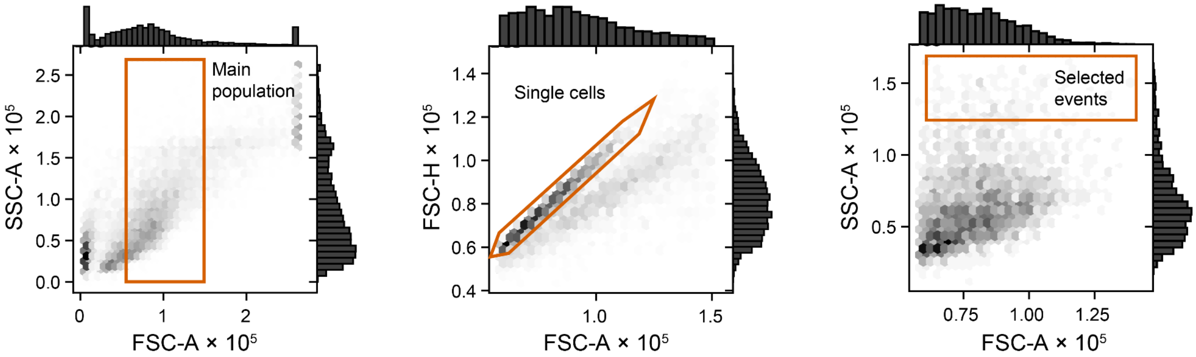


**Fig. S2. Mineral sort gating strategy.** Representative two dimensional density plots describing the strategy applied to sort metal oxide nanoparticle binding yeast clones. (**Left**) Main population is gated to exclude aggregates, dead cells and free particles. (**Center**) Singlet (single-cell) events were enriched by plotting forward scatter area (FSC-A) versus forward scatter height (FSC-H) and applying a narrow linear gate along the central diagonal of the population, thereby excluding doublets and larger aggregates. (**Right**) A sort gate capturing less than 1% of control events is set to capture binding clones.

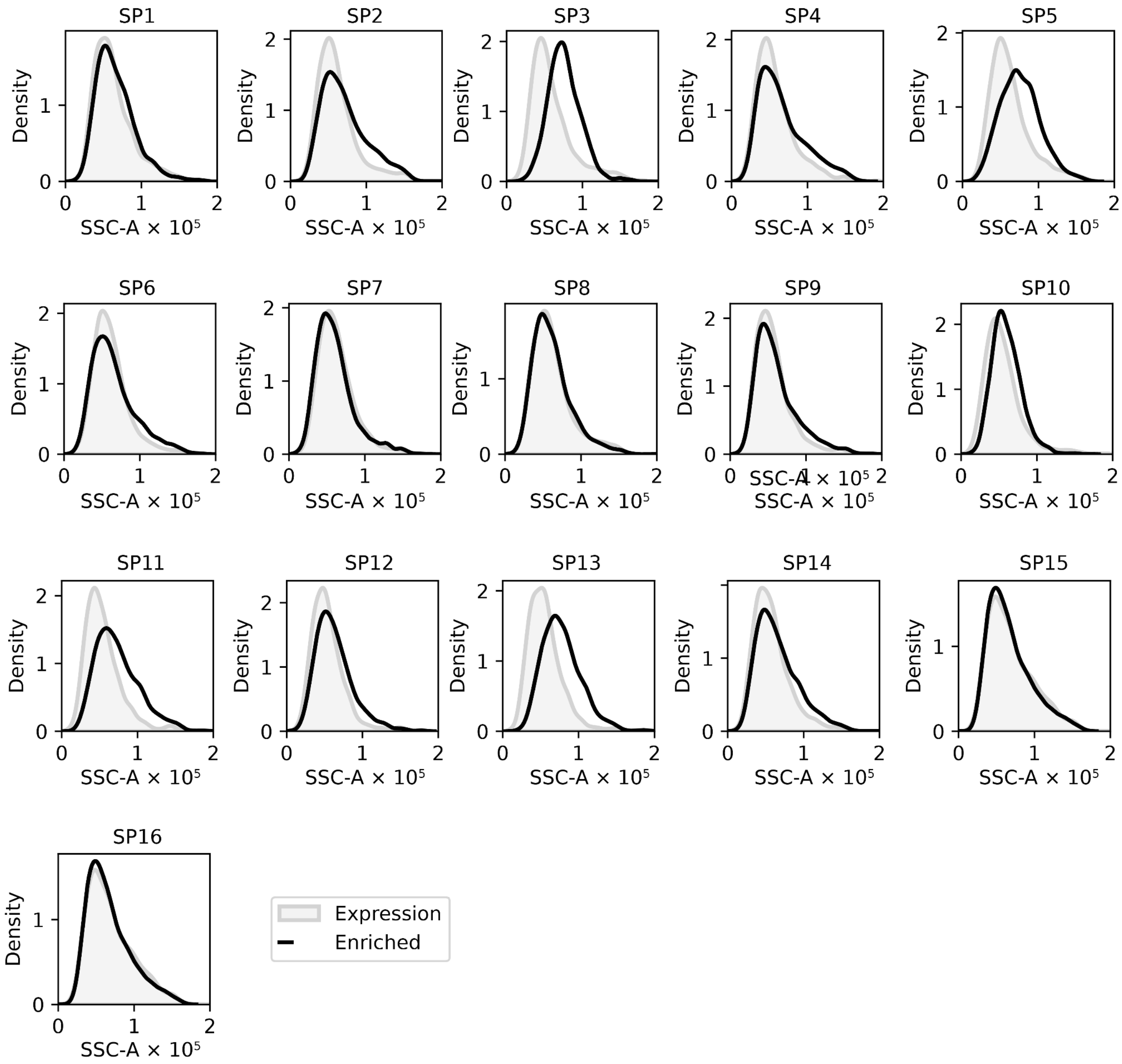


**Fig. S3. ZnO associated side scattering signal in enriched subpools, in comparison to initial expression subpools.** Histograms plotting the SSC signal of individual subpools, enriched by 3 consecutive ZnO sorts (black line) vs. expression, subpools sorted for expression only (filled light gray).

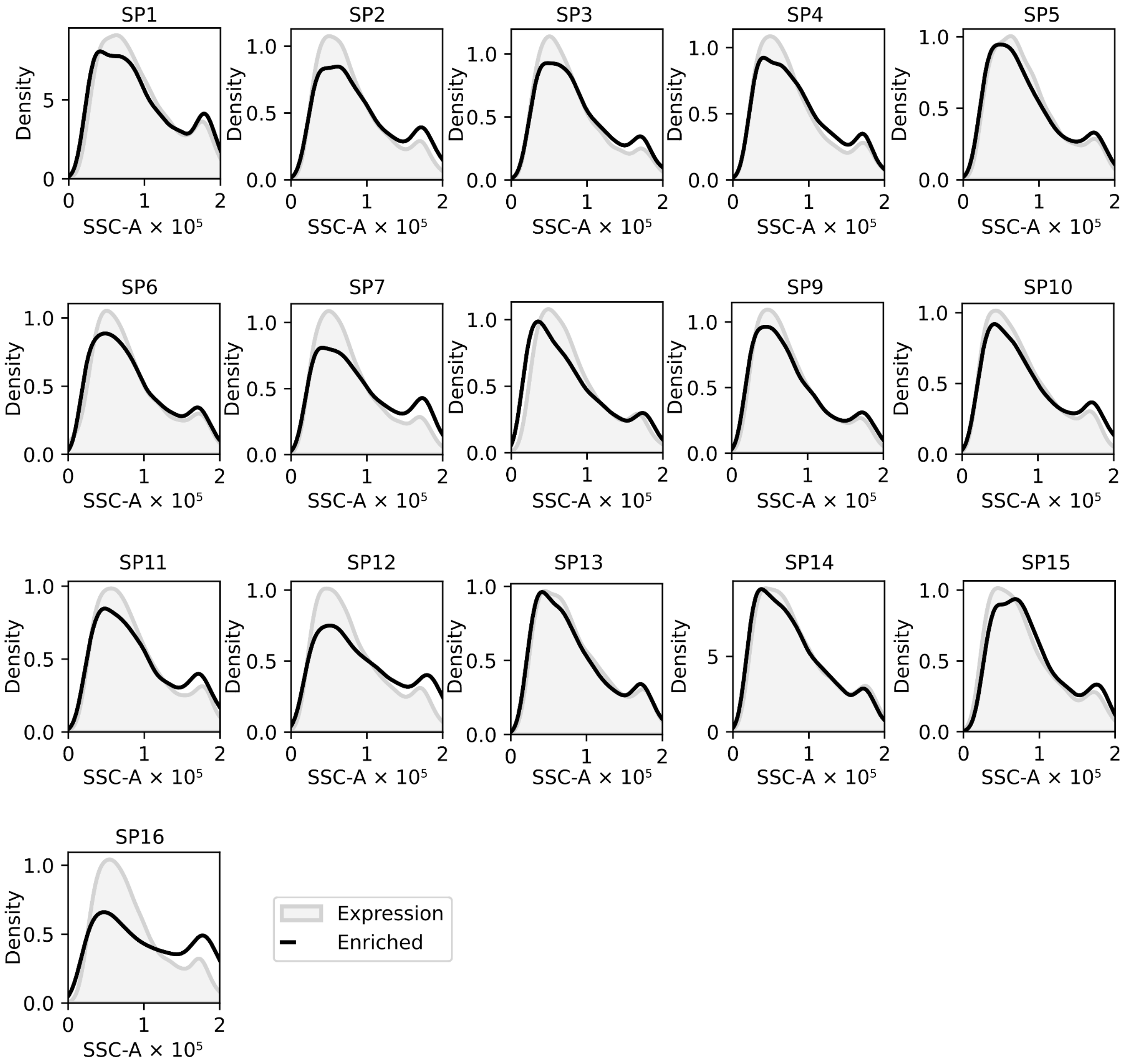


**Fig. S4. Hematite (Fe2O3) associated side scattering signal in enriched subpools, in comparison to initial expression subpools.** Histograms plotting the SSC signal of individual subpools, enriched by 3 consecutive hematite sorts (black line) vs. expression, subpools sorted for expression only (filled light gray).

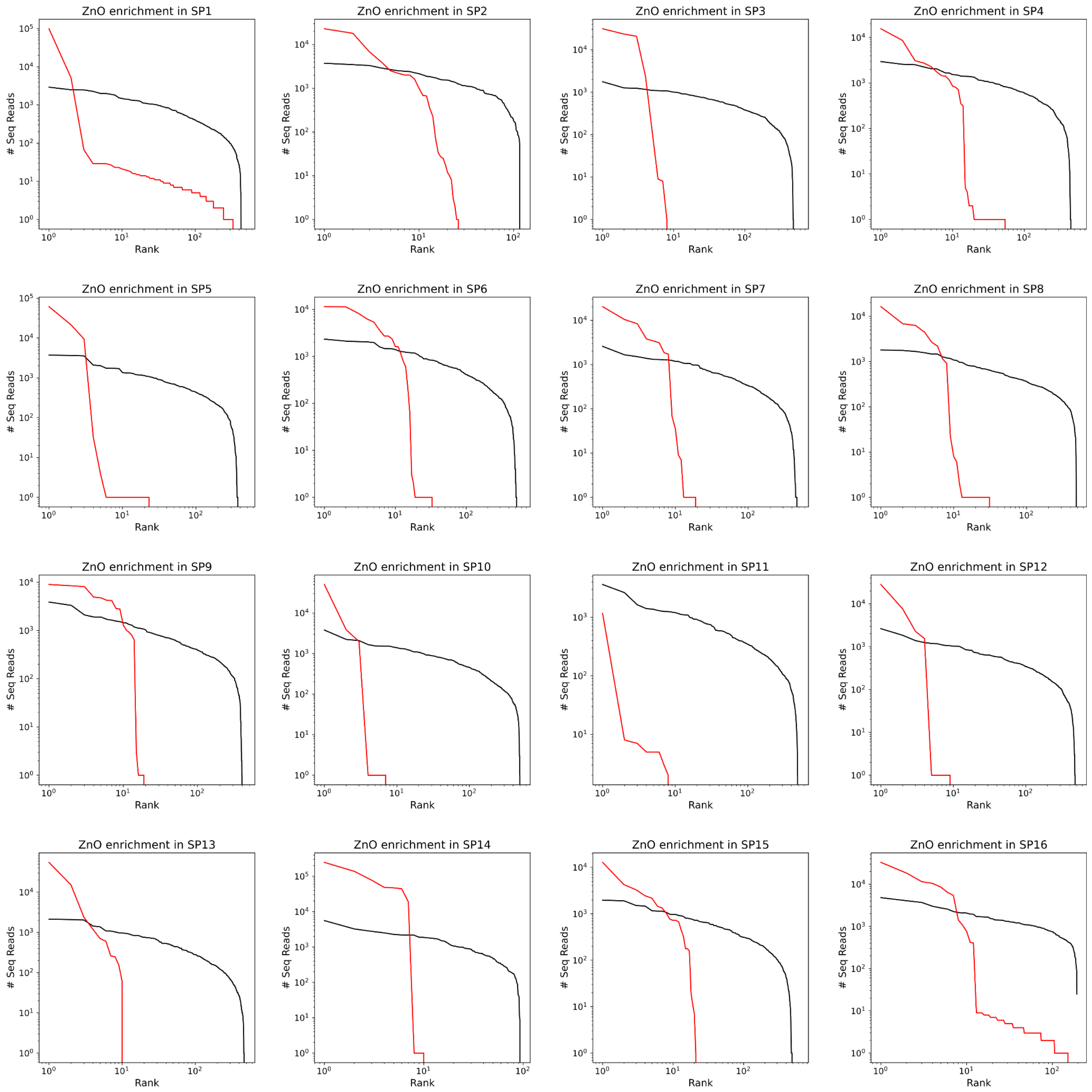


**Fig. S5. NGS enrichment plots.** NGS enrichment plots displaying individual library subpools enriched by sorting for ZnO binders (red line) vs. expression (black line). Individual sequences are plotted as a function of their rank (X-axes) and sequence count (Y-axes).

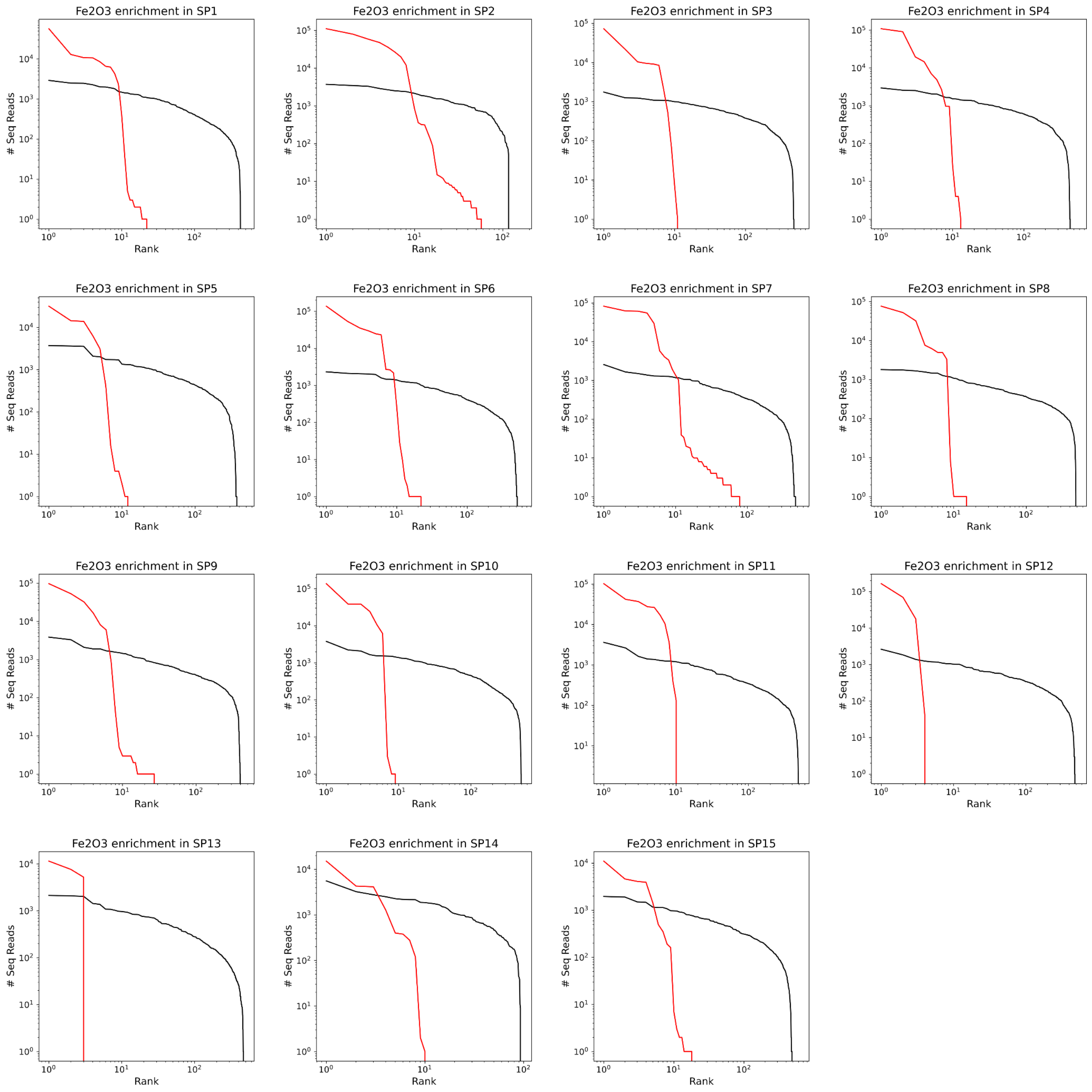


**Fig. S6. NGS enrichment plots for Hematite (Fe2O3).** NGS enrichment plots displaying individual library subpools enriched by sorting for hematite binders (red line) vs. expression (black line). Individual sequences are plotted as a function of their rank (X-axes) and sequence count (Y-axes).

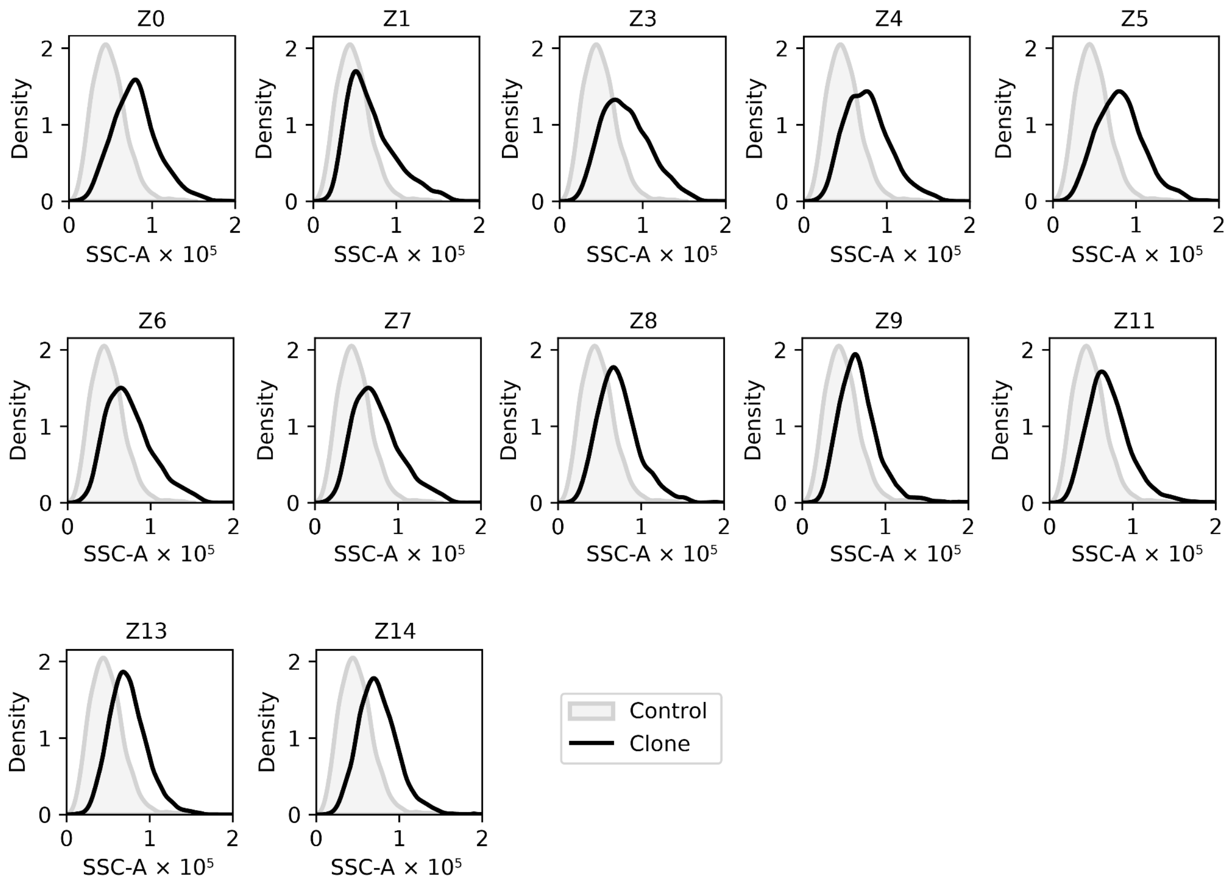


**Fig. S7. ZnO associated SSC signal in identified single clones vs. control.** Histograms plotting the SSC signal of NGS identified clones (black line) vs. control yeast expressing a mock sequence (filled light gray). Clone Z12 expressed poorly on the surface of yeast and was excluded from the assay.

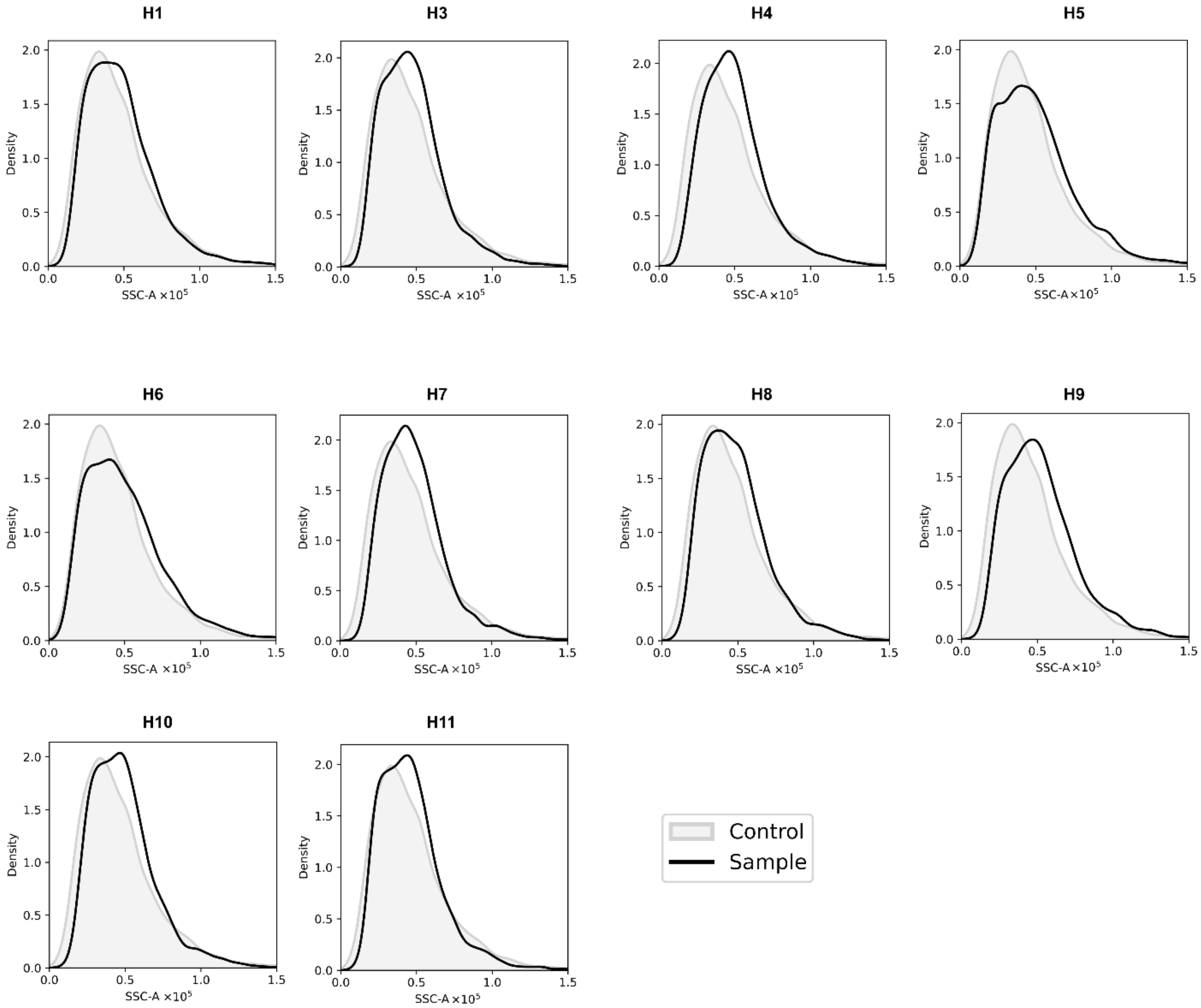


**Fig. S8. Hematite associated SSC signal in identified single clones vs. control.** Histograms plotting the SSC signal of NGS identified clones (black line) vs. control yeast expressing a mock sequence (filled light gray). Clone H2 expressed poorly on the surface of yeast and was excluded from the assay.

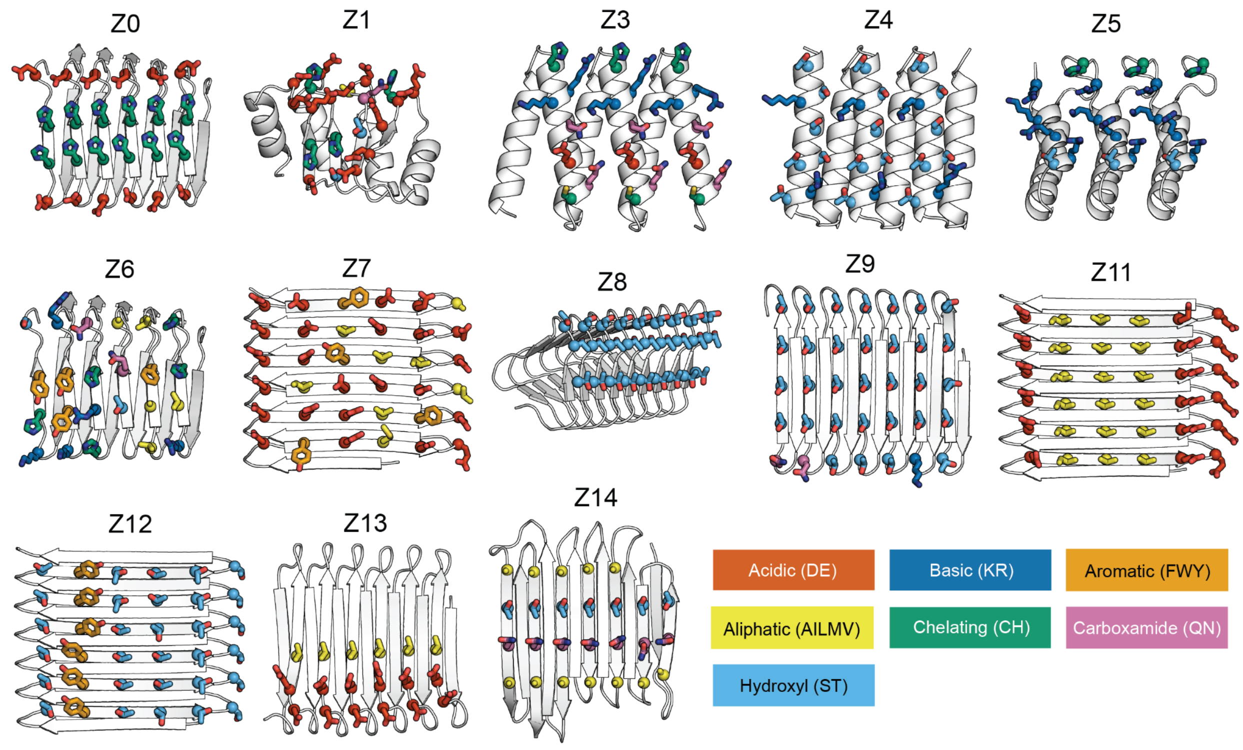


**Fig. S9. Models of interfaces enriched for ZnO binding**. Colors of amino acid side chains correspond to the chemical categories identified in rectangles.

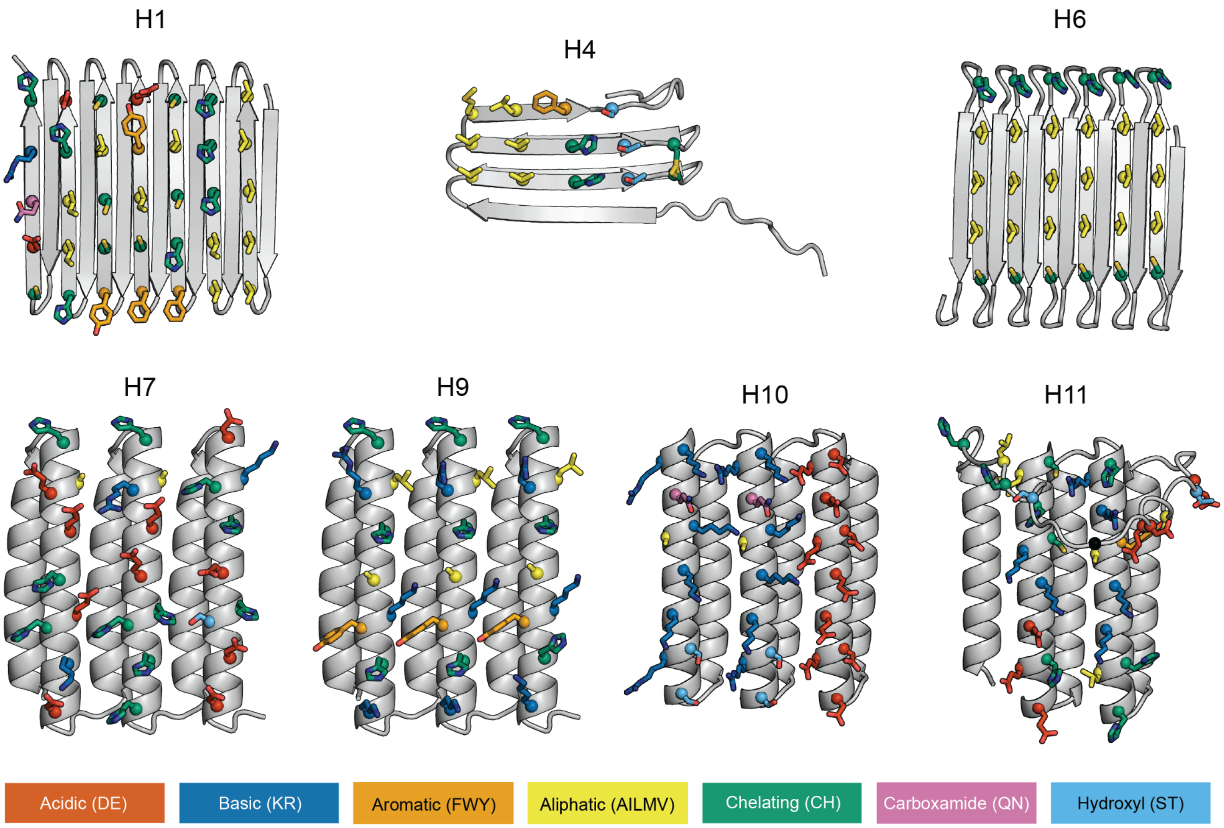


**Fig. S10. Models of interfaces enriched for hematite binding**. Colors of amino acid side chains correspond to the chemical categories identified in rectangles.

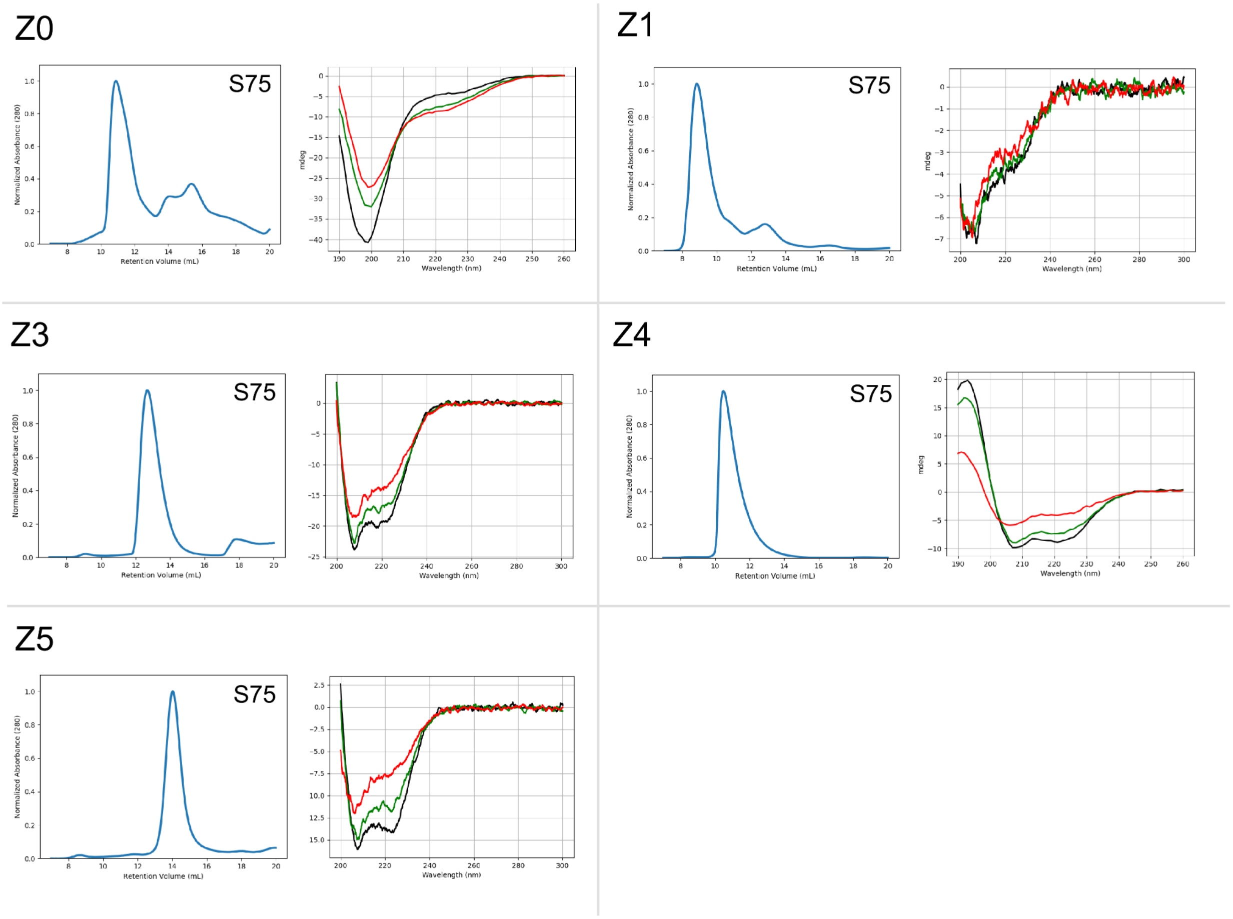


**Fig. S11. Characterization of designs that were selected for ZnO binding and soluble when expressed in *E. coli.*** (Left) SEC traces using Superdex 75 increase 10/300 GL column and (right) circular dichroism spectra at 25°C, 65°C and 95°C (black, green, and red respectively) are shown for each design.

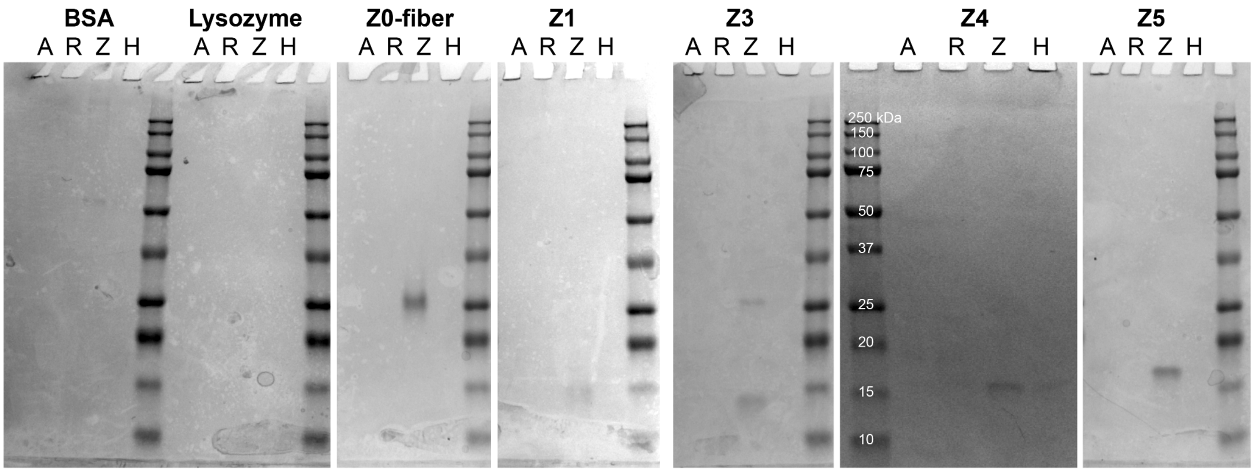


**Fig. S12. Nanoparticles based pulldown assay of selected proteins.** SDS page pull-down assay of ZnO binding designed proteins and bovine serum albumin (BSA) and lysozyme controls using target and off-target nanoparticles, A = anatase ($TiO_2$), R = rutile ($TiO_2$), Z = zinc oxide (ZnO), H = hematite ($Fe_2O_3$). All wells show protein associated with the nanoparticle pellet after three washes. The experiment was performed in solutions containing 150 mM NaCl 20 mM Tris-HCl pH 8.

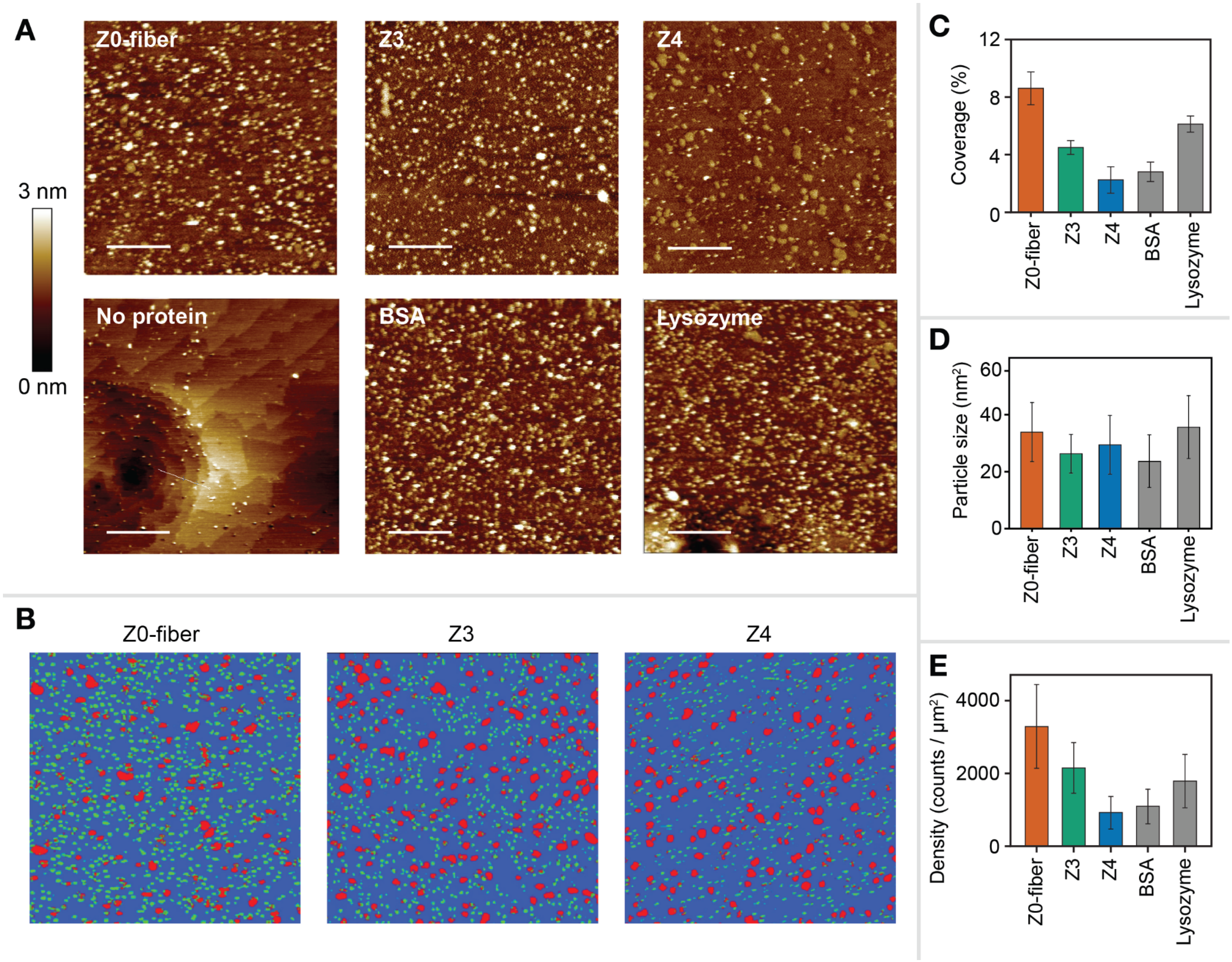


**Fig. S13. AFM and quantification of protein coverage on zinc-termination ZnO (001)** (**A**) Zinc-terminated ZnO (001) surface incubated in solutions containing 20 mM HEPES (pH 8), 50 mM NaCl, and 1 µM $Zn(NO_3)_2$, with 1 nM of Z0-fiber, Z3, or Z4, without protein, and with 1 nM BSA or lysozyme controls. Scale bars are 250 nm. (**B**) Size and morphology based identification of background (blue), proteins (green), and Zn-hydroxide (red) clusters performed using ImageJ with the Labkit plugin. Four images were analyzed for each sample, one example of which is shown for each design. (**C**) Protein coverage was determined by size and morphology from four AFM images of each case. Error bars show one standard deviation, n = 4. (**D**) Average size of blocks identified as protein (green in panel B) in four AFM images of each case. Error bars show one standard deviation, n = 4. (**E**) Number density of proteins calculated as total protein coverage / predicted average size in four AFM images of each case. Error bars show one standard deviation, n = 4.

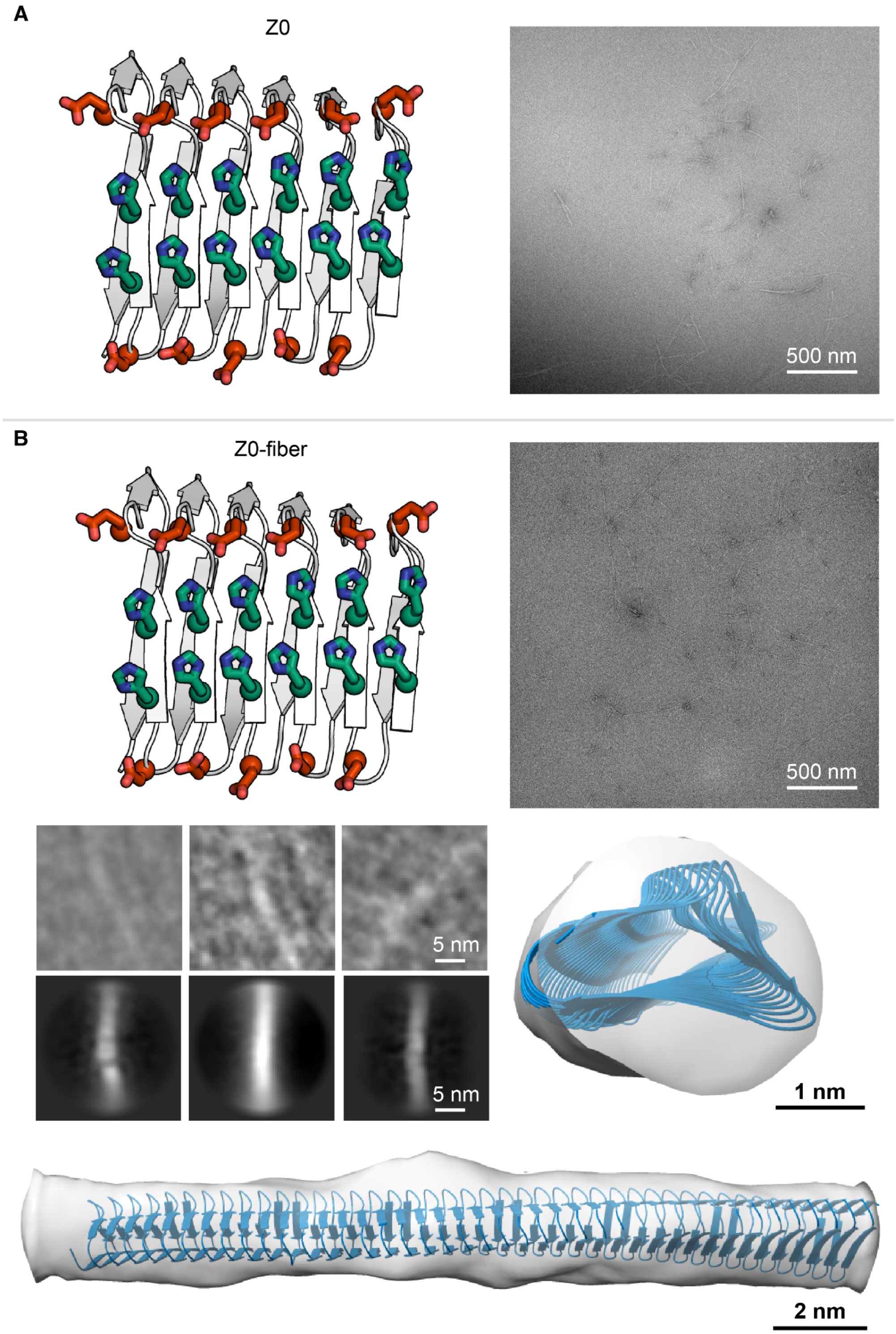


**Fig. S14. Transmission electron microscopy images of Z0-fiber.** Aspartate and glutamate sidechains shown in red, histidine sidechains shown in green. (**A**) model and negative stain TEM image of the Z0 capped DBS. (**B**) Models, negative stain and 3D averaging of the Z0-fiber uncapped DBS.

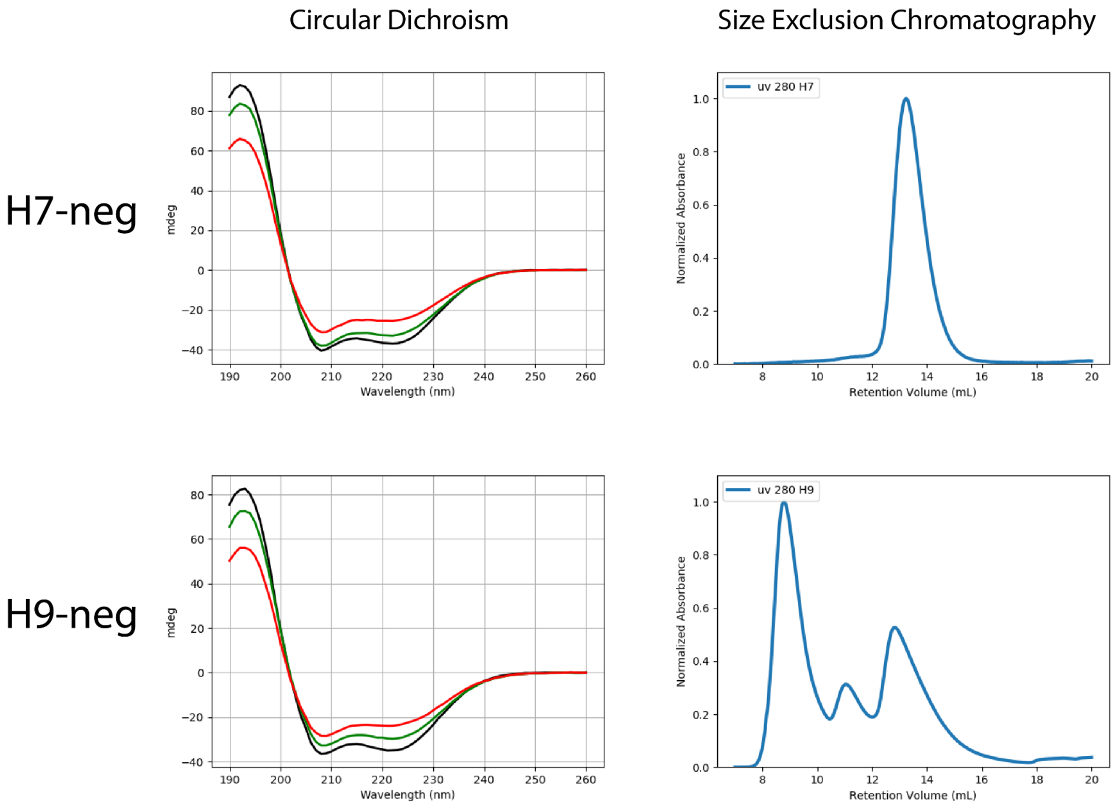


**Fig. S15. Characterization of designs that were selected for hematite binding and soluble when expressed in *E. coli,* following solubility enhancing mutations.** (Left) SEC traces using Superdex 75 increase 10/300 GL column and (right) circular dichroism spectra at 25°C, 65°C and 95°C (black, green, and red respectively) are shown for each design.

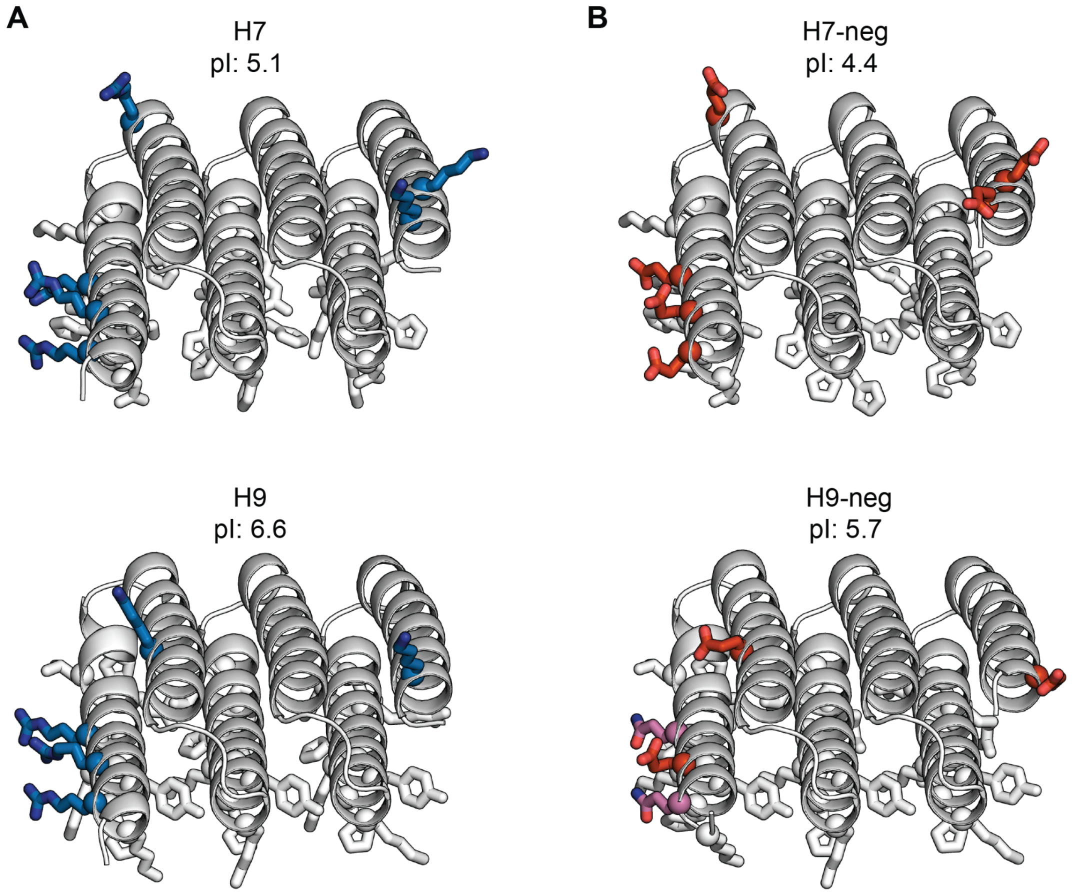


**Fig. S16. Expression enhancing mutations in H7-neg and H9-neg**. (**A**) The designs present in the library that were identified as hematite binders. (**B**) Variants of the designs with expression-enhancing mutations. Residues that vary between the original and -neg variants are shown as sticks and colored blue for arginine and lysine; red for glutamate and aspartate. Residues in the selected hematite-binding interfaces (which were unchanged) are shown as grey sticks.

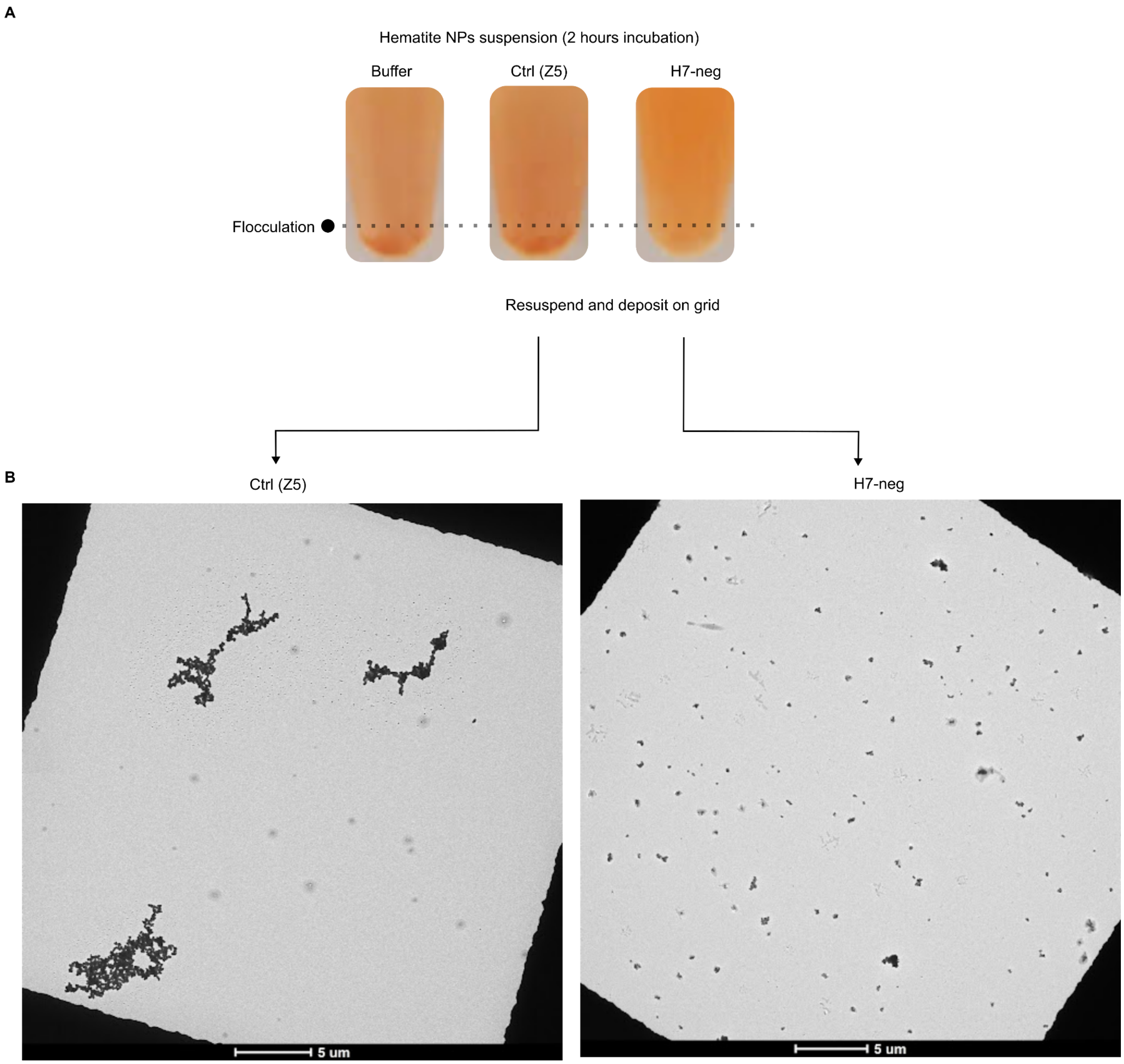


**Fig. S17. Stabilization of Hematite NP suspension by selected interfaces.** (**A**) Suspension of 0.02 wt/vol% hematite nanoparticles (≤100 nm average particle size) incubated for two hours in buffer (Tris 30 mM, NaCL 50 mM, pH8) (left), containing 0.075 mg/mL control protein (Z5) (middle), or 0.075 mg/mL identified hematite binder (H7-neg) (right). (**B**) TEM micrograph of control (Z5) and H7-neg resuspensions.

**A**

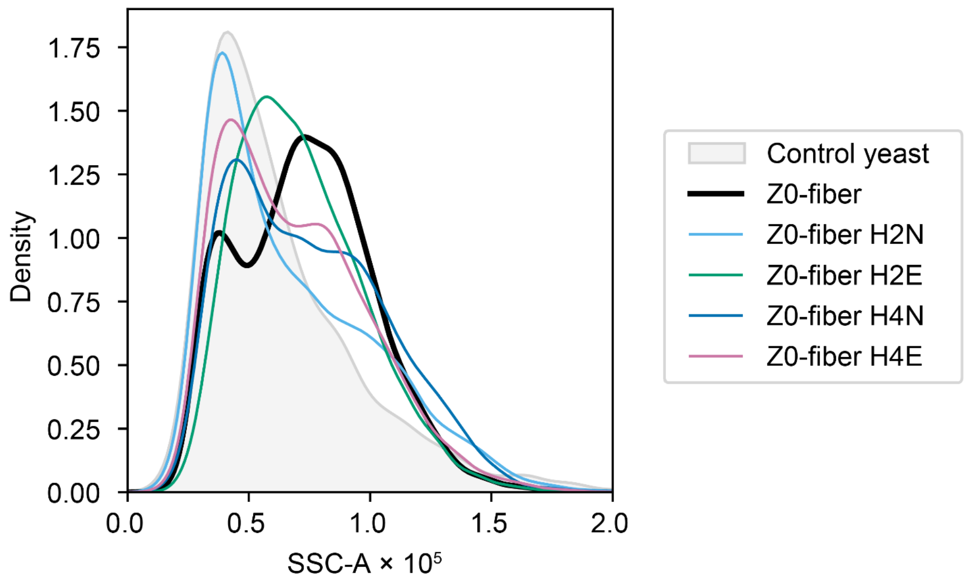


**B**

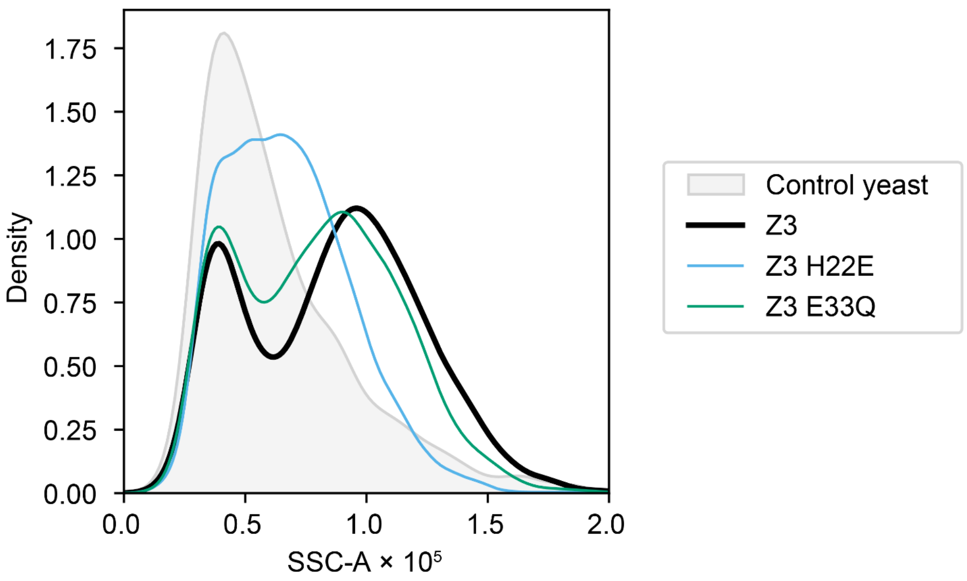


**Fig. S18. Comparison of SSC signal of ZnO-binders clones to mutated controls.** Histograms plotting the SSC signal of yeast displaying Z0-fiber (**A**) and Z3 (**B**) with respect to clones with selected mutations after incubation with ZnO NP. Mutants are named by the first residue that is mutated, but all repeated instances of these residues are mutated as well. **Fig. 2, D and E** illustrate the sets of the repeated residues.

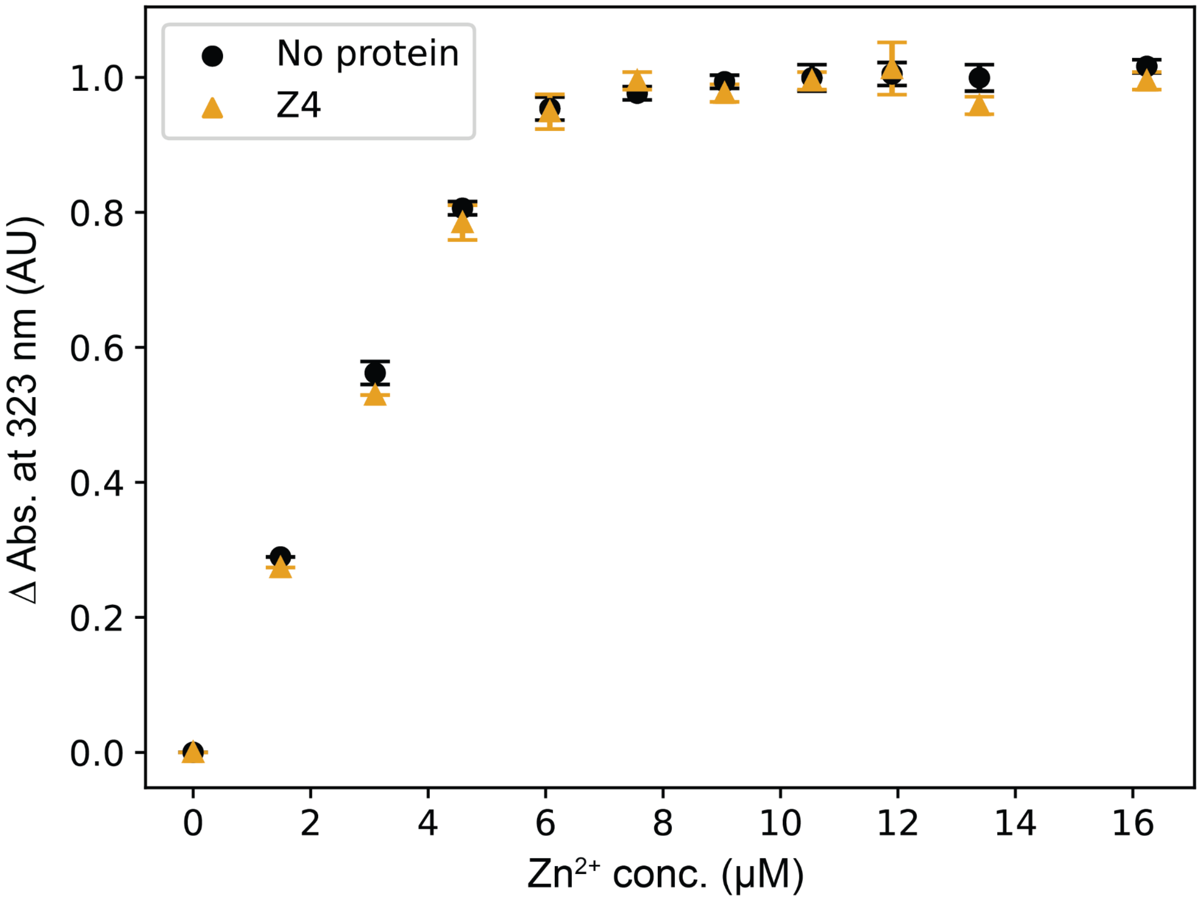


**Fig. S19. No evidence of $Zn^{+2}$ ion binding by Z4 seen in ratiometric dye (Mag-Fura-2) assay.** Kd was estimated to lie above the quantification limit of the assay at the experimental conditions (> 1 μM).

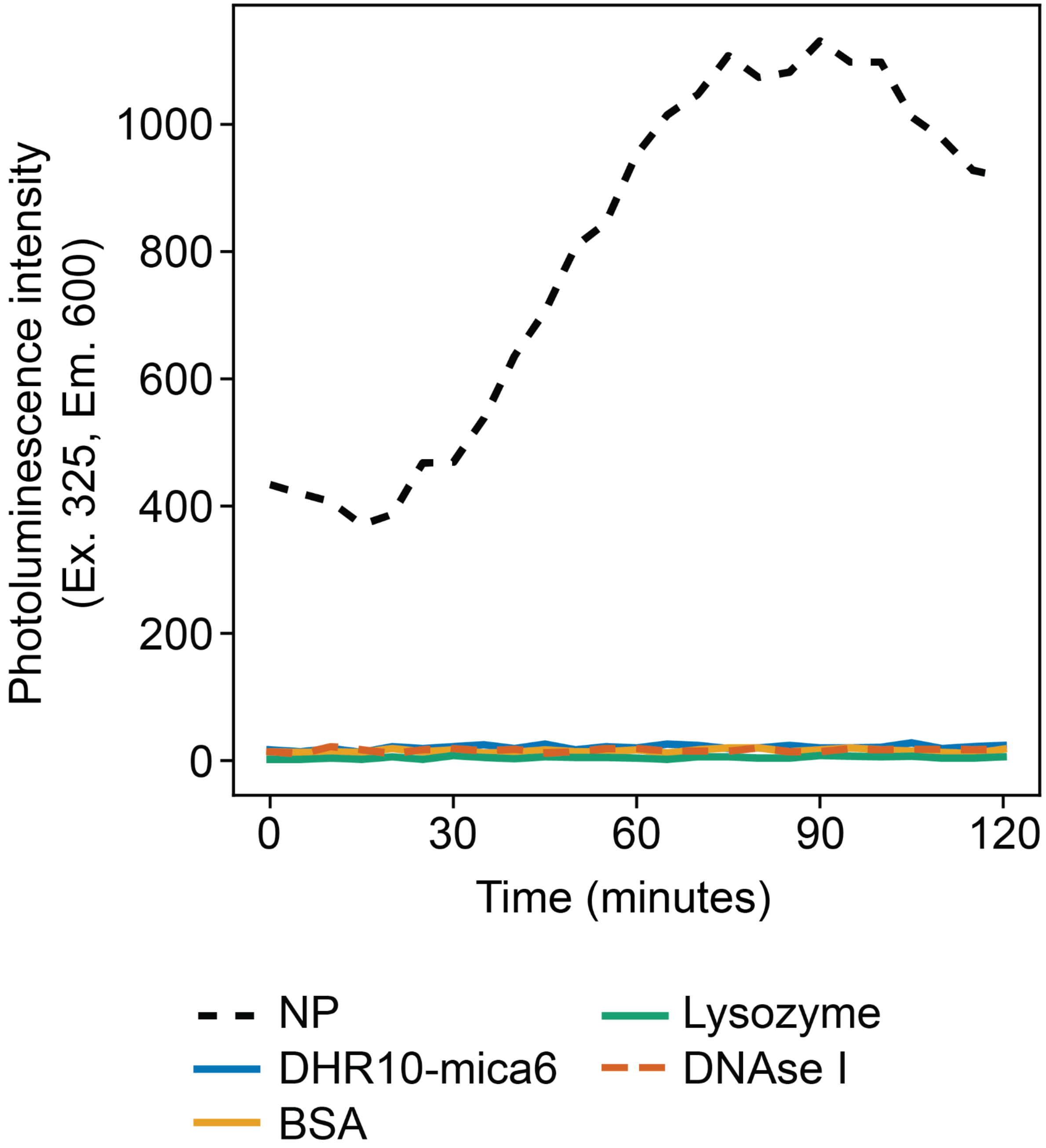


**Fig. S20. Effect of ZnO growth of diverse control proteins.** Time series of photoluminescence (Ex. 325. Em. 600) over time of solutions supersaturated for ZnO (3 mM $Zn(NO_3)_2$, 50 mM NaCl, 100 mM HEPES pH 8.2) containing negative control proteins and ZnO NP. All protein concentrations are 0.1 mg / mL and NP concentration is 12.5 µg/mL.

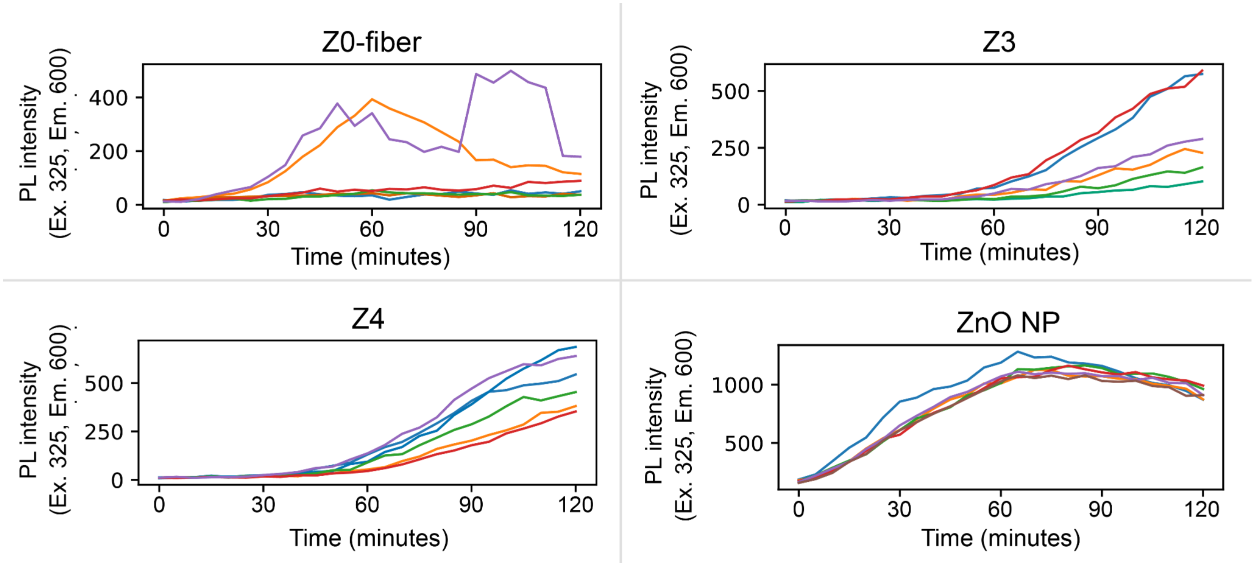


**Fig. S21. Photoluminesce (PL) emission over time for six replicates of growth assay.** Solutions contain 3 mM $Zn(NO_3)_2$, 50 mM NaCl, 100 mM HEPES pH 8.2, and contain 0.1 mg/mL protein or 12.5 µg/mL ZnO NP as indicated.

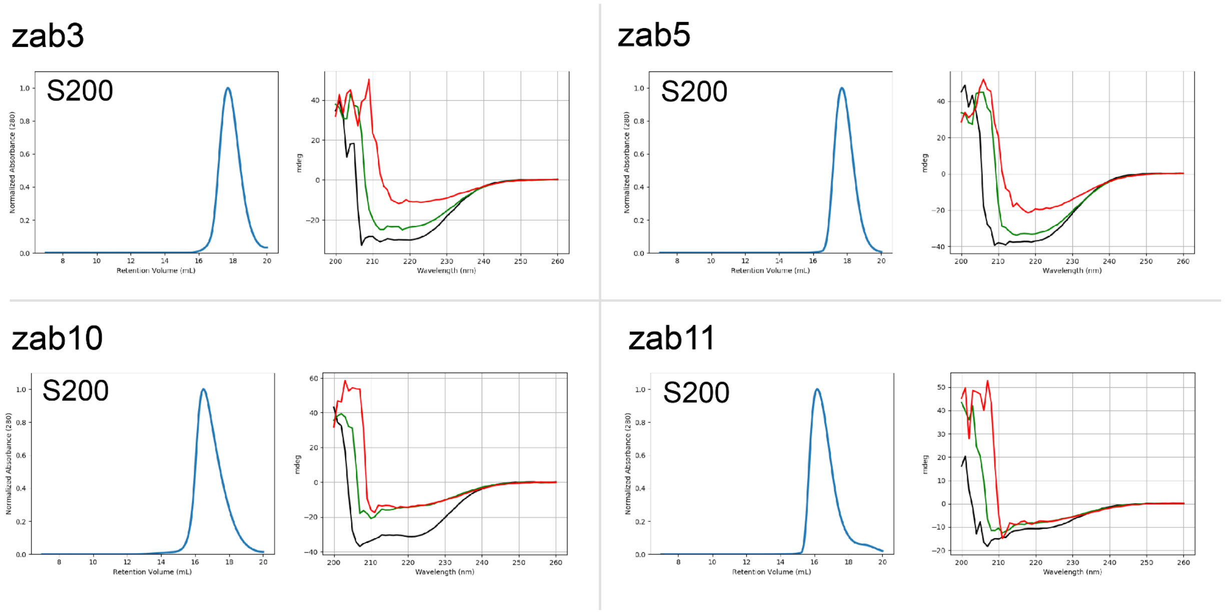


**Fig. S22. Characterization of Z0 based monomer redesigns.** SEC traces for zab3, zab5, zab10, and zab11 using Superdex 200 increase 10/300 GL column. Circular dichroism spectra for the same designs (right, black 25°C, green 65°C, red 95°C).

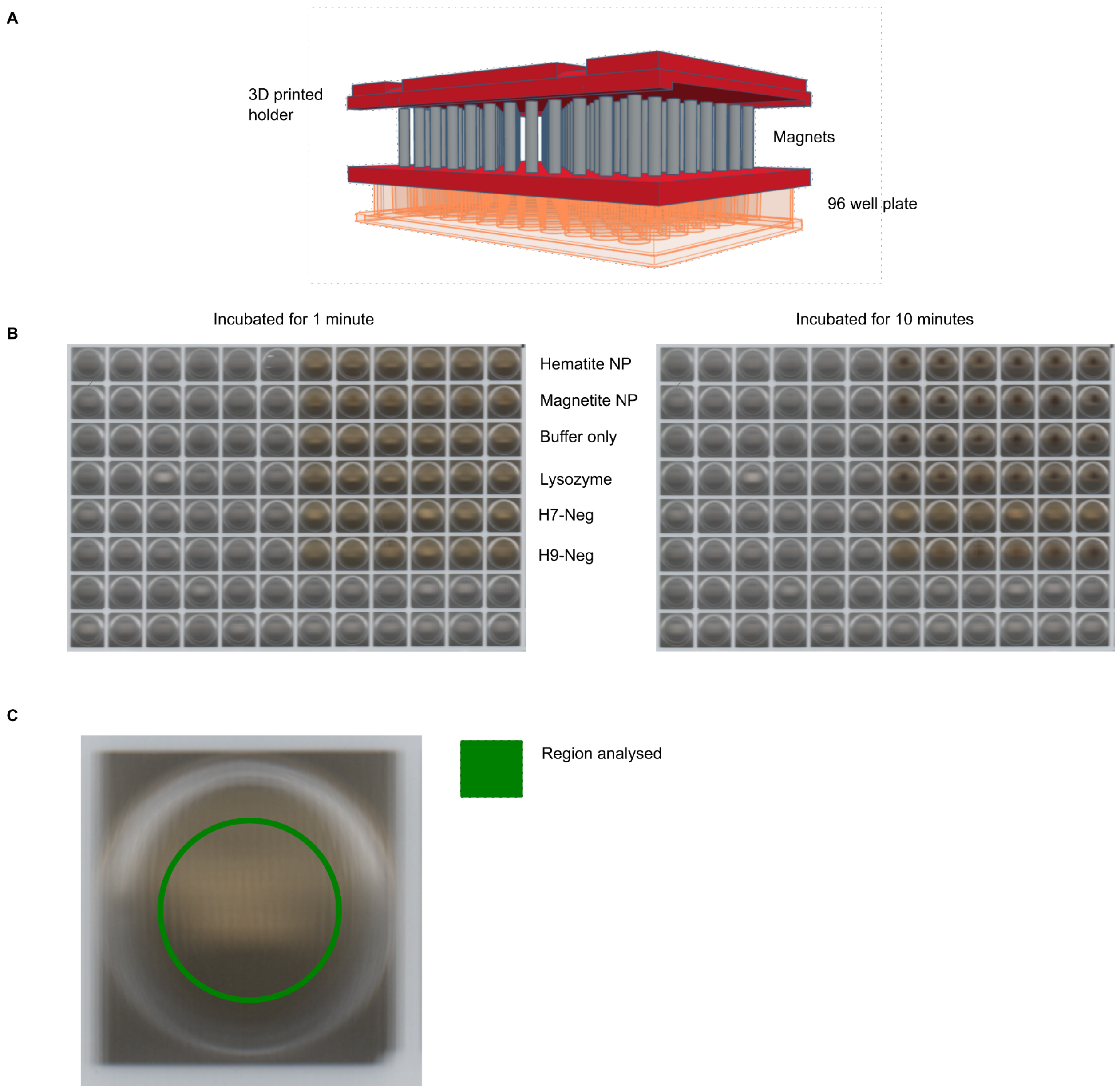


**Fig. S23. Magnetic/colorimetric assay setup.** (**A**) 3D model of the printed magnetic plate cover. (**B**) Sample plate scans showing the wells content at 1 and 10 minutes. All wells also include 100 mM CHES, 50 mM $KNO_3$, 50 mM $K^+$, 50.3 mM $NO_3^-$, 1 mM $Fe^{3+}$, 0.5 mM $Fe^{2+}$, 1 mM $NH_4^+$, and 1 mM $SO_4^{2-}$ at pH 8.7, prepared in anaerobic conditions (see Methods). (**C**) Illustration of the region analyzed to produce the time series plots.

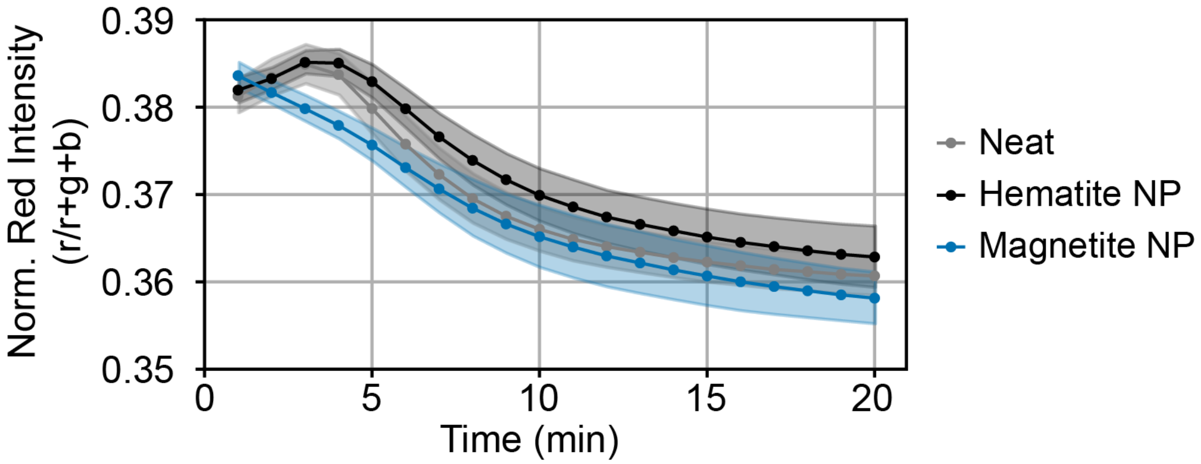


**Fig. S24. Performance of nanoparticle seeds in colorimetric assay.** In magnetite growth conditions, neither hematite or magnetite NP resulted in the formation of red material indicative of hematite.

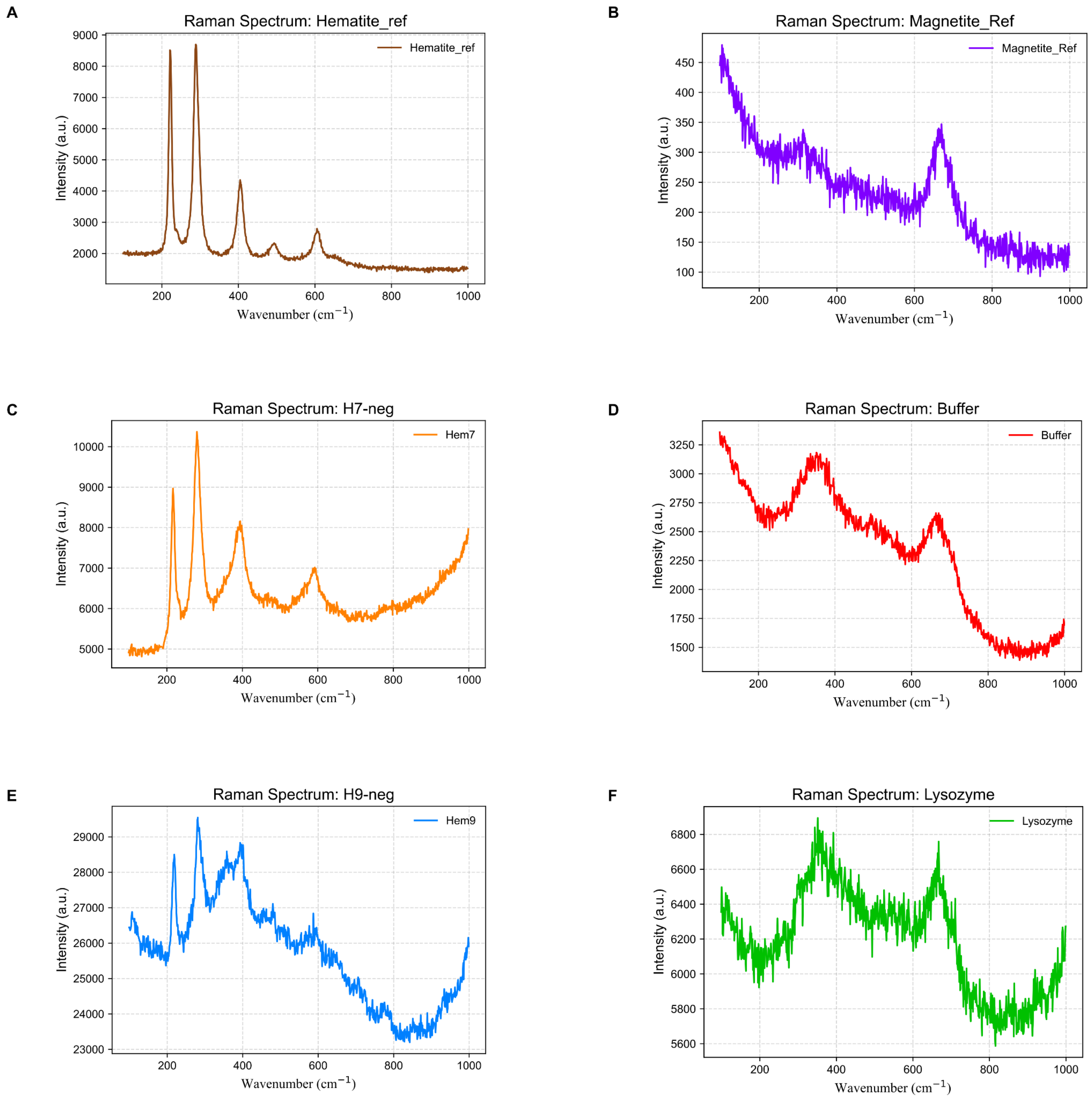


**Fig. S25. Raman spectra of individual samples.** Raw spectra of (**A**) Hematite reference NP. (**B**) Magnetite reference NP. (**C**) H7-neg sample. (**D**) Buffer only sample. (**E**) H9-neg sample. (**F**) Lysozyme sample.

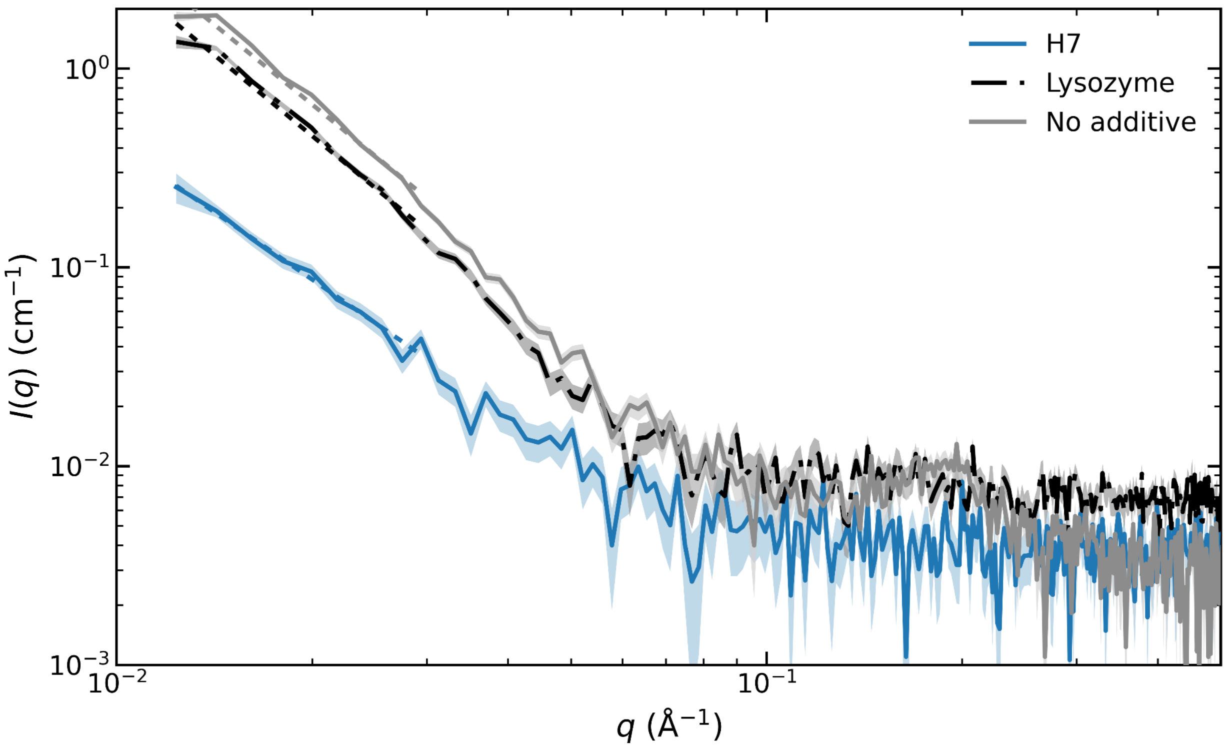


**Fig. S26. SAXS analysis of individual samples (Iron oxide).** Background-subtracted SAXS profiles of iron oxide mineralization products generated in the presence of H7-neg, lysozyme, or buffer alone. Shaded regions indicate the propagated experimental uncertainty. Dashed lines indicate power-law fits performed over the low-q range, yielding power-law exponents of −2.26 ± 0.13 (H7-neg), −2.70 ± 0.12 (lysozyme), and −2.67 ± 0.15 (no additive).

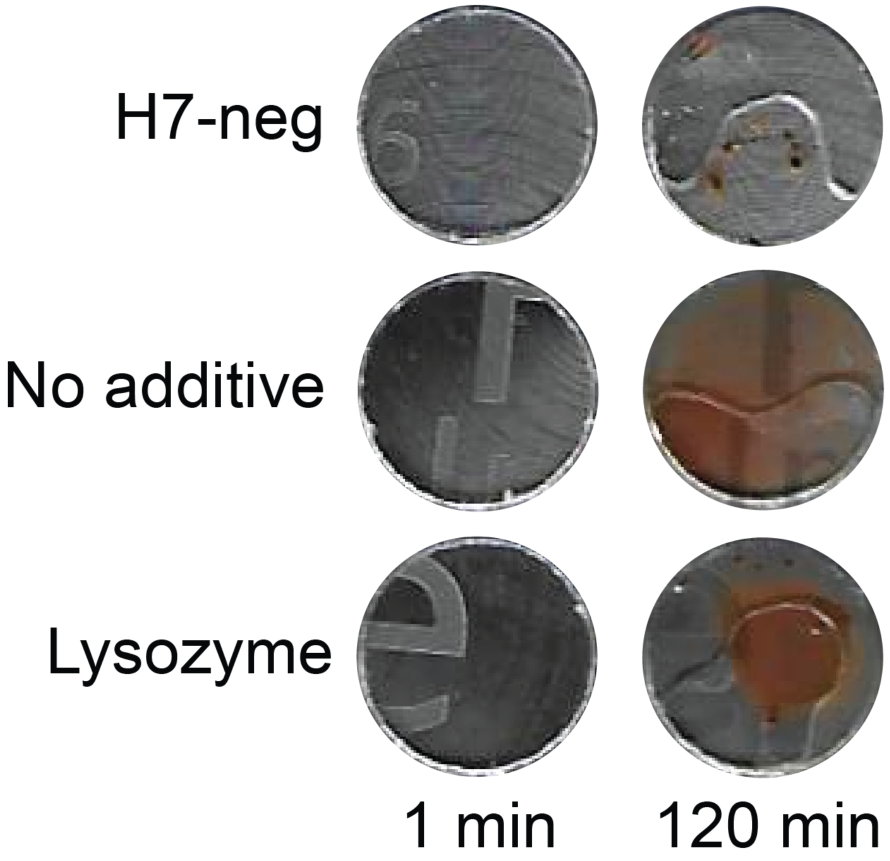


**Fig. S27. Effect on solid iron substrate.** Images from iron substrate scans prior and following two hours incubation with 10 µl samples containing 0.6 mg/ml of H7-neg or Lysozyme (0.6 mg/ml), or no additive.

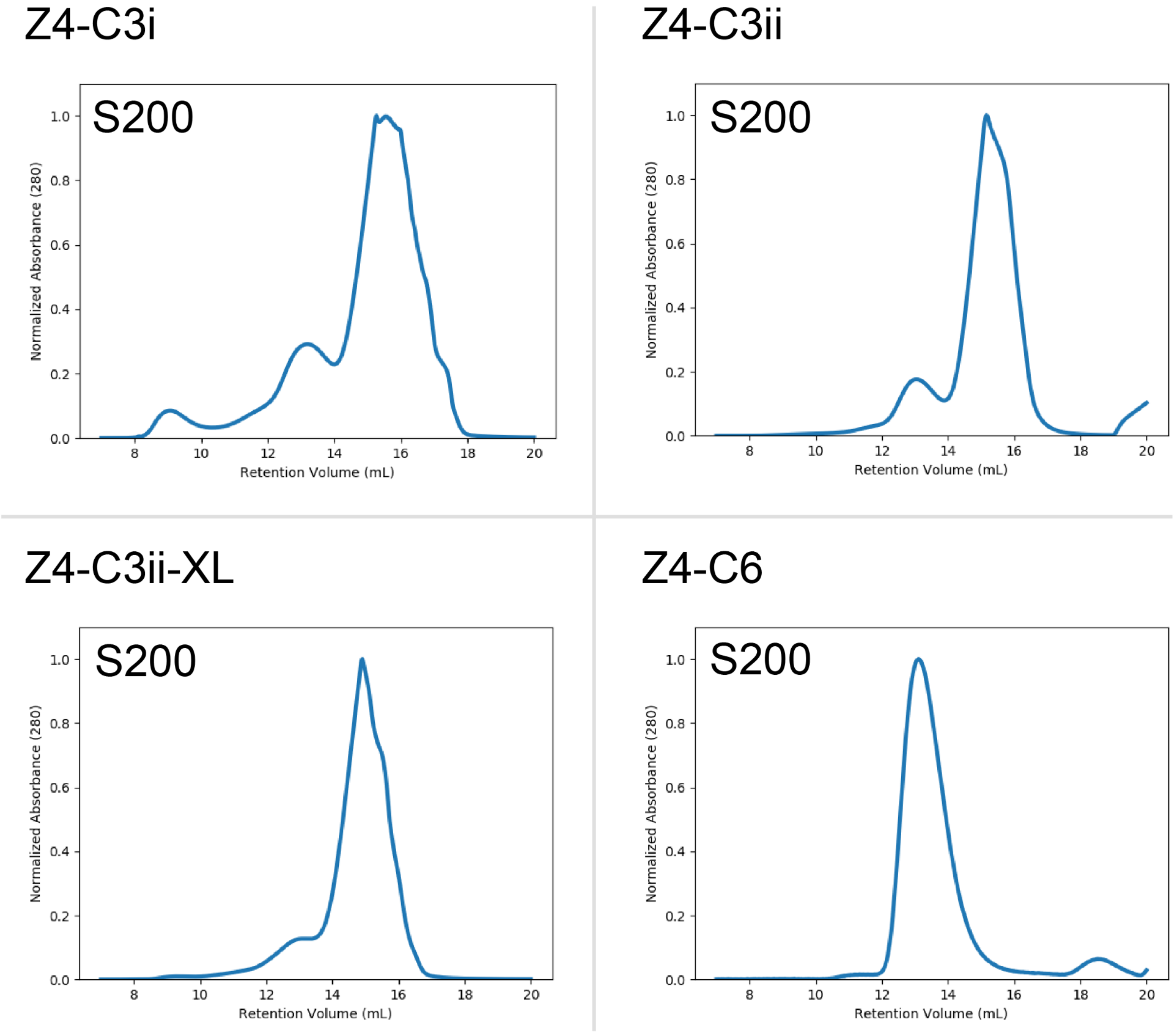


**Fig. S28. Characterization of Z4 based cyclic oligomers.** SEC traces for Z4-C3i, Z4-C3ii-XL, Z4-C3ii, and Z4-C6, using Superdex 200 increase 10/300 GL column.

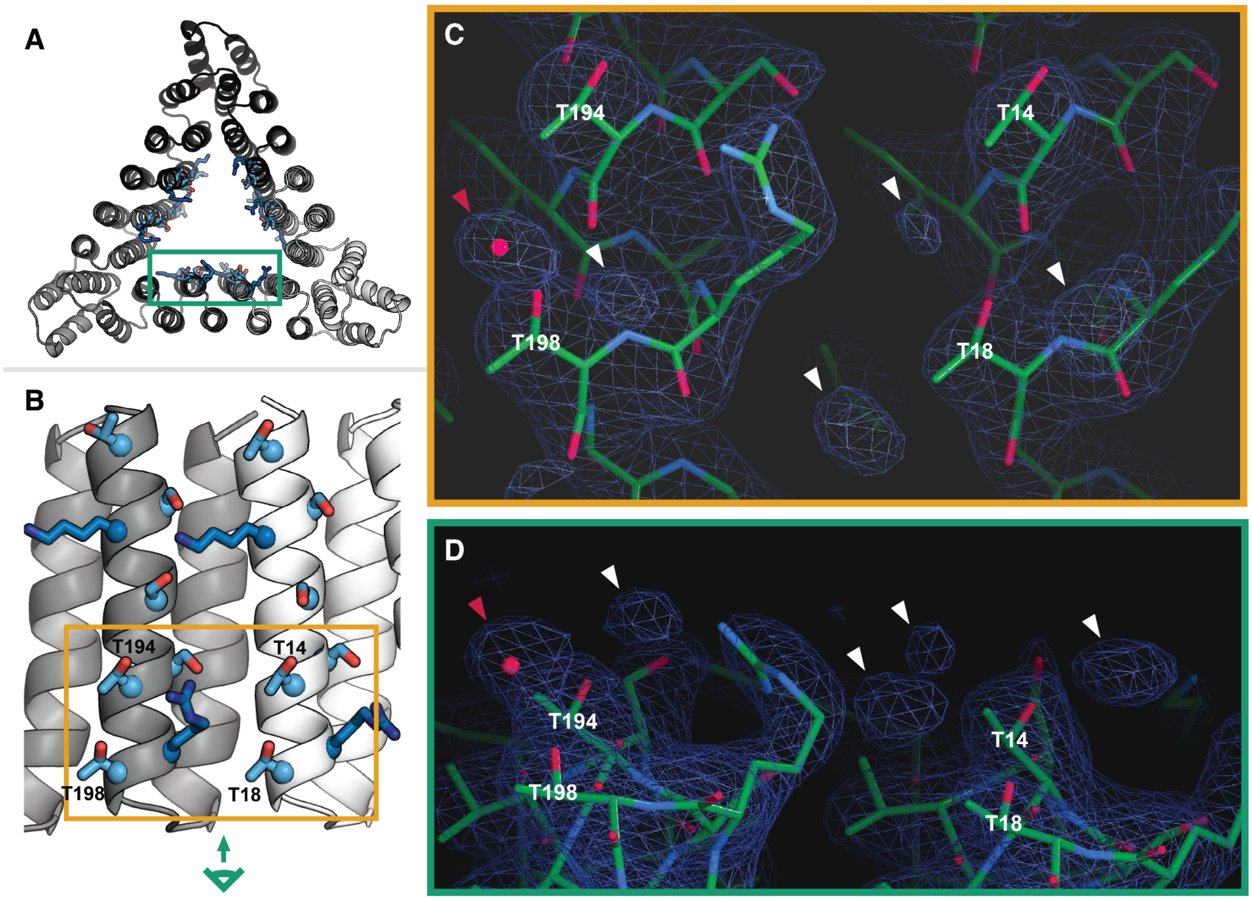


**Fig. S29. Crystallographic electron density at the Z4-C3i ZnO templating interface.** The 2mFo-DFc electron density map of the Z4-C3i crystal structure shows density near the ice-binding protein-like threonine motifs in the Z4 interface consistent with ordered waters. (**A**) Z4-C3i trimer in crystal structure (pdb_00009d92). Designed sidechains of Z4 interface lining interior cavity shown as sticks. Backbone cartoons of chains A, C, E shown in white, black, and grey respectively. (**B**) Close up view of Z4 interface highlighted in green box in panel **A**. Side chains in Z4 interface shown as sticks, threonine and serine residues colored light blue, arginines and lysines colored dark blue. Backbone shown as cartoon with chain A colored white and chain C colored grey. (**C**) and (**D**) Two views of electron density map (blue mesh) in the region within the orange rectangle in panel B. Threonine residues in Z4 interface are labeled. The red arrowhead indicates a site where water was placed during refinement, while white arrowheads indicate sites of density that are consistent with ordered waters but where waters were not placed due to insufficient resolution (3.06 Å). (**C**) Shows the same view as the orange rectangle in panel **B**, while (**D**) shows the side view indicated by the green view angle symbol in panel **B.**

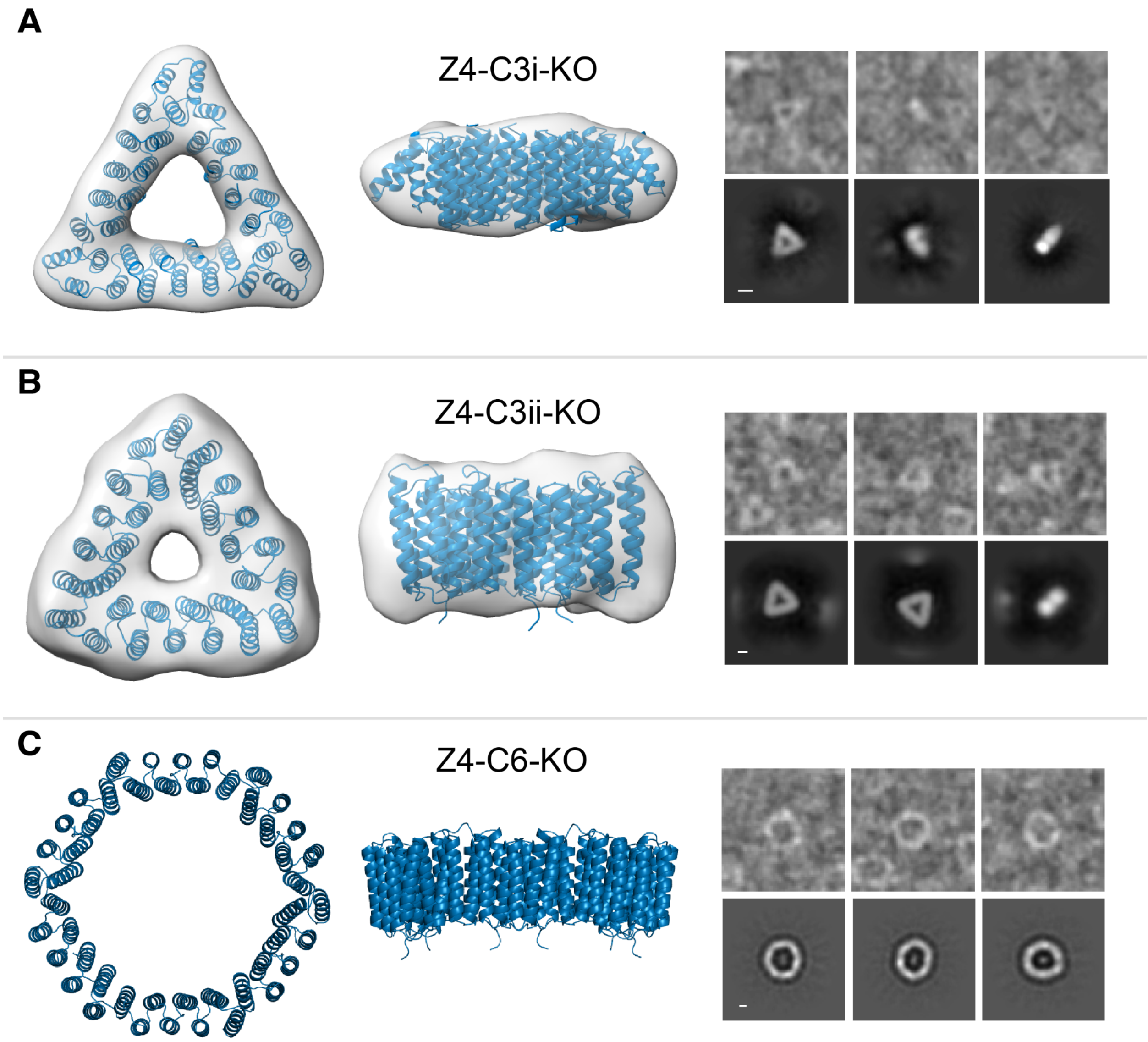


**Fig. S30. Negative stain TEM of Z4 oligomer interface knockouts.** Oligomers with the Z4 interface knocked out maintain the assembly geometry of the functional versions. Design models, 3D reconstructions, picked particles, and 2D averages of Z4-C3i KO (**A**) and Z4-C3ii KO (**B**). (**C**) Design models, picked particles, and 2D averages of Z4-C6 KO. All scale bars are 2 nm.

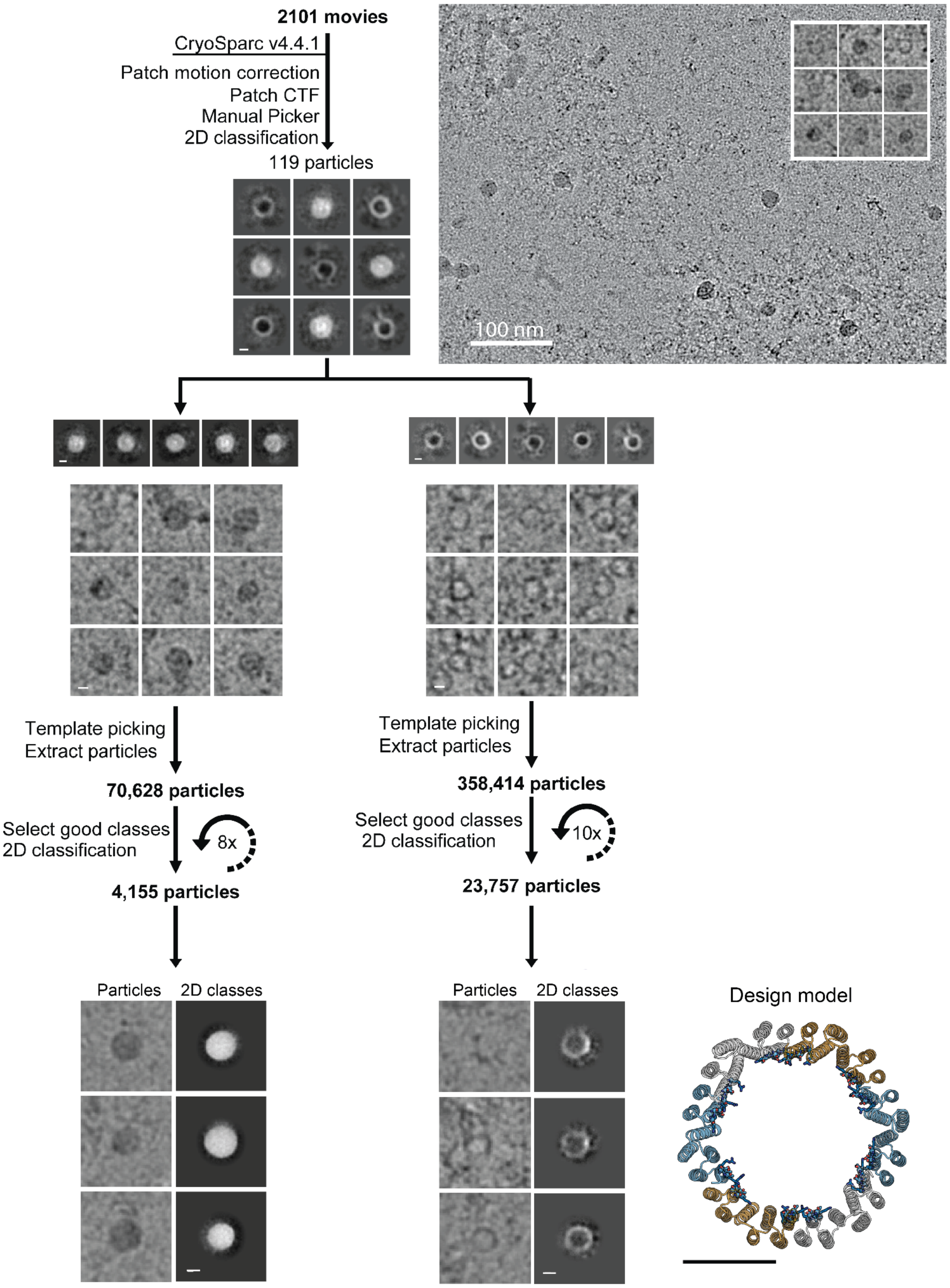


**Fig. S31. Cryo-TEM analysis pipeline of product of ZnO mineralization reaction in presence of Z4-C6 oligomer.** 2D classes resembling filled and unfilled rings were separated. All scale bars are 4 nm unless otherwise specified.

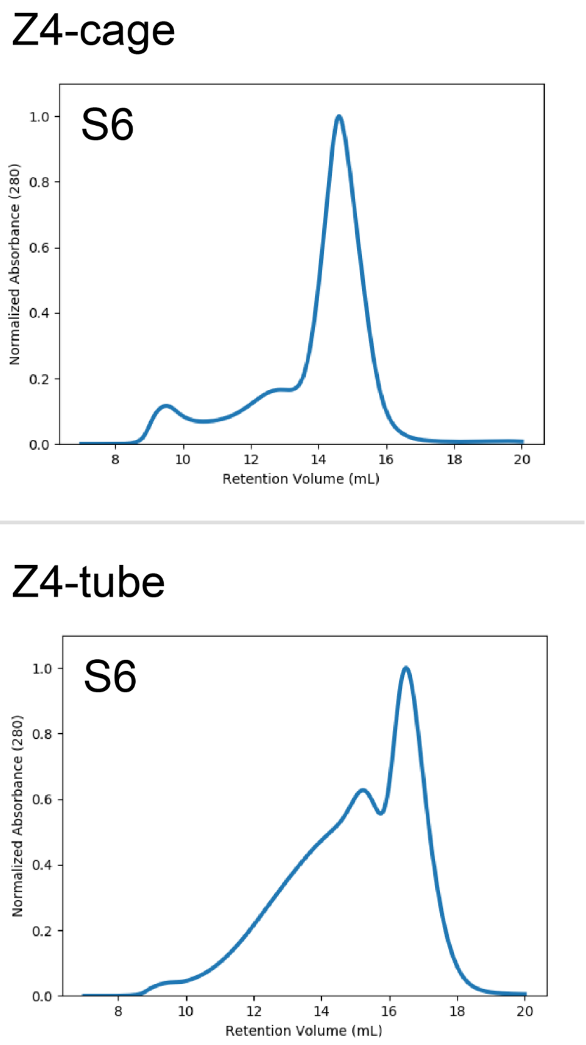


**Fig. S32. Characterization of large assemblies containing Z4 interfaces.** SEC traces for Z4-cage and Z4-tube using Superdex 6 increase 10/300 GL column.

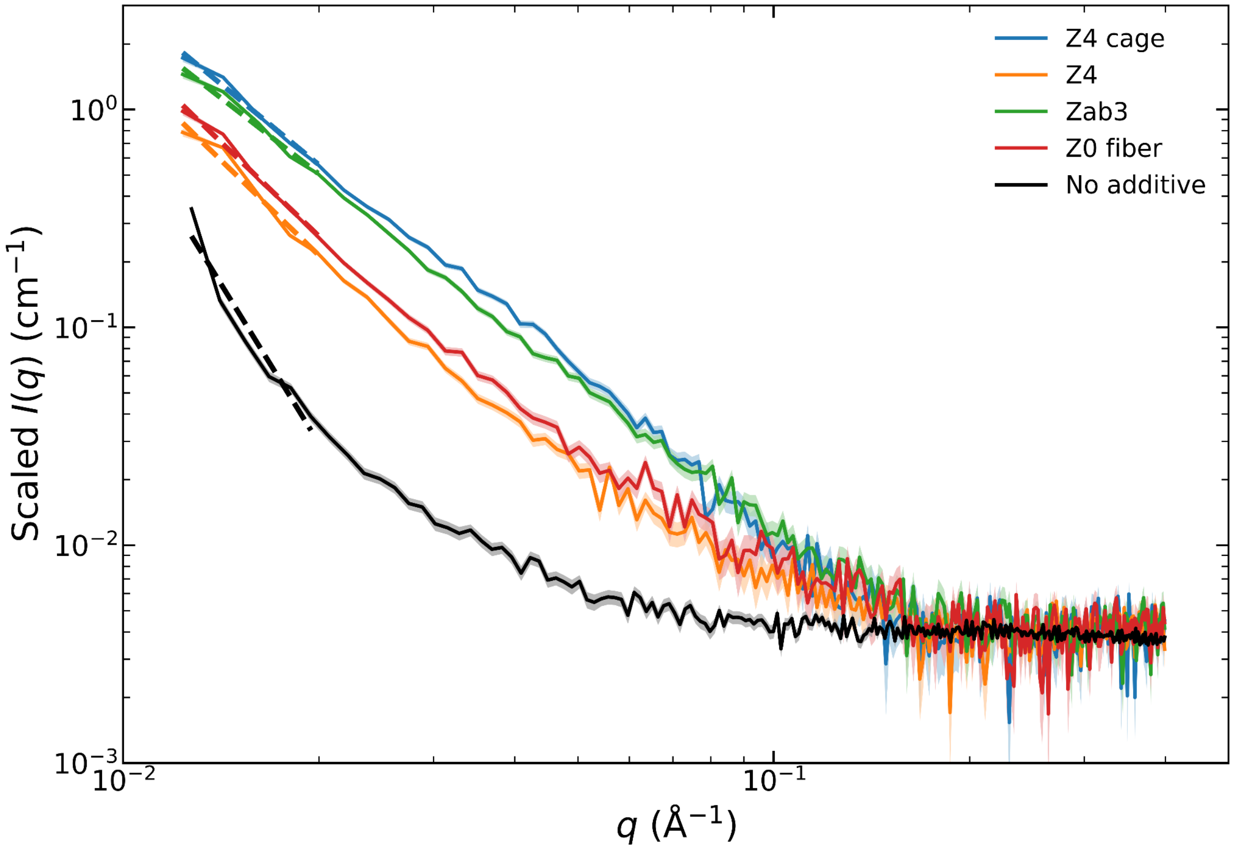


**Fig. S33. SAXS analysis of individual samples (ZnO).** Background-subtracted SAXS profiles of ZnO mineralization products generated in the presence of the designed proteins Z4-cage, Z4, zab3, and Z0-fiber, or no protein additive. The no additive sample was prepared at twofold higher precursor concentration to permit homogeneous nucleation (see Methods). Curves were vertically scaled to a common intensity level over the high-q range while preserving the measured q-values and curve shapes. Dashed lines indicate power-law fits to the low-q region. Shaded regions represent the propagated experimental uncertainty. The corresponding power-law exponents were −2.46 ± 0.17 (Z4-cage), −2.90 ± 0.31 (Z4), −2.32 ± 0.19 (zab3), −2.87 ± 0.19 (Z0-fiber), and −4.86 ± 0.65 (no additive).

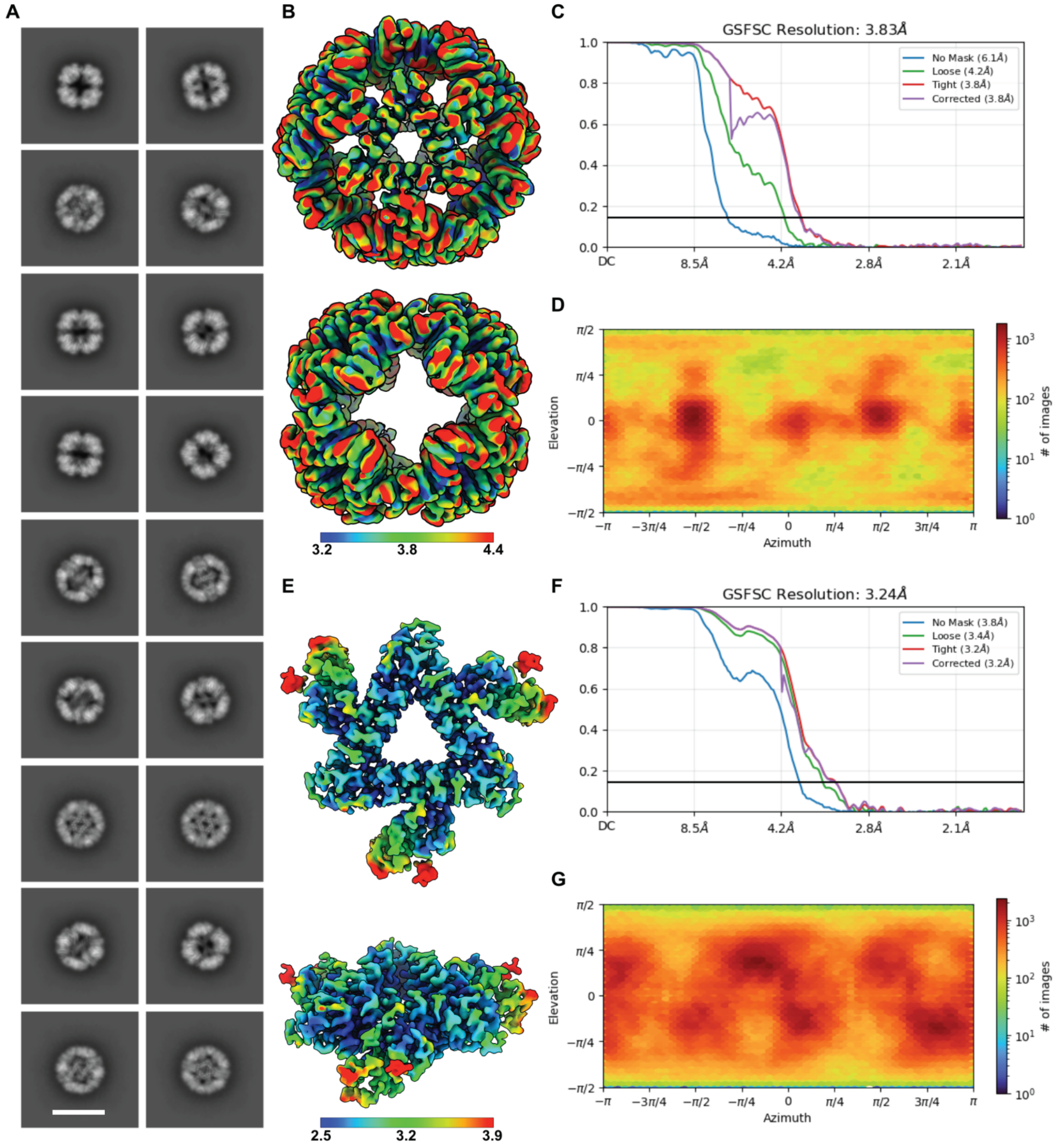


**Fig. S34. Cryo-TEM Data Processing Statistics** (**A**) Enlarged 2D class averages of the Z4-cage showing clearly designed secondary structure features consistent with the design model. Scale bar is 18 nm. (**B**) Local resolution estimation of the Z4-cage refined with relaxed octahedral symmetry. Local resolution estimation ranges from ~3.2 Å to ~4.4 Å. (**C**) Global FSC plot with an estimated global resolution of 3.83 Å. (**D**) Orientational distribution plot showing full angular sampling of the Z4-cage reconstruction. (**E**) Local resolution estimation of the locally refined Z4-C3i trimer present within the Z4-cage. Local resolution estimation ranges from ~2.5 Å along the designed ice-binding motif to ~3.9 Å along the interface between Z4-C3i trimers. (**F**) Global FSC plot with an estimated global resolution of 3.34 Å. (**G**) Orientational distribution plot showing full angular sampling of the Z4-C3i trimer local reconstruction.

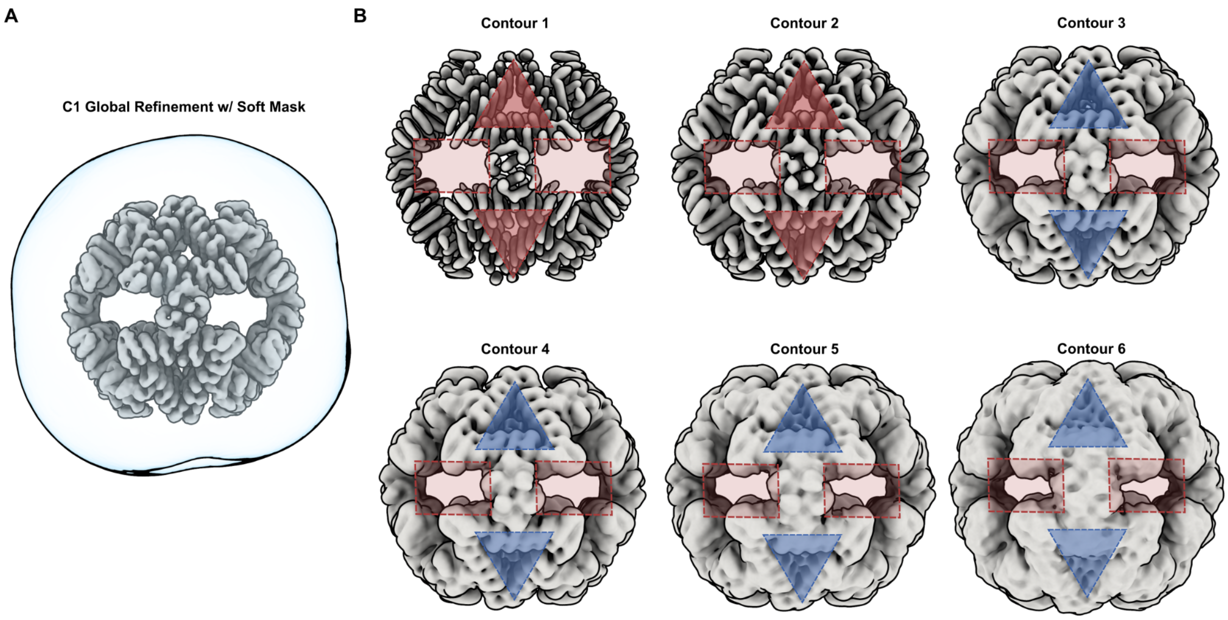


**Fig. S35. Visualization of ZnO nucleation and growth restricted to the C3 axis of the Z4-cage containing the designed ZnO interfaces. (A)** Cryo-EM 3D reconstruction of the Z4-cage (gray), generated through C1 refinement with an extremely soft mask filling all openings (blue). **(B)** The reconstruction in (A) was low-pass filtered to 8 Å and analyzed across varying contour levels. At contour level 3, non-protein density consistent with ZnO nucleation becomes apparent, specifically at the center of the C3 axis where the de novo ice-binding motif interface resides (blue triangles). No additional density is observed along the C4 axis at any contour level (red rectangles), supporting the design intent of localized ZnO nucleation and growth exclusively within the C3 axis.

**Table S1. Fold enrichments, interfacial features, and *E. coli* expression data for designs selected for ZnO binding.** Names of ordered design sequences in the library are bolded, the names of mutants of those sequences are unbolded. Fold enrichment is defined as design abundance in NGS of yeast library subpopulations after sorting for ZnO binding divided by their initial abundance in the expression library subpopulations. *E. coli* expression levels were determined visually from SDS PAGE gels on IMAC purified proteins and resuspended insoluble fractions. Topologies are identified as (AB) mixed alpha-beta designs, (DBS) designed beta-solenoids, and (DHR) designed helical repeats.

| Name | Base design | ZnO enrichment | Chelating residues | Periodic charges | Hydro-phobics | IBP motif | Topology | Soluble expression | Insoluble expression |
|---|---|---|---|---|---|---|---|---|---|
| **Z0** | n/a | 167.6 | + | + | - | - | DBS | ++ | + |
| Z0-fiber | Z0 | n/a | + | + | - | - | DBS | ++ | + |
| **Z1** | n/a | 161.7 | + | + | - | - | AB | ++ | + |
| Z2 | Z3 | n/a | + | + | - | - | DHR | - | - |
| **Z3** | n/a | 7.6 | + | + | - | - | DHR | ++ | ++ |
| **Z4** | n/a | 95.5 | - | + | - | + | DHR | ++ | + |
| **Z5** | n/a | 57.1 | + | + | - | - | AB | + | + |
| **Z6** | n/a | 310.1 | + | - | + | - | DBS | - | - |
| **Z7** | n/a | 26.7 | - | + | + | - | DBS | - | - |
| **Z8** | n/a | 197.2 | - | - | - | + | DBS | - | ++ |
| **Z9** | n/a | 553.5 | - | - | - | + | DBS | - | - |
| Z10 | Z9 | n/a | - | - | - | + | DBS | - | - |
| **Z11** | n/a | 19.0 | - | + | + | - | DBS | - | + |
| **Z12** | n/a | 6.5 | - | - | + | + | DBS | - | - |
| **Z13** | n/a | 950.5 | - | + | + | - | DBS | - | + |
| **Z14** | n/a | 27.8 | - | - | + | + | DBS | + | + |

**Table S2. Fold enrichments, interfacial features, and *E. coli* expression data for designs selected for hematite binding.** Names of ordered design sequences in the library are bolded, the names of mutants of those sequences are unbolded. Fold enrichment is defined as design abundance in NGS of yeast library subpopulations after sorting for hematite binding divided by their initial abundance. *E. coli* expression levels were determined visually from SDS PAGE gels on IMAC purified proteins and resuspended insoluble fractions. Topologies are identified as (AB) mixed alpha-beta designs, (DBS) designed beta-solenoids, and (DHR) designed helical repeats.

| Name | Base design | Hematite enrichment | Chelating residues | Periodic charges | Hydro-phobics | IBP motif | Topology | Soluble expression | Insoluble expression |
|---|---|---|---|---|---|---|---|---|---|
| **H1** | n/a | 10.4 | + | - | + | - | DBS | - | + |
| H2 | H1 | n/a | + | - | + | - | DBS | - | - |
| H3 | H1 | n/a | + | - | + | - | DBS | - | + |
| **H4** | n/a | 442.8 | + | - | + | - | DBS | - | ++ |
| H5 | H6 | n/a | + | - | + | - | DBS | - | - |
| **H6** | n/a | 18.0 | + | - | + | - | DBS | - | + |
| **H7** | n/a | 32.8 | + | - | - | - | DHR | + | + |
| H7-neg | H7 | n/a | + | - | - | - | DHR | ++ | - |
| **H8** | n/a | 70.3 | + | + | - | - | DHR | + | + |
| H9 | H8 | n/a | + | + | - | - | DHR | + | + |
| H9-neg | H9 | n/a | + | + | - | - | DHR | ++ | - |
| H10 | H8 | n/a | - | + | - | - | DHR | + | + |
| **H11** | n/a | 11581.6 | + | - | - | - | DHR | - | - |

**Table S3. Backbones and interface composition constraints of selected designs.** Chimeras and variants excluded.

| Design name | Backbone | Amino Acid Composition |
|---|---|---|
| Z0 | QAEGGQLQVQAQGNSQIEVGSNG_seq58_0004 | H40E20D20 |
| Z1 | abr_3 | Neg60 |
| Z3 | DHR14_5CWH_XtalFit | His20Cys20 |
| Z4 | DHR49_5CWJ_XtalFit | S35T35 |
| Z5 | PDL_0_4 | Pos60 |
| Z6 | QAEGGQLQVQAQGNSQIEVGSNG_seq58_0004 | posGreDiv |
| Z7 | QAQLQIQSSGSS_seq95_0002 | DEAroMidGr |
| Z8 | QIQQGT_seq132_0006 | Thr90 |
| Z9 | QIQVQAQGSNT_seq4_0003 | Thr90 |
| Z11 | QVQIQVQAQAQG_seq49_0004 | NegGre45 |
| Z12 | QVQIQVQAQAQG_seq49_0004 | Thr90 |
| Z13 | QVQVQIQSSGAS_seq131_0003 | DEAroMidGr |
| Z14 | RiAFP_4DT5 | Val10_Al50 |
| H1 | QAQLQIQASGT_seq131_0002 | His20Cys20 |
| H6 | QIQVQIQSSGGS_seq106_0005 | His20Cys20 |
| H7 | THR_8_NSR_XtalFit | H40E20D20 |
| H8 | THR_8_NSR_XtalFit | PosMidGrDv |

**Table S4. Backbones and interface composition constraints of chimeras selected during sorting and design variants.**

| Design name | Base design | Variant type |
|---|---|---|
| Z0-fiber | Z0 | Fiber |
| Z2 | Z3 | Fiber |
| Z10 | Z9 | Fiber |
| H2 | H1 | Fiber |
| H3 | H1 | Truncation |
| H4 | n/a | Chimera |
| H5 | H6 | Substitution |
| H7-neg | H7 | Solubility enhancing redesign |
| H9 | H8 | Truncation |
| H9-neg | H9 | Solubility enhancing redesign |
| H10 | n/a | Chimera |
| H11 | n/a | Chimera |

**Table S5. Amino acid sequences of enriched ZnO binders and design mutants as displayed on yeast.** Full sequences include linkers and tags.

| Name | Sequence |
|---|---|
| Z0 | MYDVPDYALQASGGGGSGGGGSGGGGSASHMDSNGSESKITRKGDRHIHV EEDATARGGTLQVEAEGDSHIHVGEDAKAEGGTLEVEAHGDSHIHVGEHA KANGGRLQVKAQGDSHIHVGENAQAKGGQLIVQAQGDSHIHVGEDAKSD GGEIRVESEGDSHIHEGENVGGGSEQKLISEEDL |
| Z1 | MYDVPDYALQASGGGGSGGGGSGGGGSASHMGEVEFDHEISEEECSHELG RANIKDVTFQADISVESIVKILGLANIKSVHFEHEISVEDCAKILGLANIKEVH FEEDISEEDERKTKGLANIGSWGGGSEQKLISEEDL |
| Z3 | MYDVPDYALQASGGGGSGGGGSGGGGSASHMSEEVNERVKQLAEKAKEA TDKHEVRKIVQELAELAQCSTDSELVNEIVKQLAEVAKEATDKHLVRKIVQI LAELAQCSTDSELVNEIVKQLEEVAKEATDKHLVRKIEQILEELKQCSTDGS WGGGSEQKLISEEDL |
| Z4 | MYDVPDYALQASGGGGSGGGGSGGGGSASHMDSTEESKRISRISTEARTSG TEESLRQAIEDVAQLAKKSQDSTVLSKAISVISTIARTSGSEEALRQAIRAVAE IAKEAQDPTVLSKAISVISTIARTSGSEEARRQAERAEEEIRRRAQGSWGGGS EQKLISEEDL |
| Z5 | MYDVPDYALQASGGGGSGGGGSGGGGSASHMMRTPLLKLIRIVRTGSEEQ QKSALEQLKRLLEAGADPNAADNHGRTPLLKLIRIVRTGSEEQQKAALELL KLLLEAGADPNAADNHGRTPQQKLKRIERTGSEEQQKAAKELLKLLEEAGA DPNAADNHGSWGGGSEQKLISEEDL |
| Z6 | MYDVPDYALQASGGGGSGGGGSGGGGSASHMDSNGSESKITRKGRRVIHV EHDATARGGTLQVEAEGLSAIFVGLDAKAEGGTLEVEAHGPSTIQVGAHAK ANGGRLQVKAQGHSRIHVGQNAQAKGGQLIVQAQGKSYIYVGRDAKSDG GEIRVESEGKSHIYEGSNVGSWGGGSEQKLISEEDL |
| Z7 | MYDVPDYALQASGGGGSGGGGSGGGGSASHMDRHSEINSSSDAYAEEMID SEDNARATLVINSDGDSEAELVIFSDGDSQAKLEINSKGDSIADLEIESLGDSS ATLRIHSKGESEAYLVIISDGDSEAQLQIVSDGDSDAVLEIDSEGSSSAQLQIIS KGESDAFSDIESASSKSVQIEQDASGSKGSWGGGSEQKLISEEDL |
| Z8 | MYDVPDYALQASGGGGSGGGGSGGGGSASHMKRDDHVTETTGSKVDDG VEINAWTTITTGTEINAGTVINERTTITTGTIINEGTEINEGTTITTGTEINAGTK INAGTTITTGTRINAGTEINEGTTITTHTTINRGTHINAGTTITTHTEINEGTTIN SGTTITTHTKIDSGSKIDSGSTITTGRGSWGGGSEQKLISEEDL |
| Z9 | MYDVPDYALQASGGGGSGGGGSGGGGSASHMTIQKERQGPTSTITVTSTSD DTQIEVKAEGPKTTITVTATGNKTRINVVAHGPTTTITVTATGNQTSINVEAE GDTTTITVTATGDDTSIRVVARGDTTTITVTATGDNTSIEVEAEGNNTTITVTA |

| | |
|---|---|
| | TGNNTQISITSRGDNTTITKTSTGSNRGSWGGGSEQKLISEEDL |
| Z10 | MYDVPDYALQASGGGGSGGGGSGGGGSASHMNTTIQVEAQGPTTTITVTAT GDDTQIEVKAEGPKTTITVTATGNKTRINVVAHGPTTTITVTATGNQTSINVE AEGDTTTITVTATGDDTSIRVVARGDTTTITVTATGDNTSIEVEAEGNNTTITV TATGNNTQISITARGDNTTITVTATGGSWGGGSEQKLISEEDL |
| Z11 | MYDVPDYALQASGGGGSGGGGSGGGGSASHMDVERDVKRESSTEVIIVVV SESETRVSITVQASADGEVIIVVVAEAEGEVSITVQAEAKGEVIIVVVAEAEG QVHISVQARADGEVIIVVVAEAEGEVHIQVEAEAKGEVIIVVVAEAEGQVVI SVSAKAEGEVIIVIVSEAEGESSVSESKSSNGGSWGGGSEQKLISEEDL |
| Z12 | MYDVPDYALQASGGGGSGGGGSGGGGSASHMDVERDVKRESSTTVYITVT STSTTRVSITVQASADGTVYITVTATATGEVSITVQAEAKGTVYITVTATATGQ VHISVQARADGTVYITVTATATGEVHIQVEAEAKGTVYITVTATATGQVVISV SAKAEGTVYITITSTATGESSVSESKSSNGGSWGGGSEQKLISEEDL |
| Z13 | MYDVPDYALQASGGGGSGGGGSGGGGSASHMDKTVVINSTKDDEIVVVIK SDKDEKVQVEIQSEGADEVVVEIESTGAKQVTVQIRSSGADEVVVQIRSHG AEQVIVEIESTGADEVVVQIQSTGAQQVQVDIKSTGADEVVVQIQSTGAKE VEVSIKSEGADESVVEIDSKGAERITVSRDDKGSEGSWGGGSEQKLISEEDL |
| Z14 | MYDVPDYALQASGGGGSGGGGSGGGGSASHMGYSCRAVGVDGRAVQDTQ GTCTAQAAGAGAMASGTSEPGSASTAQAAGRGATARSTSTGRGAATTQAA GTASATSNAIGQGAATTQAAGSAGGRATGSATTSSSASQPAQTQQIAGPGFQ TAKSFARNAATTQVAASHGSWGGGSEQKLISEEDL |
| Z0-fiber | MYDVPDYALQASGGGGSGGGGSGGGGSASHMSHIHVGENADANGGELKV TAKGDSHIHVGEDATARGGTLQVEAEGDSHIHVGEDAKAEGGTLEVEAHG DSHIHVGEHAKANGGRLQVKAQGDSHIHVGENAQAKGGQLIVQAQGDSHI HVGEDAKADGGELRVEAEWGGGSEQKLISEEDL |
| Z0-fiber H2N | MYDVPDYALQASGGGGSGGGGSGGGGSASHMSNIHVGENADANGGELKV TAKGDSNIHVGEDATARGGTLQVEAEGDSNIHVGEDAKAEGGTLEVEAHG DSNIHVGEHAKANGGRLQVKAQGDSNIHVGENAQAKGGQLIVQAQGDSNI HVGEDAKADGGELRVEAEWLEGGGSEQKLISEEDL |
| Z0-fiber H2E | MYDVPDYALQASGGGGSGGGGSGGGGSASHMSEIHVGENADANGGELKV TAKGDSEIHVGEDATARGGTLQVEAEGDSEIHVGEDAKAEGGTLEVEAHGD SEIHVGEHAKANGGRLQVKAQGDSEIHVGENAQAKGGQLIVQAQGDSEIH VGEDAKADGGELRVEAEWLEGGGSEQKLISEEDL |
| Z0-fiber H4N | MYDVPDYALQASGGGGSGGGGSGGGGSASHMSHINVGENADANGGELKV TAKGDSHINVGEDATARGGTLQVEAEGDSHINVGEDAKAEGGTLEVEAHG DSHINVGEHAKANGGRLQVKAQGDSHINVGENAQAKGGQLIVQAQGDSHI NVGEDAKADGGELRVEAEWLEGGGSEQKLISEEDL |

| | |
|---|---|
| Z0-fiber H4E | MYDVPDYALQASGGGGSGGGGSGGGGSASHMSHIEVGENADANGGELKVTAKGDSHIEVGEDATARGGTLQVEAEGDSHIEVGEDAKAEGGTLEVEAHGDSHIEVGEHAKANGGRLQVKAQGDSHIEVGENAQAKGGQLIVQAQGDSHIEVGEDAKADGGELRVEAEWLEGGGSEQKLISEEDL |
| Z3 H22E | MYDVPDYALQASGGGGSGGGGSGGGGSASHMSEEVNERVKQLAEKAKEATDKEEVRKIVQELAELAQCSTDSELVNEIVKQLAEVAKEATDKELVRKIVQILAELAQCSTDSELVNEIVKQLEEVAKEATDKELVRKIEQILEELKQCSTDLEGGGSEQKLISEEDL |
| Z3 E33Q | MYDVPDYALQASGGGGSGGGGSGGGGSASHMSEEVNERVKQLAEKAKEATDKHEVRKIVQELAQLAQCSTDSELVNEIVKQLAEVAKEATDKHLVRKIVQILAQLAQCSTDSELVNEIVKQLEEVAKEATDKHLVRKIEQILEQLKQCSTDLEGGGSEQKLISEEDL |

**Table S6. Amino acid sequences of enriched Hematite binders and design mutants as displayed on yeast.** Full sequences include linkers and tags.

| Name | Sequence |
|---|---|
| H1 | MYDVPDYALQASGGGGSGGGGSGGGGSASGGEESKKGTSQKVITRTSEVVVIVSVVVDTNAQLDIRAEGEVAILHIHAHGTNAELNISASGTFAHLCIVACGTNAHLNIRAEGTFACLIIYAEGDHAELNISASGDYACLCIIACGKSAVLRIEARGKHAILIIHAEGEDVQIDTKSQGECTDNQERKHGDGSWGGGSEQKLISEEDL |
| H2 | MYDVPDYALQASGGGGSGGGGSGGGGSASGGEESKKGTAQLVITATSEVAVLVIVAVDTNAQLDIRAEGEVAILHIHAHGTNAELNISASGTFAHLCIVACGTNAHLNIRAEGTFACLIIYAEGDHAELNISASGDYACLCIIACGKSAVLRIEARGKHAILIIHAEGEDAQLDIKAQGECADLQIRAHGGSWGGGSEQKLISEEDL |
| H3 | MYDVPDYALQASGGGGSGGGGSGGGGSASGGEESKKGSTSQKVITRTSEVVVIVSVVVDTNAQLDIRAEGEVAILHIHAHGTNAELNISASGTFAHLCNCCLWYKRAP |
| H4 | MYDVPDYALQASGGGGSGGGGSGGGGSASGGEESKKGSTSQKVITRTSEVVVIHSTVCDTNAQLDIRAEGEVAVLHITACGTNAELNISASGMLLFFISLPVALMHI |
| H5 | MYDVPDYALQASGGGGSGGGGSGGGGSASGGEESKKGSTIEKKIERTDDDCIVVVIVSHDDDEIRVEIKSDGDSCIVVVIVSHGDSQIEVSITSNGDIIVSHGDSQIVVQIQSHGDSCIGDSCIVVVIVSHGDSEIVVQITSEGDSCIVVVIVSHGDSTIEVQIQSHGDSCIVVVIVSHGNGDRKVERKSEGSWGGGSEQKLISEEDL |
| H6 | MYDVPDYALQASGGGGSGGGGSGGGGSASGGEESKKGTIEKKIERTDDDCIVVVIVSHDDDEIRVEIKSDGDSCIVVVIVSHGDSQIEVSITSNGDSCIVVVIVSHGDSQIVVQIQSHGDSCIVVVIVSHGDSEIVVQITSEGDSCIVVVIVSHGDSTIEVQIQSHGDSCIVVVIVSHGNGDRKVERKSEGNSGSWGGGSEQKLISEEDL |
| H7 | MYDVPDYALQASGGGGSGGGGSGGGGSASGPKKIVKDAKEKLEKLLEDAKDGGEHLAAEIAEELAHEAEHALKELLDEGASPELIVDLAETALRALLEIAKDGGEHLAARIAEILAELAHHALHVLLHDGASPKLIEDLAKTALDALEEIARDGGEDLAKHIDHILRDLEHSARDVLRDDGASGSWGGGSEQKLISEEDL |
| H8 | MYDVPDYALQASGGGGSGGGGSGGGGSASGPKKIVKDAKEKLEKLLEDAKDGGEHLALRIAHELAAEAKYALHELLKEGASPELIVDLAETALRALLEIAKDGGEHLALRIAHILAALAKYALHVLLKDGASPKLIEDLAKTALDALEEIARDGGEHLALRIDHILRALEKYARHVLRKDGASGSWGGGSEQKLISEEDL |
| H9 | MYDVPDYALQASGGGGSGGGGSGGGGSASSPKKIVKDAKEKLEKLLEDAKDGGEHLALRIAHELAAEVKYALHELLKEGASPELIVDLAETALRALLEIAKDGGEHLALRIAHILAALAKYALHVLLKDGASPKLIEDLAKTALDALEEIARDGGEHLALRIDHILRALEKYARHVLRKD |

| | |
|---|---|
| H10 | MYDVPDYALQASGGGGSGGGGSGGGGSASSSEEIVEEAETALKALLEEAEK<br>GGKDDAEDIAEKLAELAEDALEVLEDAGASPELIVRLAETALKALLAIAELG<br>GESLARSIAKILAKLAAKALQVLRKLGASPELIKKLAETAEEALEAIAKLGG<br>ESLARSIAKILAKLAAKAKQVQRKLGASGSWGGGSEQKLISEEDL |
| H11 | MYDVPDYALQASGGGGSGGGGSGGGGSASSSEEIVEEAETALKALLEEAEK<br>GGKHDALHIASGSSGGRDESTVITTEIEKSSEEIVEEAETALKALLEEAEKGG<br>KHDALHIAKKLAKLAAYALRVLRHSGASPELIVRLAETALKALLAIAELGGE<br>ELAEHIAEILAKLAKCAKHVQLCLGASGSWGGGSEQKLISEEDL |

**Table S7. Amino acid sequences of enriched ZnO binders, design mutants, derivative designs, and assemblies.** Full sequences including linkers and tags of the proteins expressed in *E. coli.* Proteins were expressed with 6xHis tags unless otherwise indicated with (strep) note indicating a Strep-tag was used instead. Strep-tagged proteins purified with Strep-Tactin beads (see Methods) were used to assay ion binding and mineral nucleation to eliminate potential effects of His-tags.

| Name | Sequence |
|---|---|
| Z0 | MSGDSNGSESKITRKGDRHIHVEEDATARGGTLQVEAEGDSHIHVGEDAKAEGGTLEVEAHGDSHIHVGEHAKANGGRLQVKAQGDSHIHVGENAQAKGGQLIVQAQGDSHIHVGEDAKSDGGEIRVESEGDSHIHEGENVGW |
| Z0-fiber | MSHIHEGENVDSNGSESKITRKGDRHIHVEEDATARGGTLQVEAEGDSHIHVGEDAKAEGGTLEVEAHGDSHIHVGEHAKANGGRLQVKAQGDSHIHVGENAQAKGGQLIVQAQGDSHIHVGEDAKSDGGEIRVESEW |
| Z1 | MSGGEVEFDHEISEEECSHELGRANIKDVTFQADISVESIVKILGLANIKSVHFEHEISVEDCAKILGLANIKEVHFEEDISEEDERKTKGLANIGSWGGSGSHHWGSTHHHHHH |
| Z2 | MSGSELVNEIVKQLAEVAKEATDKHLVRKIVQILAELAQCSTDSELVNEIVKQLAEVAKEATDKHLVRKIVQILAELAQCSTDSELVNEIVKQLAEVAKEATDKHLVRKIVQILAELAQCSTDGSWGGSGSHHWGSTHHHHHH |
| Z3 | MSGSEEVNERVKQLAEKAKEATDKHEVRKIVQELAELAQCSTDSELVNEIVKQLAEVAKEATDKHLVRKIVQILAELAQCSTDSELVNEIVKQLEEVAKEATDKHLVRKIEQILEELKQCSTDGSWGGSGSHHWGSTHHHHHH |
| Z4 | MSGGDSTEESKRISRISTEARTSGTEESLRQAIEDVAQLAKKSQDSTVLSKAISVISTIARTSGSEEALRQAIRAVAEIAKEAQDPTVLSKAISVISTIARTSGSEEARRQAERAEEEIRRRAQGSWGGSGSHHWGSTHHHHHH |
| Z5 | MSGRTPLLKLIRIVRTGSEEQQKSALEQLKRLLEAGADPNAADNHGRTPLLKLIRIVRTGSEEQQKAALELLKLLLEAGADPNAADNHGRTPQQKLKRIERTGSEEQQKAAKELLKLLEEAGADPNAADNHGSWGGSGSHHWGSTHHHHHH |
| Z6 | MSGGDSNGSESKITRKGRRVIHVEHDATARGGTLQVEAEGLSAIFVGLDAKAEGGTLEVEAHGPSTIQVGAHAKANGGRLQVKAQGHSRIHVGQNAQAKGGQLIVQAQGKSYIYVGRDAKSDGGEIRVESEGKSHIYEGSNVGSWGGSGSHHWGSTHHHHHH |
| Z7 | MSGGDRHSEINSSSDAYAEEMIDSEDNARATLVINSDGDSEAELVIFSDGDSQAKLEINSKGDSIADLEIESLGDSSATLRIHSKGESEAYLVIISDGDSEAQLQIVSDGDSDAVLEIDSEGSSSAQLQIISKGESDAFSDIESASSKSVQIEQDASGSKGSWGGSGSHHWGSTHHHHHH |
| Z8 | MSGGKRDDHVTETTGSKVDDGVEINAWTTITTGTEINAGTVINERTTITTGTIINEGTEINEGTTITTGTEINAGTKINAGTTITTGTRINAGTEINE |

| | |
|---|---|
| | GTTITTHTTINRGTHINAGTTITTHTEINEGTTINSGTTITTHTKIDSGS KIDSGSTITTGRGSWGGSGSHHWGSTHHHHHH |
| Z9 | MSGGGEESKKGTIQKERQGPTSTITVTSTSDDTQIEVKAEGPKTTITV TATGNKTRINVVAHGPTTTITVTATGNQTSINVEAEGDTTTITVTATGD DTSIRVVARGDTTTITVTATGDNTSIEVEAEGNNTTITVTATGNNTQISI TSRGDNTTITKTSTGSNRGSWGGSGSHHWGSTHHHHHH |
| Z10 | MSGGGEESKKGNTTIQVEAQGPTTTITVTATGDDTQIEVKAEGPKTTI TVTATGNKTRINVVAHGPTTTITVTATGNQTSINVEAEGDTTTITVTAT GDDTSIRVVARGDTTTITVTATGDNTSIEVEAEGNNTTITVTATGNNT QISITARGDNTTITVTATGGSWGGSGSHHWGSTHHHHHH |
| Z11 | MSGGGEESKKGDVERDVKRESSTEVIIVVVSESETRVSITVQASADG EVIIVVVAEAEGEVSITVQAEAKGEVIIVVVAEAEGQVHISVQARADG EVIIVVVAEAEGEVHIQVEAEAKGEVIIVVVAEAEGQVVISVSAKAEG EVIIVIVSEAEGESSVSESKSSNGGSWGGSGSHHWGSTHHHHHH |
| Z12 | MSGGGEESKKGDVERDVKRESSTTVYITVTSTSTTRVSITVQASADG TVYITVTATATGEVSITVQAEAKGTVYITVTATATGQVHISVQARADG TVYITVTATATGEVHIQVEAEAKGTVYITVTATATGQVVISVSAKAEG TVYITITSTATGESSVSESKSSNGGSWGGSGSHHWGSTHHHHHH |
| Z13 | MSGGGEESKKGDKTVVINSTKDDEIVVVIKSDKDEKVQVEIQSEGA DEVVVEIESTGAKQVTVQIRSSGADEVVVQIRSHGAEQVIVEIESTGA DEVVVQIQSTGAQQVQVDIKSTGADEVVVQIQSTGAKEVEVSIKSEG ADESVVEIDSKGAERITVSRDDKGSEGSWGGSGSHHWGSTHHHHH H |
| Z14 | MSGGGEESKKGGYSCRAVGVDGRAVQDTQGTCTAQAAGAGAMAS GTSEPGSASTAQAAGRGATARSTSTGRGAATTQAAGTASATSNAIGQ GAATTQAAGSAGGRATGSATTSSSASQPAQTQQIAGPGFQTAKSFAR NAATTQVAASHGSWGGSGSHHWGSTHHHHHH |
| Z0-fiber (strep) | MSGHIHEGENVDSNGSESKITRKGDRHIHVEEDATARGGTLQVEAEG DSHIHVGEDAKAEGGTLEVEAHGDSHIHVGEHAKANGGRLQVKAQ GDSHIHVGENAQAKGGQLIVQAQGDSHIHVGEDAKSDGGEIRVESE WSAWSHPQFEK |
| Z0-fiber H2E (strep) | MSGSEIHVGENADANGGELKVTAKGDSEIHVGEDATARGGTLQVEA EGDSEIHVGEDAKAEGGTLEVEAHGDSEIHVGEHAKANGGRLQVK AQGDSEIHVGENAQAKGGQLIVQAQGDSEIHVGEDAKADGGELRVE AEWGSGWSAWSHPQFEK |
| Z0-fiber H4E (strep) | MSGSHIEVGENADANGGELKVTAKGDSHIEVGEDATARGGTLQVEA EGDSHIEVGEDAKAEGGTLEVEAHGDSHIEVGEHAKANGGRLQVK AQGDSHIEVGENAQAKGGQLIVQAQGDSHIEVGEDAKADGGELRVE AEWGSGWSAWSHPQFEK |
| Z0-fiber H2E and H4E (strep) | MSGSEIEVGENADANGGELKVTAKGDSEIEVGEDATARGGTLQVEA EGDSEIEVGEDAKAEGGTLEVEAHGDSEIEVGEHAKANGGRLQVKA QGDSEIEVGENAQAKGGQLIVQAQGDSEIEVGEDAKADGGELRVEA |

| | |
|---|---|
| | EWGSGWSAWSHPQFEK |
| Z3 (strep) | MSGSEEVNERVKQLAEKAKEATDKHEVRKIVQELAELAQCSTDSEL VNEIVKQLAEVAKEATDKHLVRKIVQILAELAQCSTDSELVNEIVKQL EEVAKEATDKHLVRKIEQILEELKQCSTDGSWSAWSHPQFEK |
| Z3 H22E (strep) | MSGSEEVNERVKQLAEKAKEATDKEEVRKIVQELAELAQCSTDSEL VNEIVKQLAEVAKEATDKELVRKIVQILAELAQCSTDSELVNEIVKQL EEVAKEATDKELVRKIEQILEELKQCSTDGSGWSAWSHPQFEK |
| Z3 R25Q (strep) | MSGSEEVNERVKQLAEKAKEATDKHEVQQIVQELAELAQCSTDSEL VNEIVKQLAEVAKEATDKHLVQQIVQILAELAQCSTDSELVNEIVKQL EEVAKEATDKHLVQQIEQILEELKQCSTDGSGWSAWSHPQFEK |
| Z3 E33Q (strep) | MSGSEEVNERVKQLAEKAKEATDKHEVRKIVQELAQLAQCSTDSEL VNEIVKQLAEVAKEATDKHLVRKIVQILAQLAQCSTDSELVNEIVKQ LEEVAKEATDKHLVRKIEQILEQLKQCSTDGSGWSAWSHPQFEK |
| Z3 C37A (strep) | MSGSEEVNERVKQLAEKAKEATDKHEVRKIVQELAELAQASTDSEL VNEIVKQLAEVAKEATDKHLVRKIVQILAELAQASTDSELVNEIVKQL EEVAKEATDKHLVRKIEQILEELKQASTDGSGWSAWSHPQFEK |
| Z4 (strep) | MSGDSTEESKRISRISTEARTSGTEESLRQAIEDVAQLAKKSQDSTVLS KAISVISTIARTSGSEEALRQAIRAVAEIAKEAQDPTVLSKAISVISTIAR TSGSEEARRQAERAEEEIRRRAQGSWSAWSHPQFEK |
| zab1 | MSGDLHVHIGEEEAEPEEAEEKLTAAKEEIVASGEPVDLHLHVGERV SPEVVETLEEELEELLEEADVTDAHLHIGEEVEEPEEVRERLLAVFEE AGVDAHVHVGEGSSGSSGSGSHHWGSTHHHHHH |
| zab2 | MSGDLHLHVGEEATPELLAEAKELLEEALEEAAERGEKLDVHLHIGE RVLASPEDVLDLILLIAEAGADAHVHIGESFPLSSEELREEVEKRLLE KTGKPLPDVHIHVGEGSSGSSGSGSHHWGSTHHHHHH |
| zab3 | MSGDLHVHLGEEVELTEEVAERIAEAFVAAEGGDLHLHIGERVTAEP EEAEELIEELLEEIKARVKEPVDLHLHVGEEVEPHIRELFEELVEELKE AGILRDAHVHIGEGSSGSSGSGSHHWGSTHHHHHH |
| zab4 | MSGDFHFHVGEVATVELVEEAREELLEAIEAAKDAERLDLHLHIGEK VLPEAKPVVLEALEDFLAKVGETGVTLDAHIHIGENAPELKPAVEALI EEARAKYGKVGDVHVHVGEGSSGSSGSGSHHWGSTHHHHHH |
| zab5 | MSGDVHVHVGERATTRDVAETLRELLEAGVLLADLHLHVGEEFEW EEEELPALLEAFRAAFEALAEAGLPADVHLHLGENADPERAEALLEE LKALAEEAGVELRDFHVHVGEGSSGSSGSGSHHWGSTHHHHHH |
| zab6 | MSGDIHIHVGERATPEEVREALEEAEEEAEESGEPVDLHLHVGEEVE DPEAVTALLEEFIKKVDLSDLHLHLGESFAPRAEEVRARLEAAVEEAR ALGKTVEDFHLHIGEGSSGSSGSGSHHWGSTHHHHHH |
| zab7 | MSGDIHIHVGEEVKEEVEPLIEELEELAEEAEEKGVEIDAHVHIGEKP ELEVAKEALEVLKEYIEELIEEGKSVDLHVHLGERPTLEDVKELAKE LEELFEEGVTDAHLHFGEGSSGSSGSGSHHWGSTHHHHHH |
| zab8 | MSGDVHLHVGENVPEEKREELAEEVEAKVEELGEIDLHLHLGEKVE |

| | |
|---|---|
| | VDYETLLELVLRLVETVAEKGVTDAHIHVGEVATRELLERALAEVEE HPGGDLHLHVGEGSSGSSGSGSHHWGSTHHHHHH |
| zab9 | MSGDTHVHVGEAASPEAVEALREEAAKLGDAHVHLGEEATDEHVE VWLEAVREAVELAEAAGRRVDTHLHVGEAPEPEEVPPAAEEFLKAF AEALEAGEDFHYHAGEGSSGSSGSGSHHWGSTHHHHHH |
| zab10 | MSGDVHVHVGEGNEELEKELRERFEEAAAKGAADLHLHVGEKVDP SLAPAVRALAELALELAELGVDLHLHVGEGTLEEILEAQEKLKEILEE AEEEGPLDVHLHVGEGSSGSSGSGSHHWGSTHHHHHH |
| zab11 | MSGDTHVHLGENAPPEWEEELLAVAEEAGVDIHLHVGEEVDPRTLP VLKEIIKLLGEDGSDAHIHIGEKGTLELAEEALEKVKELLAAKPGEDF HLHVGEGSSGSSGSGSHHWGSTHHHHHH |
| Z4-C3i | MSGDSTVLSKAISVISTIARTSGSEEALRQAIEAVAEIAKEAQDPTVLS KALEAITKILFTSIDNEEVARQAREAVLELSQDEETRELLEKLREAED EEEKREIIEELAKRGPEAILALLAEAIILGLDVEEVLKIAIKINSKDSDA ASLLITAISELARQKGTEESLRQAIEDVAQLAKESQDSTVLSKAISVIS TIARTSGSEEALRQAIEAVAEIAKEAQGSSGSSGSGSHHWGSTHHHH HH |
| Z4-C3ii | MSGDSTVLSKAISVISTIARTSGSEEALRQAIEAVAEIAKEAQDSTVLS KAAEALAALAAEALRIGNEEALRQAIEALVEIAKELGLEEFAKLLKE LGERLEKLLREGAGIEAFWELIREFAKKAKGLDSTSLSVVIALIGAFV RTFADEITEESLRQAIEDVAQLAKESQDSTVLSKAISVISTIARTSGSEE ALRQAIEAVAEIAKEAQGSSGSSGSGSHHWGSTHHHHHH |
| Z4-C3ii-XL | MSGDSTVLSKAISVISTIARTSGSEEALRQAIEAVAEIAKEAQDSTVLS KAISVISTIARTSGSEEALRQAIEAVAEIAKEAQDSTVLSKAAEALAAL AAEALRIGNEEALRQAIEALVEIAKELGLEEFAKLLKELGERLEKLLR EGAGIEAFWELIREFAKKAKGLDSTSLSVVIALIGAFVRTFADEITEES LRQAIEDVAQLAKESQDSTVLSKAISVISTIARTSGSEEALRQAIEAVA EIAKEAQDSTVLSKAISVISTIARTSGSEEALRQAIEAVAEIAKEAQGSS GSSGSGSHHWGSTHHHHHH |
| Z4-C6 | MSGDSTVLSKAISVISTIARTSGSEEALRQAIEAVAEIAKEAQLTPEVL KAVIDAYVTIVEAAVKLGREYAEKVKEEILKKLKELPEVTPEDLASAE IDIISTIVEEDSSLTEESLRQAIEDVAQLAKESQDSTVLSKAISVISTIAR TSGSEEALRQAIEAVAEIAKEAQGSSGSSGSGSHHWGSTHHHHHH |
| Z4-C3i-KO | MSGDSEVLSKAIQVISEIARDSGSEEALRQAIEAVAEIAKEAQDPTVLS KALEAITKILFTSIDNEEVARQAREAVLELSQDEETRELLEKLREAED EEEKREIIEELAKRGPEAILALLAEAIILGLDVEEVLKIAIKINSKDSDA ASLLITAISELARQKGTEESLRQAIEDVAQLAKESQDSEVLSKAIQVIS EIARDSGSEEALRQAIEAVAEIAKEAQGSSGSSGSGSHHWGSTHHHH HH |
| Z4-C3ii-KO | MSGDSEVLSKAIQVISEIARDSGSEEALRQAIEAVAEIAKEAQDSTVLS KAAEALAALAAEALRIGNEEALRQAIEALVEIAKELGLEEFAKLLKE LGERLEKLLREGAGIEAFWELIREFAKKAKGLDSTSLSVVIALIGAFV RTFADEITEESLRQAIEDVAQLAKESQDSEVLSKAIQVISEIARDSGSE |

| | |
|---|---|
| | EALRQAIEAVAEIAKEAQGSSGSSGSGSHHWGSTHHHHHH |
| Z4-C6-KO | MSGDSEVLSKAIQVISEIARDSGSEEALRQAIEAVAEIAKEAQLTPEVLKAVIDAYVTIVEAAVKLGREYAEKVKEEILKKLKELPEVTPEDLASAEIDIISTIVEEDSSLTEESLRQAIEDVAQLAKESQDSEVLSKAIQVISEIARDSGSEEALRQAIEAVAEIAKEAQGSSGSSGSGSHHWGSTHHHHHH |
| Z4-cage (strep) | MSKEGSAWSHPQFEKGGSGSDSTVLSKAISVISTIARTSGSEEALRQAIEAVAEIAKEAQDPTVLSKALEAITKILFTSIDNEEVARQAREAVLELSQDEETRELLEKLREAEDEEEKREIIEELAKRGPEAILALLAEAIILGLDVEEVLDIAIEINKKDSDAASLLITAISELARQKGTEESLERAIRRIALLAIEAKDSTVLSKAISVISTIARTSGSEEALRLAIEAVALIALGGEDTVPARYIKISFEAELAGLDTKRLLEEGPKLYPELSIPDLMAIALFTHLHLDPYFLYRLLQQSR |
| Z4-tube (strep) | MGSAWSHPQFEKGSSGSSVEEDDDDIMRRALTTTIVIENGILRKDFTKKVVDIIWRKSDEIRDPEVAKELIEELKKLLKEEGANIKETIAILNLIKDIAIASSDVEVHKYAIDTVAEIAKEAQNQTVFSEAIQVIKEIARTSGNQEAHKIAIDAVAEIAKESKNQTVLSEAISVISTIARTSGNQEALRQAIEAVAEIAKESKNQTVLSKAISVISTIARTSGNQEALRQAIEAVAEIAKESKNQTVLSKAISVISTIARTSGEQEALRQAIEAVAEIAKESKEQAVLSKAISVISTIARTSGEKEALRQAIEAVAEIAKESKEQAVLSKAISVISTIARTSGEKEALRQAIEAVAEIAKESKEQAVLSKAISVISTIARTSGEKEALQQAIWAIAEIAKESKEKAVLSKAISEISTIALTSGEKEVLQQAIWAIKEIALESKDVNVQVKAIYLIVKIAEASGSEEVKELAKEKIKEIAKESKDKRVKLAAILALVLMKQLEELEEEATELFEKLEEEGEKVMDEILEKISEKVEKFVNLSTALELLA |
| zno4_oligo1_C3 | MSGDSTVLSKAISVISTIARTSGSEEALRQAIEAVAEIAKEAQDSTVLSKALIAIAKILLTAIDNKEVARQALQAVLELSQNQETAELLARLEAATDEAERNAIIEELAKLGPSSVLALLARAIILGLDLEEVVKTLIKINKEDSDSSSLLIEIISELARLKGTEESLRQAIEDVAQLAKESQDSTVLSKAISVISTIARTSGSEEALRQAIEAVAEIAKEAQGSSGSSGSGSHHWGSTHHHHHH |
| zno4_oligo2_C3 (renamed Z4-C3i; relisted here for clarity) | MSGDSTVLSKAISVISTIARTSGSEEALRQAIEAVAEIAKEAQDPTVLSKALEAITKILFTSIDNEEVARQAREAVLELSQDEETRELLEKLREAEDEEEKREIIEELAKRGPEAILALLAEAIILGLDVEEVLKIAIKINSKDSDAASLLITAISELARQKGTEESLRQAIEDVAQLAKESQDSTVLSKAISVISTIARTSGSEEALRQAIEAVAEIAKEAQGSSGSSGSGSHHWGSTHHHHHH |
| zno4_oligo3_C3 | MSGDSTVLSKAISVISTIARTSGSEEALRQAIEAVAEIAKEAQDSTVLSKAIEALAELAAEAIKNGNEEALRQALKAAIEIAKELGLEEFAKLLEELEKRLTELLKAGKGIKEFWEVIKEFAKKSLKLDSTSLSYVISLISAFARTNAEEITEESLRQAIEDVAQLAKESQDSTVLSKAISVISTIARTSGSEEALRQAIEAVAEIAKEAQGSSGSSGSGSHHWGSTHHHHHH |
| zno4_oligo4_C3 (renamed Z4-C3ii; | MSGDSTVLSKAISVISTIARTSGSEEALRQAIEAVAEIAKEAQDSTVLSKAAEALAALAAEALRIGNEEALRQAIEALVEIAKELGLEEFAKLLKE |

| | |
|---|---|
| relisted here for clarity) | LGERLEKLLREGAGIEAFWELIREFAKKAKGLDSTSLSVVIALIGAFVRTFADEITEESLRQAIEDVAQLAKESQDSTVLSKAISVISTIARTSGSEEALRQAIEAVAEIAKEAQGSSGSSGSGSHHWGSTHHHHHH |
| zno4_oligo5_C3 | MSGDSTVLSKAISVISTIARTSGSEEALRQAIEAVAEIAKEAQNKVLSKALEALTKILVEAIRRGLNGLEIAKLIAEKIKEVVPDWEVVVIDTADEEQFAAQWAEAKALIAAGKRVYVVVVFDTEEELRKAGELLAQLAKESQDSTLVLRVIAFYEAAAPLLTEESLRQAIEDVAQLAKESQDSTVLSKAISVISTIARTSGSEEALRQAIEAVAEIAKEAQGSSGSSGSGSHHWGSTHHHHHH |
| zno4_oligo6_C3 | MSGDSTVLSKAISVISTIARTSGSEEALRQAIEAVAEIAKEAQDSTVLSKAISVISTIARTSGSEEALRQAIEAVAEIAKEAQGDLSFKIKALSEVAGAILSLESLSDEEKRALLNELADALGLSEKIKKLVNELLDFAFRAEELTEEEVRERLERIRKLFLEAGITDSEVKSELLSLLSEIVVESLEAGRLTEEHARLIIEAVAEIAKEEFKDSTVLSKAISVISTIARTSGSEEALRQAIEAVAEIAKEAQDSTVLSKAISVISTIARTSGSEEALRQAIEAVAEIAKEAQGSSGSSGSGSHHWGSTHHHHHH |
| zno4_oligo7_C3 | MSGDSTVLSKAISVISTIARTSGSEEALRQAIEAVAEIAKEAQDSTVLSKAISVISTIARTSGSEEALRQAIEAVAEIAKEAQDSTVLDKAIAAILELAAIATARGWEEAARQAAEALLEIAKDAGYSEKILEALKKILQLLLDIIENKPSDAELQEKYLQIAKLLVEAIKLRKEEGLDASAALSAFSKLIGTLLELEPRITEETARQIIELVAEAAKEIGDSTVLSKAISVISTIARTSGSEEALRQAIEAVAEIAKEAQDSTVLSKAISVISTIARTSGSEEALRQAIEAVAEIAKEAQGSSGSSGSGSHHWGSTHHHHHH |
| zno4_oligo8_C3 | MSGDSTVLSKAISVISTIARTSGSEEALRQAIEAVAEIAKEAQDSTVLSKAISVISTIARTSGSEEALRQAIEAVAEIAKEAQGETRLEATLKLISTVLSLIAQRKDLSKETIEEFVKKALELVKETAKKTGKEEEAKALKEIFLLLLELLEDYRSGKLDEEKLKKYLKKLKELIKKLTSTQLSKVLSTISTVVRNSGFSEEFLRQIIEFVAEIAKEAGDSTVLSKAISVISTIARTSGSEEALRQAIEAVAEIAKEAQDSTVLSKAISVISTIARTSGSEEALRQAIEAVAEIAKEAQGSSGSSGSGSHHWGSTHHHHHH |
| zno4_oligo9_C4 | MSGDSTVLSKAISVISTIARTSGSEEALRQAIEAVAEIAKEAQDSTVLSKAISVISTIARTSGSEEALRQAIEAVAEIAKEAQDSTVLSKALSAIFWIASTLIRQGFDVSEALKAVEEFKEHPLAPVVRQAIEILEAIRAGAEEEEIREKIKELAKEVEKIKSLSALSLVSEVASELARAGREEAIRQIIEAVAEVAKKIKDSTVLSKAISVISTIARTSGSEEALRQAIEAVAEIAKEAQDSTVLSKAISVISTIARTSGSEEALRQAIEAVAEIAKEAQGSSGSSGSGSHHWGSTHHHHHH |
| zno4_oligo10_C4 | MSGDSTVLSKAISVISTIARTSGSEEALRQAIEAVAEIAKEAQDSTVLSKAISVISTIARTSGSEEALRQAIEAVAEIAKEAQDSTVLSKAESARTELEFELLLVRGASEEELREFLEKQLKKAEEALKKGDKTKLSKALSVISTFLRKSGASEEALREAIEKVAELAKESGDSTVLSKAISVISTIARTSGSEEALRQAIEAVAEIAKEAQDSTVLSKAISVISTIARTSGSEEALRQAIEAVAEIAKEAQGSSGSSGSGSHHWGSTHHHHHH |

| | |
|---|---|
| zno4_oligo11_C4 | MSGDSTVLSKAISVISTIARTSGSEEALRQAIEAVAEIAKEAQDETVLSKAISAISKILAEAVKNGYEEVAKYAEEVKKEILKYGKEELGLSEQKIKNIEALAVIDYYETLAPYLTEESLRQAIEDVAQLAKESQDSTVLSKAISVISTIARTSGSEEALRQAIEAVAEIAKEAQGSSGSSGSGSHHWGSTHHHHHH |
| zno4_oligo12_C6 (renamed Z4-C6; relisted here for clarity) | MSGDSTVLSKAISVISTIARTSGSEEALRQAIEAVAEIAKEAQLTPEVLKAVIDAYVTIVEAAVKLGREYAEKVKEEILKKLKELPEVTPEDLASAEIDIISTIVEEDSSLTEESLRQAIEDVAQLAKESQDSTVLSKAISVISTIARTSGSEEALRQAIEAVAEIAKEAQGSSGSSGSGSHHWGSTHHHHHH |
| zno4_oligo13_C6 | MSGDSTVLSKAISVISTIARTSGSEEALRQAIEAVAEIAKEAQLNSEVLKAVIDAAVTIVAAAVELGRETAERIKKEILEALKSLPSVTSSDLASAEISIISTIVEKDESLTEESLRQAIEDVAQLAKESQDSTVLSKAISVISTIARTSGSEEALRQAIEAVAEIAKEAQGSSGSSGSGSHHWGSTHHHHHH |
| zno4_oligo14_C6 | MSGDSTVLSKAISVISTIARTSGSEEALRQAIEAVAEIAKEAQDSTVLSKAISALSDIIVAAVRAGHPQSVVDEFEEYVDKVIDELIKQLKNEEDKKKLKNAYVLAKIDRKAAIAPYLTEESLRQAIEDVAQLAKESQDSTVLSKAISVISTIARTSGSEEALRQAIEAVAEIAKEAQGSSGSSGSGSHHWGSTHHHHHH |
| zno4_oligo15_C6 | MSGDSTVLSKAISVISTIARTSGSEEALRQAIEAVAEIAKEAQDLTVLSKAISALNTIIVAAVRAGLPQEVVEEFQKYVDETVKELIEELKEEEDKKALKNAYVIANITQKEAIAPYLTEESLRQAIEDVAQLAKESQDSTVLSKAISVISTIARTSGSEEALRQAIEAVAEIAKEAQGSSGSSGSGSHHWGSTHHHHHH |
| zno4_oligo16_C6 | MSGDSTVLSKAISVISTIARTSGSEEALRQAIEAVAEIAKEAQDLTVYSKAIEAQETIIVAAAENGLPQEFIDEARQKVAETVKELIAKEKSEENKKELENKYVIAYIDRLEAIAPLLTEESLRQAIEDVAQLAKESQDSTVLSKAISVISTIARTSGSEEALRQAIEAVAEIAKEAQGSSGSSGSGSHHWGSTHHHHHH |
| zno4_oligo17_C6 | MSGDSTVLSKAISVISTIARTSGSEEALRQAIEAVAEIAKEAQDSTVLSKAISVISTIARTSGSEEALRQAIEAVAEIAKEAQDSTVLSKALSVIVTTILETKDLKTAAKYLEYLAEEFKKQGLPEEVVTTVQVLEKILKGAPLEEIEKLIKNLSSTNLSKLLSLISTAAKNLPEIDALLAIEAVAEIAKEAGDSTVLSKAISVISTIARTSGSEEALRQAIEAVAEIAKEAQDSTVLSKAISVISTIARTSGSEEALRQAIEAVAEIAKEAQGSSGSSGSGSHHWGSTHHHHHH |
| zno4_oligo18_C6 | MSGDSTVLSKAISVISTIARTSGSEEALRQAIEAVAEIAKEAQDSTVLSKAISVISTIARTSGSEEALRQAIEAVAEIAKEAQDSTVLSKAISALLDIARLAQELGYEEAARQAIEAAEKLLEEIKKKGLWDSTVVSKEFSLLVLKNKLEGKSGEQTIEELAKYIKDKKLTDSTVLSKAISVISTIARTSGSEEALRQAIEAVAEIAKEAQDSTVLSKAISVISTIARTSGSEEALRQAIEAVAEIAKEAQGSSGSSGSGSHHWGSTHHHHHH |
| zno4_oligo19_C6 | MSGDSTVLSKAISVISTIARTSGSEEALRQAIEAVAEIAKEAQDSTVLS |

| | |
|---|---|
| | KAISVISTIARTSGSEEALRQAIEAVAEIAKEAQDDVKVKADLVIFELKIELLKLEGASDDEILEELIKELEELFKKYKDNVDVLIKIIEKASTIARKYLSPEKLPALIEKVAELAKEAKDSTVLSKAISVISTIARTSGSEEALRQAIEAVAEIAKEAQDSTVLSKAISVISTIARTSGSEEALRQAIEAVAEIAKEAQGSSGSSGSGSHHWGSTHHHHHH |
| zno4_oligo20_C6 | MSGDSTVLSKAISVISTIARTSGSEEALRQAIEAVAEIAKEAQDSTVLSKAISVISTIARTSGSEEALRQAIEAVAEIAKEAQDSTVLSKAESVKLELELELAKRKPLTPELIKEIIELLKKHYDNLLKLGADAATALSLISTAALTIGINEELARLLIELVAEIAKKEKDSTVLSKAISVISTIARTSGSEEALRQAIEAVAEIAKEAQDSTVLSKAISVISTIARTSGSEEALRQAIEAVAEIAKEAQGSSGSSGSGSHHWGSTHHHHHH |
| zno4_oligo21_C6 | MSGDSTVLSKAISVISTIARTSGSEEALRQAIEAVAEIAKEAQDSTVLSKAISVISTIARTSGSEEALRQAIEAVAEIAKEAQGEVKLKATFVQQILEFELLLLEGASEEELLKAFETALKTLLDEGASAEVIFKAIETFIELLREAGASEELLRQVIEVVAEKAKEIQDSTVLSKAISVISTIARTSGSEEALRQAIEAVAEIAKEAQDSTVLSKAISVISTIARTSGSEEALRQAIEAVAEIAKEAQGSSGSSGSGSHHWGSTHHHHHH |
| zno4_oligo22_C6 | MSGDSTVLSKAISVISTIARTSGSEEALRQAIEAVAEIAKEAQDSTVLSKAISVISTIARTSGSEEALRQAIEAVAEIAKEAQGEARSKALSVAASLVATLIKKVGGEAVKPLVEKLAKLAREAGDGAALAEIAVLELLIGNKEALKKLLADPRIDPLALISAAVTAAKTGADALQLIELVAEKAKEAQDSTVLSKAISVISTIARTSGSEEALRQAIEAVAEIAKEAQDSTVLSKAISVISTIARTSGSEEALRQAIEAVAEIAKEAQGSSGSSGSGSHHWGSTHHHHHH |
| zno4_oligo23_C6 | MSGDSTVLSKAISVISTIARTSGSEEALRQAIEAVAEIAKEAQDSTVLSKAISVISTIARTSGSEEALRQAIEAVAEIAKEAQDSTVLVKALSARAELAATAGDREIIPEILKAIKELAKAGADDTGLEKVGKALGKLLIAVGASVEEIIKIIKELAEINLSLAVAVAEGLTEPEVIEQLAKLLKEKGADSTVLSKAISVISTIARTSGSEEALRQAIEAVAEIAKEAQDSTVLSKAISVISTIARTSGSEEALRQAIEAVAEIAKEAQGSSGSSGSGSHHWGSTHHHHHH |

**Table S8. Amino acid sequences of enriched Hematite binders, design mutants, derivative designs.** Full sequences including linkers and tags of the proteins expressed in *E. coli.* Proteins were expressed with 6xHis tags unless otherwise indicated with (strep) note indicating a Strep-tag was used instead. Strep-tagged proteins purified with Strep-Tactin beads (see Methods) were used to assay ion binding and mineral nucleation to eliminate potential effects of His-tags.

| **Name** | **Sequence** |
|---|---|
| H1 | MSGGGEESKKGTSQKVITRTSEVVVIVSVVVDTNAQLDIRAEGEVAILHIHAHGTNAELNISASGTFAHLCIVACGTNAHLNIRAEGTFACLIIYAEGDHAELNISASGDYACLCIIACGKSAVLRIEARGKHAILIIHAEGEDVQIDTKSQGECTDNQERKHGDGSWGGSGSHHWGSTHHHHHH |
| H2 | MSGGGEESKKGTAQLVITATSEVAVLVIVAVDTNAQLDIRAEGEVAILHIHAHGTNAELNISASGTFAHLCIVACGTNAHLNIRAEGTFACLIIYAEGDHAELNISASGDYACLCIIACGKSAVLRIEARGKHAILIIHAEGEDAQLDIKAQGECADLQIRAHGGSWGGSGSHHWGSTHHHHHH |
| H3 | MSGGGEESKKGSTSQKVITRTSEVVVIVSVVVDTNAQLDIRAEGEVAILHIHAHGTNAELNISASGTFAHLCNCCLWYKRAPGSWGGSGSHHWGSTHHHHHH |
| H4 | MSGGGEESKKGSTSQKVITRTSEVVVIHSTVCDTNAQLDIRAEGEVAVLHITACGTNAELNISASGMLLFFISLPVALMHIGSWGGSGSHHWGSTHHHHHH |
| H5 | MSGGGEESKKGSTIEKKIERTDDDCIVVVIVSHDDDEIRVEIKSDGDSCIVVVIVSHGDSQIEVSITSNGDIIVSHGDSQIVVQIQSHGDSCIGDSCIVVVIVSHGDSEIVVQITSEGDSCIVVVIVSHGDSTIEVQIQSHGDSCIVVVIVSHGNGDRKVERKSEGSWGGSGSHHWGSTHHHHHH |
| H6 | MSGGGEESKKGTIEKKIERTDDDCIVVVIVSHDDDEIRVEIKSDGDSCIVVVIVSHGDSQIEVSITSNGDSCIVVVIVSHGDSQIVVQIQSHGDSCIVVVIVSHGDSEIVVQITSEGDSCIVVVIVSHGDSTIEVQIQSHGDSCIVVVIVSHGNGDRKVERKSEGNSGSWGGSGSHHWGSTHHHHHH |
| H7 | MSGGPKKIVKDAKEKLEKLLEDAKDGGEHLAAEIAEELAHEAEHALKELLDEGASPELIVDLAETALRALLEIAKDGGEHLAARIAEILAELAHHALHVLLHDGASPKLIEDLAKTALDALEEIARDGGEDLAKHIDHILRDLEHSARDVLRDDGASGSWGGSGSHHWGSTHHHHHH |
| H8 | MSGGPKKIVKDAKEKLEKLLEDAKDGGEHLALRIAHELAAEAKYALHELLKEGASPELIVDLAETALRALLEIAKDGGEHLALRIAHILAALAKYALHVLLKDGASPKLIEDLAKTALDALEEIARDGGEHLALRIDHILRALEKYARHVLRKDGASGSWGGSGSHHWGSTHHHHHH |
| H9 | MSGSPKKIVKDAKEKLEKLLEDAKDGGEHLALRIAHELAAEVKYALHELLKEGASPELIVDLAETALRALLEIAKDGGEHLALRIAHILAALAKYALHVLLKDGASPKLIEDLAKTALDALEEIARDGGEHLALRIDHILRALEKYARHVLRKDGSWGGSGSHHWGSTHHHHHH |
| H10 | MSGSSEEIVEEAETALKALLEEAEKGGKDDAEDIAEKLAELAEDALEVLEDAGASPELIVRLAETALKALLAIAELGGESLARSIAKILAKLAAKALQVLRKLGASPELIKKLAETAEEALEAIAKLGGESLARSIAKILAKLAAKAKQVQR |

| | |
|---|---|
| | KLGASGSWGGSGSHHWGSTHHHHHH |
| H11 | MSGSSEEIVEEAETALKALLEEAEKGGKHDALHIASGSSGGRDESTVITTEIEKSSEEIVEEAETALKALLEEAEKGGKHDALHIAKKLAKLAAYALRVLRHSGASPELIVRLAETALKALLAIAELGGEELAEHIAEILAKLAKCAKHVQLCLGASGSWGGSGSHHWGSTHHHHHH |
| H7-neg | MSPEKIVEDAKEKLEKLLEDAKDGGEHLAAEIAEELAHEAEHALKELLDEGASPELIVDLAETALRALLEIAKDGGEHLAARIAEILAELAHHALHVLLHDGASPKLIEDLAKTALDALEEIAEDGGEDLAKHIDHILEDLEHSAEDVLEDDGASGSWGGSGSHHWGSTHHHHHH |
| H9-neg | MSPKEIVKDAKEKLEKLLEDAKDGGEHLALRIAHELAAEVKYALHELLKEGASPELIVDLAETALRALLEIAKDGGEHLALRIAHILAALAKYALHVLLKDGASPKLIEDLAETALDALEEIARDGGEHLALRIDHILQALEKYAEHVLQKDGASGSWGGSGSHHWGSTHHHHHH |
| H7-neg (Strep) | MSPEKIVEDAKEKLEKLLEDAKDGGEHLAAEIAEELAHEAEHALKELLDEGASPELIVDLAETALRALLEIAKDGGEHLAARIAEILAELAHHALHVLLHDGASPKLIEDLAKTALDALEEIAEDGGEDLAKHIDHILEDLEHSAEDVLEDDGASGSGWSAWSHPQFEK |
| H9-neg (Strep) | MSPKEIVKDAKEKLEKLLEDAKDGGEHLALRIAHELAAEVKYALHELLKEGASPELIVDLAETALRALLEIAKDGGEHLALRIAHILAALAKYALHVLLKDGASPKLIEDLAETALDALEEIARDGGEHLALRIDHILQALEKYAEHVLQKDGASGSGWSAWSHPQFEK |

**Table S9. Amino acid sequences of previously reported ZnO-binding peptides used in this study.** Sequence of Small Ubiquitin-like Modifier (SUMO) construct used to aid expression and solubility of ZnO-binding peptides, and the four peptides fused to it. Peptides names are last name of first author in source publications.(*7–10*)

| | |
|---|---|
| SUMO-fusion | SHHHHHHGSWGSDSEVNQEAKPEVKPEVKPETHINLKVSDGSSEIFFKIKKTTPLRRLMEAFAKRQGKEMDSLRFLYDGIRIQADQTPEDLDMEDNDIIEAHREQIGSGSGS(Peptide) |
| Umetsu | EAHVMHKVAPRP |
| Tomczak | GLHVMHKVAPPRGGG |
| Wei | GAMHLPWHMGTL |
| Golec | TMGANLGLKWPV |

**Table S10. X-ray crystallography data collection and refinement statistics.** Highest-resolution shells are shown in parentheses.

| | Z4-C3i (PDB code: pdb_00009D92) | Z4-C3ii (PDB code: pdb_000037TT) |
|---|---|---|
| Resolution range | 34.75 - 3.06 (3.14 - 3.06) | 32.84 - 2.99 (3.15 - 2.99) |
| Space group | $P\ 2_1$ | $P\ 2_1$ |
| Unit cell | 86.66, 72.15, 119.52; 90.00, 109.78, 90.00 | 67.59, 68.52, 111.75; 90.00, 90.88, 90.00 |
| Unique reflections | 28082 (2140) | 19063 (3022) |
| Multiplicity | 5.0 (5.3) | 5.1 (5.5) |
| Completeness (%) | 98.16 (97.79) | 90.7 (99.8) |
| Mean I/sigma(I) | 3.10 (0.52) | 6.3 (1.1) |
| Wilson B-factor | 84.12 | 91.38 |
| R-merge | 0.2755 (3.187) | 0.113 (1.265) |
| R-pim | 0.1316 (1.488) | 0.060 (0.683) |
| $CC_{1/2}$ | 0.993 (0.343) | 0.993 (0.523) |
| Reflections used in refinement | 26014 (1992) | 18927 (1466) |
| R-work | 0.2400 (0.3788) | 0.2312 (0.3337) |
| R-free | 0.2987 (0.3984) | 0.2905 (0.3933) |
| Number of non-hydrogen atoms | 10006 | 9249 |
| macromolecules | 9996 | 9248 |
| water | 10 | 1 |
| Protein residues | 1320 | 1236 |
| RMS(bonds) | 0.002 | 0.003 |
| RMS(angles) | 0.35 | 0.50 |
| Ramachandran favored (%) | 97.55 | 96.48 |
| Ramachandran allowed (%) | 2.37 | 3.19 |
| Ramachandran outliers (%) | 0.08 | 0.33 |
| Average B-factor | 92 | 99 |
| macromolecules | 92 | 99 |
| water | 71 | 88 |

### Table S11. Cryo-EM data collection and processing statistics.

| Data Collection | | Z4-cage | |
|---|---|---|---|
| Microscope | | Glacios (Thermo) | |
| Voltage (kV) | | 200 | |
| Detector | | K3 (Gatan) | |
| Energy Filter | | N/A | |
| Recording mode | | Counting | |
| Magnification | | 45,000X | |
| Movie micrograph pixel size (Å) | | 0.885 | |
| Dose rate ($e^{-}$/Å$^2$/s) | | 8.4 | |
| No. of frames per movie micrograph | | 100 | |
| Frame exposure time (ms) | | 50 | |
| Movie micrograph exposure time (s) | | 10 | |
| Total dose ($e^{-}$/Å$^2$) | | 50 | |
| Under focus range (μm) | | 0.8-1.8 | |
| Total number of movies collected | | 4,018 | |
| **Map Processing** | **O Z4-Cage** | **C1 Z4-Cage** | **Z4-C3i (Local)** |
| Extraction Box Size (pix) | 520 | 520 | 520 |
| Fourier crop to Box Size (pix) | N/A | N/A | N/A |
| Initial particle images (no.) | 608,979 | 608,979 | 608,979 |
| Final particle images (no.) | 551,868 | 551,868 | 1,332,943 |
| Applied Symmetry | O (Relaxed) | C1 | C1 |
| Map resolution (Å) | 3.8 | 3.9 | 3.3 |
| FSC threshold | 0.143 | 0.143 | 0.143 |
| Map resolution range (Å) | 3.2-4.4 | 3.2-4.4 | 2.5-3.9 |

**Movie S1. Iron oxide growth reactions.** Video shows digital scans of iron oxide growth reactions (see Methods) over 120 minutes with one frame captured per minute. Video is at 30x speed. Rows contain six replicates for each condition containing from top to bottom: hematite nanoparticle (NP), magnetite NP, no additive, lysozyme, H7-neg, and H9-neg. Full reaction conditions provided in Methods and experimental setup show in **fig. S23.**

**Data S1. Raw data.**
Contains all plotted data in raw format.
